# Remodeling of extracellular matrix collagen IV by MIG-6/papilin regulates neuronal architecture

**DOI:** 10.1101/2025.02.10.637428

**Authors:** Malika Nadour, Robert I. Valette Reveno Leatis, Marie Biard, Noémie Frébault, Lise Rivollet, Philippe St-Louis, Cassandra R. Blanchette, Andrea Thackeray, Paola Perrat, Carlo Bevilacqua, Robert Prevedel, Laurent Cappadocia, Georgia Rapti, Maria Doitsidou, Claire Y. Bénard

**Author notes:** **Author Contributions**: Conceptualization, M.N., and C.Y.B.; Methodology, M.N., L.R., M.B., R.I.V., N.F., L.C., G.R., M.D., and C.Y.B.; Investigation, M.N., L.R., M.B., R.I.V., N.F., C.R.B., P.S., P.P., L.C, C.B., G.R., M.D., and C.Y.B. Writing - Original Draft, M.N. and C.Y.B.; Writing - Review & Editing, M.N., R.I.V., N.F., C.R.B, L.C., G.R., and C.Y.B.; Funding Acquisition, C.Y.B.; Resources, C.Y.B.; Supervision, M.N., L.R., and C.Y.B. **Competing Interest Statement:** The authors declare no competing interests. **Materials & Correspondence**: Correspondence and material requests should be addressed to C.Y.B.

## Abstract

Neuronal architecture established embryonically must persist lifelong to ensure normal brain function. However, little is understood about the mechanisms behind the long-term maintenance of neuronal organization. To uncover maintenance mechanisms, we performed a suppressor screen in *sax-7*/*L1CAM* mutants, which exhibit progressive disorganization with age. We identified the conserved extracellular matrix protein MIG-6/papilin as a key regulator of neuronal maintenance. Combining incisive molecular genetics, structural predictions, *in vivo* quantitative imaging, and cutting-edge Brillouin microscopy, we show that MIG-6/papilin remodels extracellular matrix collagen IV, working in concert with the secreted enzymes MIG-17/ADAMTS and PXN-2/peroxidasin. This remodeling impacts tissue biomechanics and ensures neuronal stability, even under increased mechanical stress. Our findings highlight an extracellular mechanism by which MIG-6/papilin supports the integrity of neuronal architecture throughout life. This work provides critical insights into the molecular basis of sustaining neuronal architecture and offers a foundation for understanding age-related and neurodegenerative disorders.

## INTRODUCTION

Proper nervous system function depends on both developing and maintaining the intricate architecture of its neural circuits. The dynamic processes of development and maturation of the nervous system begin *in utero* and extend well into adulthood ^1^. Key neuronal features, established earlier in development, must be preserved throughout life to ensure continuity in neuronal architecture and function ^2^. The structural organization of the nervous system faces numerous challenges, including physical stresses induced by the organism’s postnatal growth, anatomical remodeling, and the integration of new neurons, body movements, and injury ^3^. Failure to stably maintain neuronal architecture over the long-term can impair and compromise neuronal function and contribute to the manifestation of neurological conditions ^4,5^. Notably, a defining feature of many neurodegenerative diseases is the destabilization of axons and dendrites, and accompanying loss of synapses ^6–8^. Gaining insights into the mechanisms that preserve nervous system architecture could inform the development of therapeutics aimed at preventing or reversing such neurological conditions. Despite their importance, the mechanisms that sustain neuronal organization throughout life remain poorly understood.

A key contribution to the preservation of the structural integrity of multicellular assemblies, including those in the nervous system, comes the extracellular matrix (ECM) ^9^. Cells interact with the ECM and neighboring cells via cell adhesion molecules, and the coordinated actions of these molecules and the ECM are key for cellular behavior, enabling, for instance, neuronal structures to withstand physical stresses ^5^. However, how cellular regulation implicating ECM interactions evolve through time, or when subjected to physical challenges, remains elusive.

The ECM is of crucial importance in the nervous system, playing critical roles during development ^10^, but also in the adult mammalian brain, where the ECM constitutes approximately 20% of its volume. Diffuse ECM is found near and between synapses across the brain, and condensed ECM is organized as basement membranes associated with blood vessels, or as lattice-like structures called perineuronal nets, surrounding the soma and dendrites of several neuron types in multiple brain regions ^11^. These organized forms of ECM influence neuronal biology in the adult brain, including dendritic spine stability, synapse plasticity, and axon regeneration ^11,12^. Also, changes in ECM composition and structure have been linked to various neurological diseases (e.g., Alzeihmer’s disease and schizophrenia), as well as to brain injury and aging ^13,14^. This underscores the critical role of the neurons’ extracellular environment in maintaining normal neuronal physiology and its involvement in pathological conditions ^5^. However, our understanding of the long-term regulation of the ECM in the mature nervous system is limited and remains a dauting problem for neuropathology ^15^. For instance, how the intricate interactions between ECM proteases and their substrates, regulate ECM dynamics for sustaining neuronal structure and function is poorly understood.

The nematode *C. elegans* provides a powerful *in vivo* genetic model to study the lifelong maintenance of nervous system architecture, and particularly the role of the ECM in this process. A significant number of *C. elegans* neurons are organized into ganglia, with most neuronal processes running along major fascicles, such as the neuropil and the ventral nerve cord ^16^. The multicellular assemblies of ganglia and nerve cords are ensheathed by a specialized ECM, namely basement membranes ^17^. After hatching into a larva, *C. elegans* undergoes a nearly 100-fold increase in size until it reaches adulthood ^18^. Yet, the overall architecture of its nervous system is established during embryogenesis and remains largely intact throughout post-embryonic growth ^17,19^. Indeed, serial-section electron microscopy and connectome reconstruction of multiple *C. elegans* brains at successive stages of development revealed that the shape and positioning of most neurons and neurites established at birth remains consistent through adulthood, including approximately 70% of adult brain synapses being part of stable connections that are proportionally maintained from birth to adulthood ^19^. Individual neurons and ECM components can be readily visualized in living animals throughout their lifespan, using fluorescent reporters thanks to *C. elegans* transparency and small size ^20,21^. Combined with its genetic tractability, including using cell-specific promoters and conditional knockdowns, these features enable the investigation of the mechanisms sustaining nervous system architecture across its lifetime.

Thus, *in vivo* genetic studies using *C. elegans* has yielded critical insights into the mechanisms of the long-term maintenance of neuronal assemblies. These investigations have revealed post-natal molecular mechanisms that actively preserve neuronal organization, particularly within ganglia and nerve cords. Notably, several immunoglobulin superfamily molecules play crucial roles in maintaining the architecture of these multi-neuronal assemblies over time. These include the cell adhesion molecule SAX-7/L1CAM ^22–27^, the large ECM protein DIG-1 ^28,29^, the secreted two-immunoglobulin domain containing proteins ZIG-3 and ZIG-4 ^30,31^, and the ectodomain of the FGF receptor EGL-15 ^32^. Mutations in the genes encoding these molecules lead to neuronal defects that arise later in development, well after the normal initial establishment of neuronal morphology of the affected neurons. Strikingly, neuronal maintenance defects are suppressed by paralysis, highlighting that neuronal structures experience internal mechanical stress generated by body or organ movements, and that the identified neuronal maintenance molecules counteract these stresses, which otherwise would lead to neuronal disorganization ^28,31,33^. Orthologues of some of these molecules have been found to sustain neural circuits in other systems as well; for instance, in mice, knockout of L1CAM specifically in the adult brain results in behavioral deficits and synaptic transmission changes ^34^, and L1CAM maintains neocortical axo-axonic innervation into adulthood ^35^. While we have evidence that cell adhesion molecule SAX-7/L1CAM and ECM molecule DIG-1 play important roles in maintaining nervous system architecture, how the underlying ECM landscape and molecular interactions contribute to sustaining nervous system architecture throughout life is not understood. Given the extensive evolutionary conservation of ECM and neuronal cell surface molecules from worms to mammals, the maintenance mechanisms unravelled in *C. elegans* will provide insights on general principles by which the nervous system architecture is preserved lifelong.

To expand our understanding of the molecular mechanisms governing the preservation of neuronal architecture throughout life, we conducted a forward genetic suppressor screen in the *sax-7* mutant background, and identified *mig-6*, which encodes papilin, an extracellular matrix protein with structural similarities with ADAMTS metalloproteases ^36^. Papilin is conserved across metazoans, but its function remains elusive. Our findings reveal that MIG-6S/papilin is required post-developmentally for neuronal maintenance, by modulating the major extracellular matrix component collagen IV. Using Brillouin microscopy, we find that loss of *mig-6* changes the biomechanical properties of tissues in the region comprising the neurons. We demonstrate that MIG-6S/papilin cooperates with the ECM remodeling metalloproteinase MIG-17/ADAMTS in regulating the collagen IV network, and that both collagen IV levels and crosslinking are critical to sustain neuronal organization lifelong. Our study underscores the critical role of ECM regulation by the extracellular matrix protein papilin in preserving neuronal architecture throughout an organism’s lifetime, including under conditions of significant mechanical stress. We propose a model in which a balance between flexibility and adhesion, mediated by ECM remodeling and cell adhesion, ensures the structural stability of the embryonically established nervous system over time and improve its ability to withstand mechanical stress.

## MATERIALS AND METHODS

Please see Supplementary Information.

### Data Availability Statement

All data is available in the main text or the supplementary materials.

## RESULTS

### Loss of function of *mig*-*6*, which encodes the conserved extracellular matrix protein papilin, suppresses neuronal maintenance defects of *sax-7* mutants

To identify novel genes involved in the long-term maintenance of neuronal organization, we conducted a forward genetic screen. We refrained from searching directly for mutants with neuronal maintenance defects, as previous efforts using this approach invariably yielded numerous alleles of the large neuronal maintenance gene *dig-1* (^28^; C.Y.B., unpub. results). Rather, we reasoned that screening for suppressors of the defects of previously known neuronal maintenance mutants (**Fig. S1**), *sax-7*, would identify genes that directly or indirectly counteract defective long-term maintenance of neuronal architecture, providing insights into the basis of this process. In wild-type animals, the soma of chemosensory neurons ASH and ASI are located posterior to the nerve ring, where their axons project (neurons visualized with reporter P*sra-6::*DsRed2, **Fig. 1A**)^27^. This stereotypical positioning acquired during embryogenesis is preserved throughout life, making it a reliable indicator of neuronal organization. In *sax-7* mutants, although the soma and axons of ASH/ASI initially exhibit normal positioning during earlier development, they later become displaced from the 4th larval stage onward (**Fig. 1A,B,F**) ^22,25,27^, with the ASH/ASI soma ending up anteriorly displaced, and the nerve ring shifting posteriorly, resulting in the soma aligning with or even anterior to the nerve ring (**Fig. 1A**). In our F2 clonal genetic screen for suppressors of *sax-7* neuronal maintenance defects, we mutagenized *sax-7* mutants with ethyl methanesulfonate and screened F3 broods by fluorescence microscopy to find suppressors of the *sax-7* mutants ASH/ASI position defects. We isolated mutation *qv18*, which significantly suppressed the neuronal position defect in adult *sax-7(qv24)* animals, thus reducing the incidence of animals with mispositioned neurons (**Fig. S1**).

**Figure 1.**
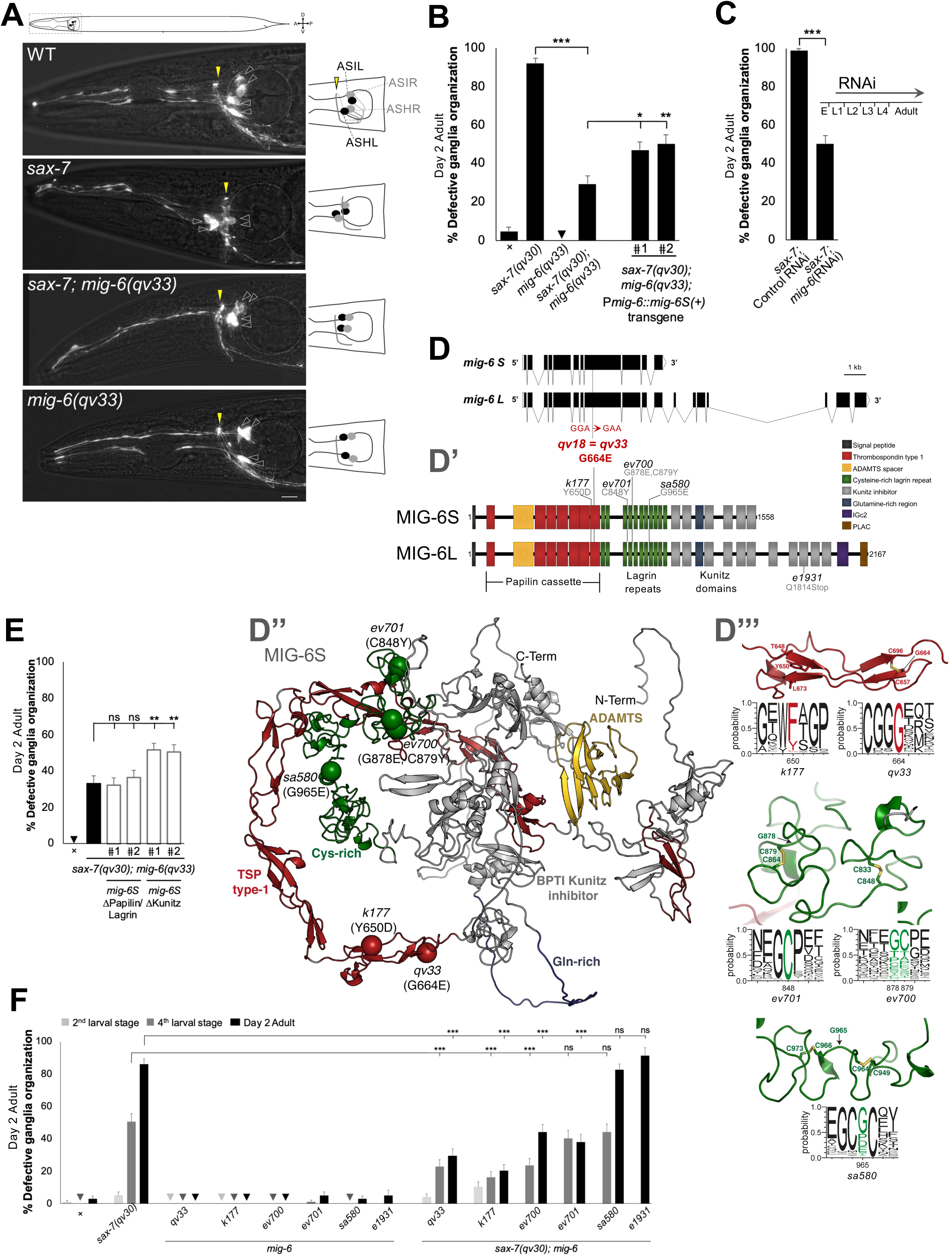
Loss of function of *mig*-*6*, which encodes the conserved ECM protein papilin, suppresses neuronal maintenance defects of *sax-7* mutants. (**A**) *qv33* is a newly identified allele of *mig-6* that suppresses the neuronal defects of *sax-7* mutants. Fluorescence images of the head region of 2-day-old adults (as indicated on worm schematics); chemosensory neurons ASH and ASI were visualized using reporter P*sra-6::DsRed2* (soma indicated by empty arrow heads; schematized on the right). In the wild type, the soma of the four neurons (each ASH and ASI pair has a soma in the left ganglion and in the right ganglion), are positioned posterior to the nerve ring (indicated by the yellow arrowhead) throughout life. In *sax-7* mutants, ASH and ASI soma are initially positioned normally but by the 4th larval stage or older they become progressively mispositioned relative to the nerve ring. Mutation *mig-6(qv33)* suppresses the neuronal disorganization of *sax-7(qv30)* mutants. Scale bar, 10 µm. (**B**) Quantification of the neuronal disorganization of ASH and ASI neurons (depicted in **A**), including in double mutants *sax-7; mig-6*, expressing a transgene of *mig-6S(+)* under the gene’s endogenous promoter which rescues the *mig-6*-mediated suppression of *sax-7-*neuronal disorganization defects. (**C**) Post-developmental depletion of *mig-6* by RNAi (from the 1st larval stage onwards) suppresses *sax-7* neuronal maintenance defects. (**D**) Schematic structure of the two isoforms of the gene *mig-6*, which encode a short and a long isoform of MIG-6/papilin (**D’**). The new allele *mig-6(qv33)* is a G664E amino acid substitution. Previously identified *mig-6* alleles are also indicated. (**D’’**) Overall structure of the MIG-6S protein predicted using AlphaFold version 2.3.1 implemented in ColabFold 1.5.2, with protein domains are colored as in **D’**. *qv33* and other mutations used are indicated. (**D’"**) Structural environment of residues Y650 (top left), G664 (top right), C848 (middle left), G878 and C879 (middle right), and G965 (bottom) in the structure of the MIG-6S protein predicted using ColabFold 1.5.2. These residues, and selected neighboring residues, are in stick representation. Residue conservation at each position is indicated through sequence logos generated using Weblogo 3.7.12 and an alignment of 250 protein sequences from Ecdysozoa species, including nematodes and arthropods, obtained using MMSeqs2 14-7e284. (**E**) Quantification of rescue assays with transgenes encoding recombinant versions of *mig-6S* (**Δ** indicates deleted domain). (**F**) Quantification of head ganglia organization phenotype in different *mig-6* alleles alone or in the *sax-7(qv30)* mutant background at the 2nd and 4th larval stages and 2-day-old adults. Age-progressive neuronal disorganization of *sax-7(qv30)* mutants is suppressed by some, but not all, *mig-6* alleles. Error bars are the standard error of the proportion. Comparisons made with z-tests; *P*-values were corrected by multiplying by the number of comparisons, Bonferroni correction. **<u>Note:</u>** Throughout all figures of this work, asterisks denote significant difference (\**P* ≤ 0.05, \*\**P* ≤ 0.01, \*\*\**P* ≤ 0.001); n.s., not significant; appropriate *post hoc* tests were performed for multiple comparisons (see **Materials and Methods**); “+” indicates wild-type strain. Sample sizes and source data in **Supplementary Information**.

Through whole genome sequencing and bioinformatic analyses ^37–39^, we identified as a candidate mutation a G-to-A transition at transcript nucleotide 1991 of the gene *mig-6*, which causes a glycine to glutamic acid substitution at residue 664 (**Fig. 1D’**). To determine if this mutation was the causal suppressor, we used CRISPR-Cas9 technology to reintroduce the candidate *qv18* molecular lesion in the *sax-7(qv30)* null mutant background ^27^. The resulting allele of *mig-6*, *qv33*, profoundly suppresses the *sax-7(qv30)* defects, reducing the percentage of affected animals from 90% to 30% (**Fig.1 A,B**), indicating that the suppressor mutation is indeed an allele of the gene *mig-6*. Except for gonad defects (similarly found in previously reported *mig-6* mutations) ^40^, single mutant *mig-6(qv33)* animals are fully viable and fertile, displaying a wild-type phenotype including for neurons ASH and ASI, overall nervous system morphology, body wall musculature, and pharynx (**Figs. 1A,B, S2, S6**). In contrast, null alleles of *mig-6* are sterile and embryonic and larval lethal ^40^. Knockdown of *mig-6* by RNA interference (RNAi) mimicked the effect of *mig-6(qv33)*, significantly suppressing *sax-7* neuronal defects (**Fig. 1C**). This result further confirms our molecular identification of the suppressor, and indicates that *qv33* is a loss-of-function mutation.

### The short isoform MIG-6S is key in neuronal maintenance, acting through its papilin cassette and lagrin repeats

The gene *mig-6* encodes a short isoform, *mig-6S*, and a long isoform, *mig-6L* (**Fig. 1D**) ^40^. We used five other *mig-6* alleles previously studied in the context of distal tip cell migration ^40^ to analyze their effect on *sax-7-*mediated neuronal maintenance (**Fig. 1D’, F**). An allele that specifically affects *mig-6L*, *e1931*, failed to suppress the neuronal maintenance defects in *sax-7(qv30); mig-6(e1931)* (**Fig.1F**), indicating that *mig-6L* is not implicated in this context. In contrast, other *mig-6* alleles that like *qv33* affect both the short and the long isoforms (*k177, ev700* and *ev701*) profoundly suppressed the ASH/ASI neuronal maintenance defects in *sax-7(qv30); mig-6* double mutant animals (**Fig.1F**). We further validated that *mig-6S* is the functional isoform in neuronal maintenance by performing rescue assays using a *mig-6S* transgene expressed under its endogenous promoter (minigene pZH125 ^40^). Since the loss of *mig-6* suppresses *sax-7* neuronal maintenance defects, restoration of *mig-6* function in double mutant animals *sax-7; mig-6* is expected to result in the reappearance of *sax-7* defects. Transgenic *sax-7(qv30); mig-6(qv33)* animals carrying wild-type transgenic copies of *mig-6S(+)* showed partial but significant rescue (**Fig. 1B**). The semi-dominant behavior of *mig-6* mutations described in other contexts ^40^ and in our analyses (below, **Figs. 3,5**) could explain the partial rescue. Collectively, these results firmly establish that *mig-6S* is central in neuronal maintenance.

*mig-6* encodes the conserved extracellular matrix MIG-6/papilin, which is orthologous to *Drosophila* and vertebrate papilin ^36,40–43^. Papilin is important for proper organogenesis in flies ^36^ and for gonad and pharynx development in worms ^21,40,44^ but its role in the nervous system remains largely unknown. Recently *papilin* was isolated in a screen for brain morphogenesis mutants, but its function awaits study ^45^, and in *C. elegans*, loss of *mig-6* was shown to affect PVD neuron 1o dendrite development ^46^. Little is known about the molecular mechanism of papilin function. MIG-6/papilin is a multidomain glycoprotein harboring thrombospondin type 1 (TSP1) repeats, cysteine rich lagrin repeats, and Kunitz protease inhibitor domains, among other domains (**Fig. 1D’** ^36,43^, and belongs to the ADAMTSL family of proteins (A Disintegrin and Metalloproteinase with Thrombospondin motifs-Like). Papilins, as other ADAMTSL proteins, are structurally related to ADAMTS metalloproteinases but lack the catalytic domain characteristic of ADAMTS, and no catalytic activity has been reported ^47,48^. Importantly, ADAMTSL proteins are characterized by the “papilin cassette" (**Fig. 1D’**), a region containing TSP1 domains and an ADAMTS spacer (homologous to non-catalytic domains of ADAMTS metalloproteinases) ^36,43^. Yet, the papilin cassette present in *Drosophila* papilin has been shown, *in vitro*, to bind and inhibit the activity of a procollagen N-proteinase, a vertebrate ADAMTS ^36^. In the *C. elegans* gonad, *mig-6* function influences the localization and levels of ADAMTS proteins MIG-17 and GON-1 ^21,40^. This raises the possibility that MIG-6/papilin may play important roles in the extracellular matrix via its papilin cassette.

As a first step to identify the critical domains of MIG-6S in neuronal maintenance, we modeled the molecular consequences of these tested *mig-6* alleles on MIG-6S. The *mig-6(qv33)* mutation results in a substitution of a glycine to a glutamic acid at amino acid 664 (G664E), located in a TSP1 repeat in the papilin cassette. A sequence alignment as well as the predicted structure of MIG-6S, modeled using ColabFold (**Fig. 1D", S3**), shows that G664 is a highly conserved residue located at the entrance of a beta-strand (**Fig. 1D"’**). Replacement of a glycine with a larger residue in the *qv33* mutant is likely to cause a steric clash with the disulfide bond formed by residues C657 and C696 (**Fig. 1D’"**) and could alter a potential binding of MIG-6S to the ECM, or other interactions.

Similarly, other *mig-6* alleles that suppress *sax-7* defects (**Fig. 1F**) affect residues located in the papilin cassette or in the adjacent lagrin repeats (**Fig. 1D’, D"**). Indeed, *mig-6(k177)* (Y650D) missense allele impacts a residue in a TSP1 domain of the papilin cassette. In the ColabFold predicted structure of MIG-6S, the aromatic portion of Y650 interacts with both L673 and T648 (**Fig. 1D’’’**). Consistently, homologs of MIG-6S typically possess either a tyrosine or a phenylalanine at position Y650 (**Fig. 1D’’’**). Missense allele *mig-6(ev701)* (C848) affects a residue in a lagrin repeat of MIG-6S, where it forms a disulfide bridge with C833 (**Fig. 1D’’’**), perhaps explaining the strong conservation of a cysteine residue at this position. Finally, missense allele *ev700* (harboring substitutions G878E and C879Y) affects two residues located within the lagrin repeats cysteine-rich region of MIG-6S that are also well conserved, with C879 forming a disulfide bridge with C864 (**Fig. 1D’’’**). Thus, mutation at G878 and C879 could perturb the pattern of disulfide bridges within this region. Finally, missense allele *mig-6(sa580)* (G965E) affects a glycine residue located in a lagrin domain, which could disrupt the adjacent cysteine residues involved in interactions with C949 and C973 (**Fig. 1D’"**). However, *mig-6(sa580)* does not supress *sax-7* neuronal defects (**Fig. 1F**), highlighting the specific effects of distinct *mig-6* mutations. In sum, these analyses support the idea that the papilin cassette and neighboring lagrin repeats are crucial for the function of MIG-6S in neuronal maintenance.

To experimentally validate which are the most critical regions of the MIG-6S protein in the context of neuronal maintenance, we carried out rescue assays using recombinant versions (**Fig. 1E, S4**). A recombinant *mig-6S* transgene lacking the sequence that encodes the C-terminal region of the protein, which contains the Kunitz domains, retained rescuing activity in *sax-7(qv30); mig-6(qv33)* double mutant animals with reappearance of neuronal disorganization, similar to the full length *mig-6S(+)* (compare to **Fig.1B**). This supports that the Kunitz domains are not essential for MIG-6S function in neuronal maintenance. In contrast, a recombinant *mig-6S* transgene lacking the sequence that encodes the N-terminal region containing the papilin cassette and lagrin repeats failed to rescue the function of *mig-6* in *sax-7(qv30); mig-6(qv33)* double mutants. These results, together with our analyses of a series of mutant alleles (molecular impact in **Fig. 1D** and functional consequences in **Fig. 1F**), demonstrate that the papilin cassette and the nearby lagrin repeats are the key domains for MIG-6S functionality in neuronal maintenance.

### *mig-6/papilin* modulates neuronal maintenance post-embryonically and in specific neuronal contexts

Since several *mig-6* mutations suppress the progressive head ganglia disorganization of *sax-7* mutants, we asked whether losing *mig-6* function after the development of ASH/ASI neurons would be sufficient to suppress *sax-7* neuronal maintenance defects. As ASH/ASI neurons complete their development in embryogenesis, we depleted *mig-6* function post-embryonically by feeding *sax-7(qv30)* animals with *mig-6*(RNAi) bacteria starting from the mid/late-L1 larval stage onward. Our results showed that the post-embryonic depletion of *mig-6* function strongly suppressed *sax-7* ASH/ASI neuronal maintenance defects (**Fig. 1C**), highlighting its post-developmental in maintaining ASH/ASI neuronal architecture.

We next explored if *mig-6*’s impact on neuronal maintenance is context-specific or generalized across the nervous system. *sax-7* mutants are known to exhibit defects in maintaining axon positioning along the ventral nerve cord: axons of bilateral neurons PVQL/R initially develop normally, projecting ipsilaterally during embryogenesis, but later get displaced to the opposite fascicle of the ventral nerve cord in *sax-7* animals ^25^, coinciding with remodeling of the underlying tissue during late first larval stage ^30,49^. We found that *mig-6(qv33)* partially suppresses these *sax-7* axon “flip-over” defects (**Fig. 2A, B**), indicating that *mig-6* participates in the maintenance of nerve cord organization as well.

**Figure 2.**
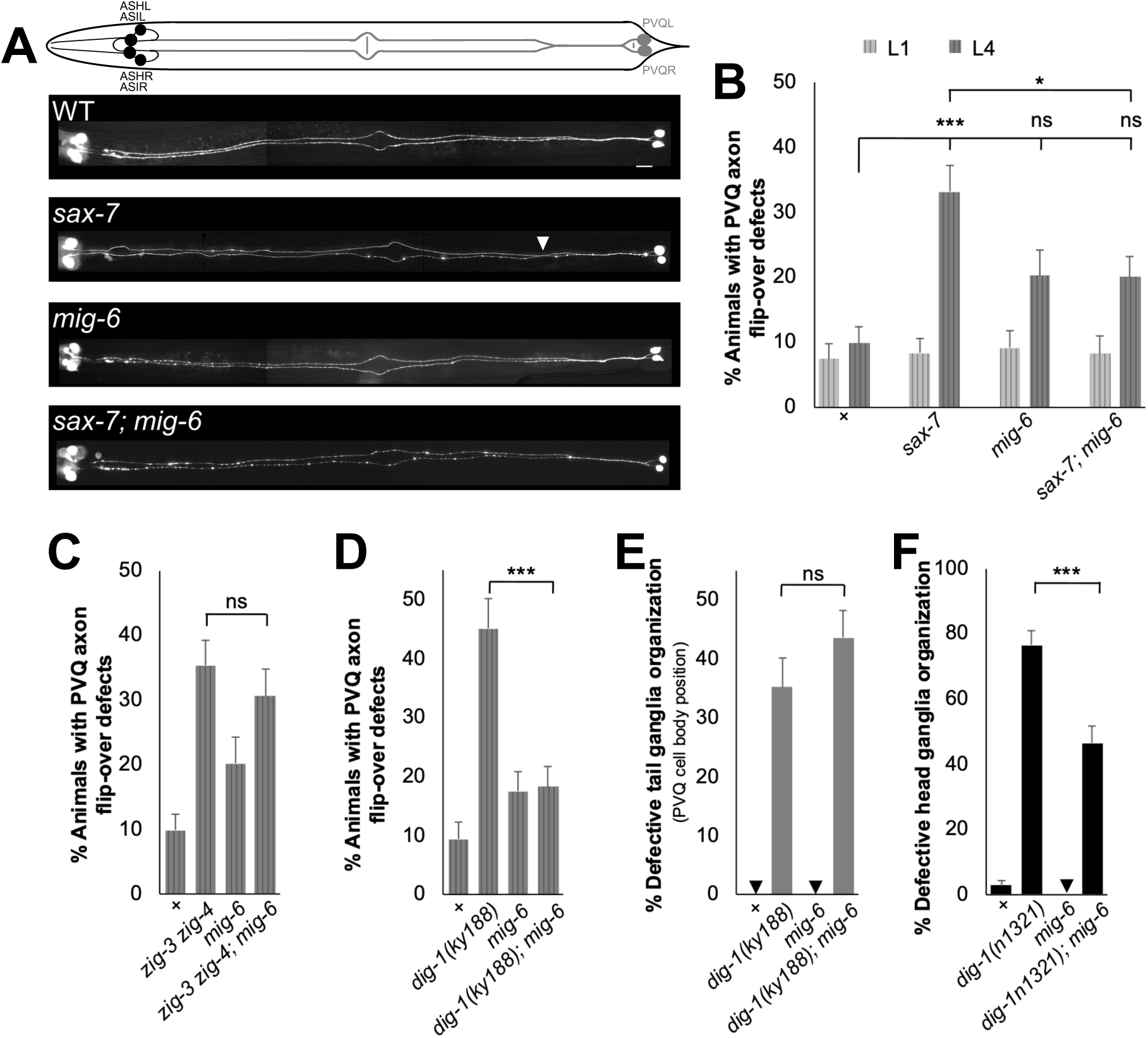
*mig-6/papilin* impacts the maintenance of axon and soma position in specific neuroanatomical and genetic contexts. (**A**) Fluorescence images of 4th larval stage (L4) animals; PVQ neurons visualized using P*sra-6::DsRed2*. In the wild type, the axon of each of two PVQ neurons (L and R) extends ipsilaterally along the ventral nerve cord during embryogenesis and remains on the ipsilateral side throughout life. In *sax-7* mutants, whereas PVQ axons develop normally and are positioned like the wild type in early 1st larval stage (L1) animals (not shown), they later become displaced to the opposite side of the ventral nerve cord (L4 shown). Scale bar, 20 µm. (**B,C,D**) Quantification of PVQ axons (L4 animals) shows that *mig-6*(*qv33*) suppresses the axonal defects of *sax-7(qv30)* and *dig-1(ky188)* mutants, but not of *zig-3(tm924) zig-4(gk34)*. (**E, F**) *mig-6(qv33)* does not supress PVQ soma displacement in the lumbar ganglia in the tail of *dig-1(ky188)* L4 animals, but partially supresses the ASH and ASI soma position relative to the nerve ring in head ganglia of *dig-1(n1321)* examined as 2-day-old adult age animals. Error bars are the standard error of the proportion; z-tests were performed.

The small secreted two-immunoglobulin proteins ZIG-3 and ZIG-4 also function to maintain axon position in the *C. elegans* ventral nerve cord ^30,31,33^, as does the giant basement membrane protein DIG-1, which is also required for ganglia maintenance ^28,29^. To determine if disrupted *mig-6* function could also suppress the defective maintenance of axon position in these other known neuronal maintenance mutants, we generated mutant combinations between *mig-6* and either *zig-3* and *zig-4*, or *dig-1*. We found that *mig-6(qv33)* did not supress the axon flip-over defects in the double mutant of small secreted two-immunoglobulin proteins *zig-3(tm924) zig-4(gk34)* (**Fig. 2C**), highlighting the specificity of *mig-6* effects. However, *mig-6(qv33)* did suppress axon flip-over in *dig-1(ky188)* mutants (*ky188* is the *dig-1* allele with the most penetrant axonal defects; **Fig. 2D**). As *dig-1* mutants also exhibit defects in the maintenance of both tail and head neuronal ganglia organization, we examined the impact of *mig-6(qv33)* on *dig-1* ganglia organization maintenance. We found that loss of *mig-6* did not supress defective soma positioning of PVQ neurons in the tail ganglia of *dig-1(ky188)* mutants (*ky188* display progressive and penetrant PVQ soma displacement; **Fig. 2E**), but did partially suppress head ganglia organization in *dig-1(n1321)* mutants (*n1321* is the most severe *dig-1* allele for head ganglia; **Fig. 2F**). These results indicate that the role of *mig-6* in neuronal maintenance is specific and context-dependent, varying with the type of neuronal maintenance molecule affected and neuronal structure.

### *mig-6/papilin* functions non-autonomously, together with *mig-17/ADAMTS* to impact neuronal maintenance

Given that disrupting the function of the extracellular matrix protein MIG-6/papilin counteracts the progressive neuronal disorganization observed in head ganglia and the nerve cord of *sax-7* and *dig-1* mutants, we hypothesized that the absence of functional MIG-6/papilin protein would affect the ECM surrounding these neuronal structures. In the developing gonad, *mig-6* genetically interacts with *mig-17/ADAMTS*, which encodes a secreted metalloprotease of the ADAMTS family ^40^, thought to remodel the gonadal basement membrane ^21,50–53^. We therefore sought to investigate the functional relationship between *mig-6/papilin* and *mig-17* in neuronal maintenance. We first examined *mig-17(k174)* putative null mutant animals ^54^ (null allele used throughout this study) and found that ASH and ASI neurons are normally positioned with respect to the nerve ring (**Fig. 3A,C**). *mig-17* mutants exhibit an elongated pharynx ^55^ (**Fig. S5**), but this did not affect neuronal position with respect to body length. Indeed our data show that there is no difference in the relative positions of the nerve ring, or of ASH and ASI soma, between mutants with normal pharynx length (such as *mig-6(qv33)*) and *mig-17* mutants (**Fig. S5**), indicating that neuronal positioning is independent from pharynx length. To test if loss of the ECM remodeling molecule *mig-17/ADAMTS* would impact the *sax-7* head ganglia disorganization, we looked at double mutants lacking both *sax-7* and *mig-17*. In double mutants *sax-7; mig-17,* only 27% of animals display head ganglia disorganization, compared to ∼90% in *sax-7* single mutants (**Fig. 3A,C**). This result shows that, similar to loss of *mig-6*, loss of *mig-17* significantly supresses the neuronal maintenance defects in *sax-7* mutants. In contrast, loss of another secreted ADAMTS metalloprotease, ADT-2/ADAMTS, involved in *C. elegans* body size regulation and cuticle structure ^56^, did not supress *sax-7* defects nor enhance their suppression by *mig-6* mutation (**Fig. 3C**). This highlights the specific role of *mig-17/ADAMTS* in neuronal maintenance.

**Figure 3.**
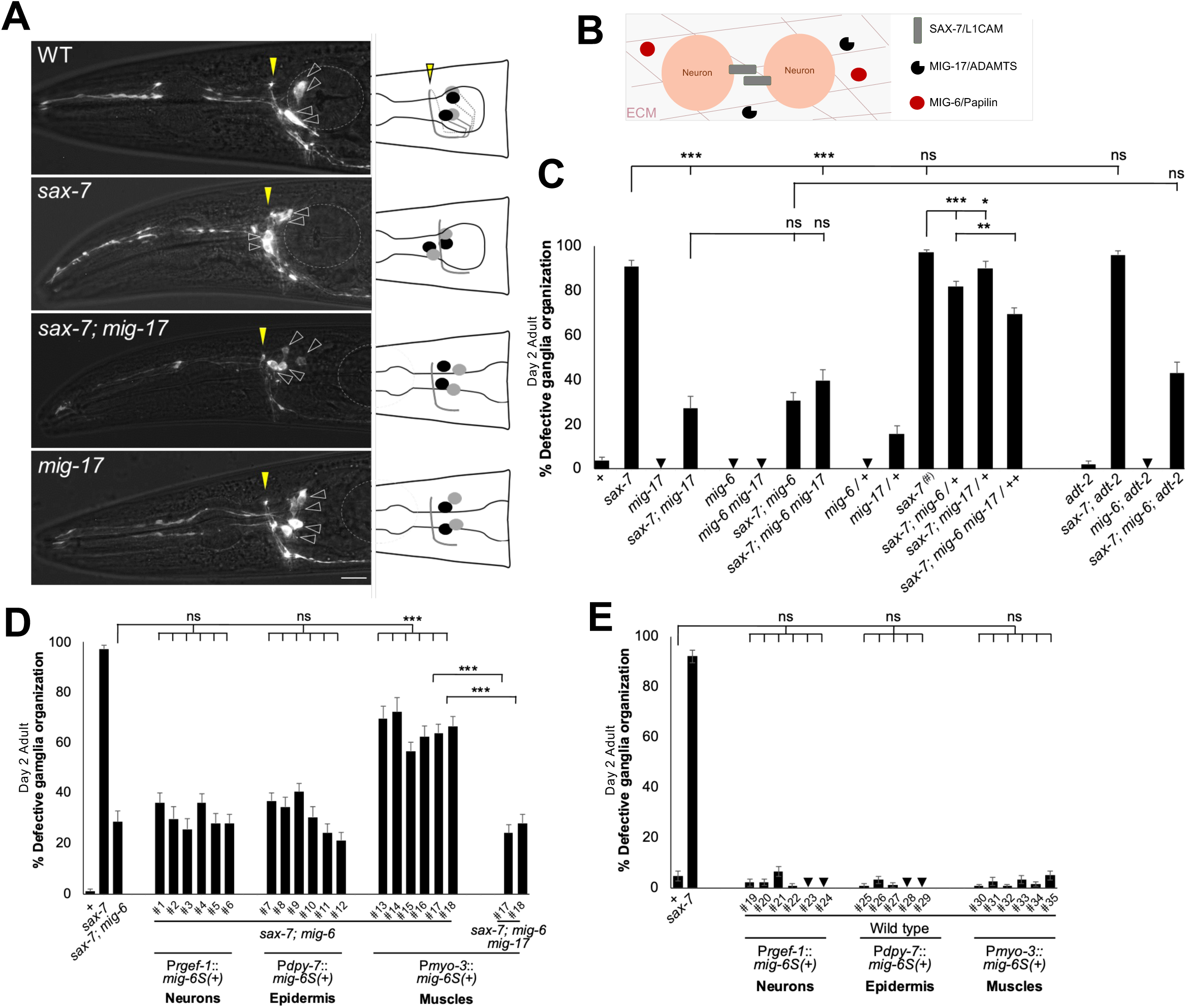
*mig-6*/*papilin* and *mig-17*/ADAMTS function together in neuronal maintenance. (**A**) Fluorescence images of the head region of 2-day-old adults (schematized on the right), with soma and axons of neurons ASH and ASI visualized using P*sra-6::DsRed2* (as in Fig. 1). Scale bar, 10 µm. (**B**) A model proposing functional cooperation between MIG-17/ADAMTS and MIG-6/papilin in the ECM to mediate neuronal maintenance. (**C**) Quantification of ASH and ASI displacement in 2-day adults of wild type, *sax-7(qv30)*, *mig-6(qv33)*, *mig-17(k174)*, and *adt-2(wk156)* single mutants, and their combinations, as homozygous or heterozygous (written as *"mig-6*" or *"mig-6/+*", respectively). ’#’ indicates an additional control for *sax-7*, with *dpy-11* in the background, as *dpy-11* was used to distinguish heterozygous animals (see **Supplementary Information** for full genotypes). Similar to the effect of some *mig-6* mutations, loss of *mig-17* also suppressed *sax-7* neuronal maintenance defects (but not loss of *adt-2*). Simultaneous loss of *mig-6* and *mig-17* did not enhance the suppression of *sax-7* defects. (**D**) Quantification of assays to rescue the function of *mig-6* with tissue-specific expression of *mig-6S(+)*. (**E**) Quantification of control assays showing that *mig-6S(+)* transgenic expression in wild-type animals, including from body wall muscles, does not affect neuronal organization. Error bars are the standard error of the proportion; z-tests.

To then determine whether *mig-6* and *mig-17* function in the same genetic pathway in this context, we constructed a triple *sax-7; mig-6(qv33) mig-17(k174)* mutant strain. We found that the simultaneous loss of *mig-6* and *mig-17* did not further enhance the suppression of neuronal maintenance defects in *sax-7* mutants (**Fig. 3C)**, suggesting that *mig-6* and *mig-17* may function in the same pathway to maintain neuronal architecture. To further probe the notion that *mig-6* and *mig-17* function collaboratively to impact neuronal maintenance, we analyzed the effect of partially losing the function of both genes using double heterozygous animals *mig-6(qv33) mig-17(k174) / mig-6(+) mig-17(+),* abbreviated as *mig-6 mig-17/++*. Single heterozygous animals *mig-17(k174)*/+ only slightly supressed *sax-7* neuronal defects (**Fig. 3C**). Meanwhile, single heterozygous *mig-6(qv33)/+* mildly suppressed *sax-7* defects; **Fig. 3C**), consistent with observations that *mig-6* alleles (*ev700*, *ev701* and *k177*) behave in a semi-dominantly due to haploinsufficiency during gonad and PVD neuron development ^40,46^. Notably, in double heterozygous animals *sax-7; mig-6 mig-17/++*, the suppression of *sax-7* defects was significantly greater, with only 69% of animals showing defects, compared to 81% in *sax-7; mig-6/+* animals (**Fig. 3C**). This result indicates that the two extracellular matrix genes *mig-6/papilin* and *mig-17/ADAMTS* act within the same pathway to influence the long-term maintenance of neuronal organization.

Given that the neurons’ environment controls their maintenance, *mig-6/papilin* may be expected not to function from the neurons themselves. Indeed, extracellular matrix components, including MIG-6, are produced by mesodermal cells, particularly body wall muscles, and the epidermis ^21,40,44,57,58^. We thus generated transgenic *sax-7; mig-6* double mutants animals expressing wild-type *mig-6S* under different tissue-specific promoters: P*rgef-1* for neurons, P*dpy-7* for epidermis (hyp7), and P*myo-3* for mesodermal cells (body wall muscles). Since the loss of *mig-6* suppresses *sax-7* neuronal maintenance defects, restoring *mig-6* function is expected to lead to the reappearance of neuronal disorganization in *sax-7; mig-6* double mutants. We found that expression of *mig-6S(+)* in the neurons or the epidermis did not rescue *mig-6* function. In contrast, expression of *mig-6S(+)* in the body wall muscles in *sax-7; mig-6* double mutant animals robustly rescued the neuronal maintenance defects, with an increase of neuronal defects from 29% up to 72%, depending on the transgenic line (**Fig. 3D**). This indicates that *mig-6* functions cell non-autonomously from muscles to impact neuronal maintenance. As a control, we expressed these transgenes in wild-type animals and observed no neuronal defects (**Fig. 3E**), ruling out the possibility that *mig-6S(+)* overexpression by muscles induces artefactual neuronal disorganization. Overall, this confirms that the neuronal defects observed in transgenic animals expressing *mig-6S(+)* in body wall muscles represent *bona fide* rescue of *mig-6* function.

We then investigated whether *mig-17* is necessary for *mig-6*’s function in neuronal maintenance. To test whether the loss of *mig-17* would affect the ability of *mig-6S(+)* transgene to rescue, we generated *sax-7; mig-6 mig-17* triple mutants carrying transgenic lines expressing wild-type *mig-6S(+)* from body wall muscles (lines #17 and 18 used above, **Fig. 3D**). Unlike the successful rescue of *mig-6* function observed in *sax-7; mig-6* transgenic animals, the loss of *mig-17* prevented the *mig-6S(+)* transgene from rescuing neuronal maintenance defects, as the transgenic triple mutant *sax-7; mig-6 mig-17* animals did not exhibit a reappearance of neuronal defects (**Fig. 3D**). This may indicate that the normal function of MIG-6/papilin depends on MIG-17/ADAMTS, or alternatively, an excess of MIG-6S cannot compensate for the loss of MIG-17. In conclusion, we find that *mig-6/papilin* acts non-autonomously from muscles to suppress *sax-7* neuronal maintenance defects in a manner that requires the function of *mig-17/ADAMTS*.

### Loss of *mig-6S/papilin* results in increased EMB-9/collagen IV levels that accumulates as extracellular fibrotic-like structures

Given that MIG-6/papilin is an extracellular ADAMTS-like protein which interacts functionally with MIG-17/ADAMTS, we hypothesized that the *mig-6* mutation may suppress *sax-7* neuronal maintenance defects by modulating the extracellular environment, including nearby neurons. To directly test whether loss of *mig-6* leads to changes in the extracellular matrix, we analyzed the distribution of a key ECM component, EMB-9/collagen IV α1 (hereafter referred to as ’EMB-9/collagen IV’), known to genetically interact with *mig-6* during gonadal development ^40^ or to be altered in the gonadal basement membrane in *mig-6*(RNAi)-treated animals ^21^. In *C. elegans*, collagen IV, similar to its vertebrate counterparts, is a heterotrimeric molecule consisting of two EMB-9 (α1-like) chains and one LET-2 (α2-like) chain, previously shown to colocalize ^59–62^. We therefore used a P*emb-9*::EMB-9::mCherry fluorescent reporter ^63^ (gift from David Sherwood) to examine the distribution of collagen IV, focusing particularly on the head region (**Fig. 4**). In *mig-6(qv33)* mutants, like in the wild type, we observed collagen IV signal along the contour of the pharynx and the surface of body wall muscles (**Fig. 4A, Fig. S6**), corresponding to the basement membranes of these structures ^21^, as well as in spherical accumulations within muscle cells where collagen IV is produced. However, in *mig-6* mutants, the overall abundance of EMB-9/collagen IV is higher compared to the wild type (**Fig. 4A**; see **Fig. S7** for quantification of collagen IV in entire head region including muscle cells), including inside muscle cells. Moreover, *mig-6* mutants display notable enrichments of collagen IV, often elongated, which we have termed "fibrotic-like structures" (**Fig. 4A,B**; **Fig. S6**). These fibrotic-like structures are extremely rare in the wild type and seen only in aging adults (5-day old adults), but are detected as early as the 1st larval stage in *mig-6(qv33)* mutants, and by the 4th larval stage and adulthood, 100% of the *mig-6* mutants present fibrotic collagen IV (**Fig. 4A,B**). These fibrotic-like structures in *mig-6* mutants are typically located in the posterior region of the head (**Fig. 4C,D**). Interestingly, our repeated observations of the same *mig-6(qv33)* animals in a longitudinal analysis over several days (**Fig. 4E**, n=8) show that the *mig-6* mutants’ collagen IV fibrotic-like structures stably persist over time. Measuring the size of these structures at different developmental stages further confirmed that they lengthen in an age-progressive manner (**Fig. 4F**).

**Figure 4.**
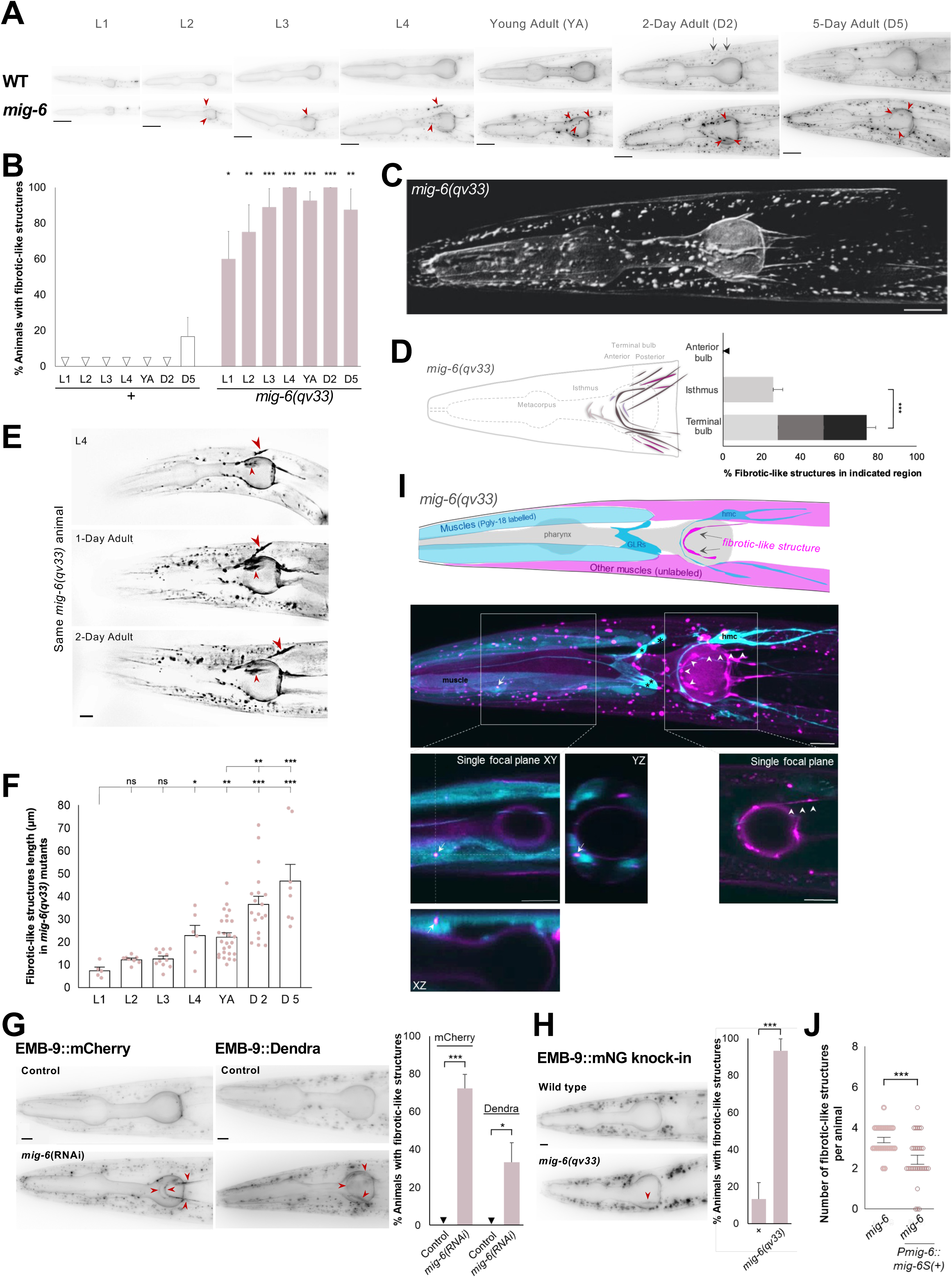
Loss of *mig-6/papilin* function disrupts proper extracellular collagen IV remodeling and causes fibrotic structures. (**A**) Fluorescence images of animals expressing collagen IV reporter P*emb-9*::EMB-9::mCherry at larval stages L1, L2, L3, and L4, and adult ages in control and *mig-6(qv33)* mutants. Thin lines of collagen IV located in the basement membrane of muscle cells are observed in wild-type and *mig-6* animals (indicated by two black arrows), and *mig-6* fibrotic-like structures are indicated by arrowheads. Scale bar, 20 µm. (**B**) Percentage of animals displaying collagen IV fibrotic-like structures at different ages in the wild type and *mig-6(qv33)* mutants expressing EMB-9::mCherry. (**C**) 3D-isosurface renderings (made with Imaris) of collagen IV/EMB-9::mCherry in the head region of a 2-day-old adult *mig-6(qv33)* mutant animal. Scale bar, 20 µm. (**D**) *mig-6* mutants’ collagen IV fibrotic-like structures are typically located in the posterior region of the head, in the general area near the terminal bulb and the posterior of the isthmus of the pharynx. Schematic compilation of fibrotic-like structures’ positions in *mig-6(qv33)* mutants (n>100), and percentage in indicated antero-posterior head regions. These "pharyngeal regions" serve as antero-posterior landmarks only, as these fibrotic-like structures localize in the extracellular space outside the pharynx; also, fibrotic-like structures only occasionally contact the basement membrane of the pharynx (see videos of fluorescence images in **Supplementary Information**). (**E**) Longitudinal analysis of a *mig-6(qv33)* mutant animal expressing EMB-9::mCherry repeatedly examined as a L4, 1-, and 2-day-old adult shows that the collagen IV fibrotic-like structures (indicated by arrowheads) persist over time. Scale bar, 20 µm. (**F**) Average length of the fibrotic-like structures per *mig-6(qv33)* mutant animal expressing EMB-9::mCherry at different ages. (**G**) Fluorescence images and quantification of animals displaying collagen IV fibrotic-like structures visualized by EMB-9::mCherry or EMB-9::Dendra in 2-day-old adult animals of wild-type animals treated with *mig-6*(RNAi) or empty vector control. Scale bar, 10 µm. (**H**) Collagen IV fibrotic-like structures are also detected in 2-day-old-adult *mig-6* mutants using the endogenous collagen IV reporter *qy24* EMB-9::mNG knock-in. Scale bar, 10 µm. (**I**) Fluorescence image of a young adult *mig-6* mutant, expressing EMB-9::mCherry (magenta), along with *dnIs13* P*gly-18::gfp* (cyan) that labels a subset of head wall muscles, the head mesodermal cell (hmc), and glial cells (GLR, indicated by asterisks); a schematic rendering is shown above. ColIagen IV (EMB-9::mCherry reporter) is observed as intracellular dots or accumulations inside muscle cells (indicated by the arrow on single focal planes on XY, XZ and YZ); and long extracellular fibrotic-like structures (one such structure is pointed to by white arrowheads on main image and on the focal plane on the right panel, which does not coincide with the cytoplasmic extensions of the hmc, nor with other mesodermal cells such as muscles or GLR cells). Magenta circle on single planes is signal from the basement membrane of the pharynx, which also contains collagen IV. (**J**) Rescue of the collagen IV fibrotic-like structures phenotype of *mig-6(qv33)* mutants expressing EMB-9::mCherry. Expression of the *mig-6S(+)* minigene under its own promoter decreases the number of fibrotic-like structures. Error bars are the standard error of the proportion (B, D, G, H) or of the mean (F, J). z-tests (in B, D, G, H), ANOVA in F), or non-parametric Wilcoxon test (in J).

We next studied the localization of the collagen IV fibrotic-like structures that occur in *mig-6* mutants. In *C. elegans*, collagen IV is produced by body wall muscles and other mesodermal cells, including the head mesodermal cell (hmc) and GLR glia ^64,65^. We thus generated a strain of *mig-6(qv33)* mutants carrying both the EMB-9::mCherry reporter and *dnIs13 gly-18p::gfp* ^66^ to simultaneously label collagen IV and anterior head wall muscles, the hmc, and GLR glia, respectively. Confocal microscopy revealed that collagen IV fibrotic-like structures in *mig-6* mutants do not overlap with muscle cells, nor with hmc, nor the GLR glia, and are located extracellularly (**Fig. 4I**). In addition, spherical collagen IV deposits seen in body wall muscles, appear to be more intense in *mig-6* mutants than in wild type (**Fig. 4A**; **Fig. S6**), and these are indeed intracellular (**Fig. 4I**).

To further support our findings, given that mCherry protein fusions can form aggregates ^67^, we employed an alternative multicopy reporter, EMB-9::Dendra2 ^63^ (gift of David Sherwood). With this reporter we also observed that EMB-9 accumulates as fibrotic-like structures in 33% of animals following *mig-6*(RNAi) knockdown (**Fig. 4G**). Importantly, using an endogenous CRISPR knock-in reporter for collagen IV ^21^, EMB-9::mNG, 90% of the *mig-6(qv33)* mutants show fibrotic-like structures (**Fig. 4H**).

Since collagen IV accumulates in *mig-6* mutants, we assessed the stability of EMB-9::mCherry by fluorescence recovery after photobleaching (FRAP), which we performed on muscle and pharyngeal basement membranes that contain collagen IV, present in both the wild type and *mig-6(qv33)* mutants (**Fig. S8A**). Our FRAP measurements revealed no significant difference between *mig-6* mutants and the wild type, with very limited collagen IV recovery across different time points (**Fig. S8B**), supporting the notion that loss of *mig-6* does not affect collagen IV short-term dynamics *per se*, which is consistent with its described stable association with the gonadal basement membrane ^21^.

We further strengthened our findings that loss of *mig-6* alters collagen IV levels and distribution by looking at collagen IV in other *mig-6* loss-of-function backgrounds. We generated animals with the *k177* allele of *mig-6* (Y650D mutation in the same domain as *qv33* G664E) carrying EMB-9::mCherry. Similar to *qv33* animals, *k177* mutants exhibit a significant accumulation of EMB-9/collagen IV, with approximately 80% of adult animals displaying fibrotic-like structures (**Fig. 6A,B; Fig. S6**). Similarly, RNAi-mediated knockdown of *mig-6* resulted in 72% of adult animals exhibiting EMB-9::mCherry fibrotic-like structures (**Fig. 4G**). In contrast, specifical loss of *mig-6L*, using allele *e1931*, did not alter the collagen IV pattern nor led to fibrotic-like structures (**Fig. 6A,B; Fig. S6**). These results suggest that *mig-6S*, but not *mig-6L*, is essential for proper extracellular collagen IV organization in the head region. Overall, consistent with the findings of papilin affecting collagen IV in the gonadal basement membrane ^21^, we found that the formation of extracellular collagen IV fibrotic-like structures in the head region is a robust phenotype linked to the loss of *mig-6S/papilin* function.

### *mig-6*/papilin affects the biomechanical properties of the animal’s tissues

Our findings demonstrate that disruption of MIG-6/papilin results in a dramatic collagen IV fibrotic phenotype (**Fig. 4**) and that the progressive neuronal disorganization in head ganglia and the nerve cord of *sax-7* and *dig-1* mutants is counteracted in *mig-6* mutants (**Fig. 1 and 2**). We therefore hypothesized that the environment surrounding these neuronal structures may be modified in *mig-6(qv33)* mutants in such a way that results in enhanced maintenance of neuronal architecture. To start addressing this possibility, we characterized the biomechanical state of tissues in *mig-6(qv33)* mutants by measuring their viscoelasticity properties using Brillouin microscopy. This label-free imaging technique allows for the assessment of the viscoelastic properties of biological samples through photon–phonon scattering interactions ^68^. The key parameters measured are the Brillouin scattering induced frequency shift and linewidth, which provide information on the high-frequency longitudinal modulus and therefore the elastic and viscous properties of the sample, respectively ^69–71^. We first ensured that the refractive indexes of the head region were similar between *mig-6* and wild-type animals at the examined ages (**Fig. S9A**), which is an important prerequisite to render the Brillouin microscopy results comparable to one another. Next, using a confocal Brillouin microscope ^72^, we imaged a head region at the level of the studied neuronal cell bodies, which includes other cells such as other neurons, muscles, epidermis, glia, as well as the associated basement membranes/ECM (**Fig. 5A,B**). Our findings show a significant decrease in Brillouin shift in *mig-6* mutants compared to the wild type at the L4 larval stage (**Fig. 5C,D; Fig. S9B**), indicating a reduction in tissue elasticity. This change was not detected at the earlier L2 stage, suggesting that the loss of *mig-6* affects tissue elasticity more prominently at later developmental stages. The decreased elasticity was most pronounced in the posterior zone of the region of interest (ROI), which exhibited lower Brillouin elastic contrast (**Fig. S9C**). Furthermore, Brillouin linewidth measurements revealed that tissue viscosity was also reduced in *mig-6* mutants at both L2 and L4 stages (**Fig. 5C,D; Fig. S9B**). To assess tissue viscoelasticity in wild-type and *mig-6* mutants, we measured the Brillouin loss tangent parameter ^73^, which showed decreased tissue viscoelasticity in *mig-6* mutants compared to controls at both the L2 and L4 stages (**Fig. 5C,D; Fig. S9B**). These results demonstrate that MIG-6/papilin is essential for maintaining proper tissue mechanical properties *in vivo* during age-progression. Furthermore, as the altered biomechanical properties of the tissues in the head of *mig-6* mutants are detected earlier than the appearance of ASH/ASI neuronal disorganization in *sax-7* mutants, this suggests that properties of the underlying head ganglia environment play an important role in maintaining neuronal architecture over time.

**Figure 5.**
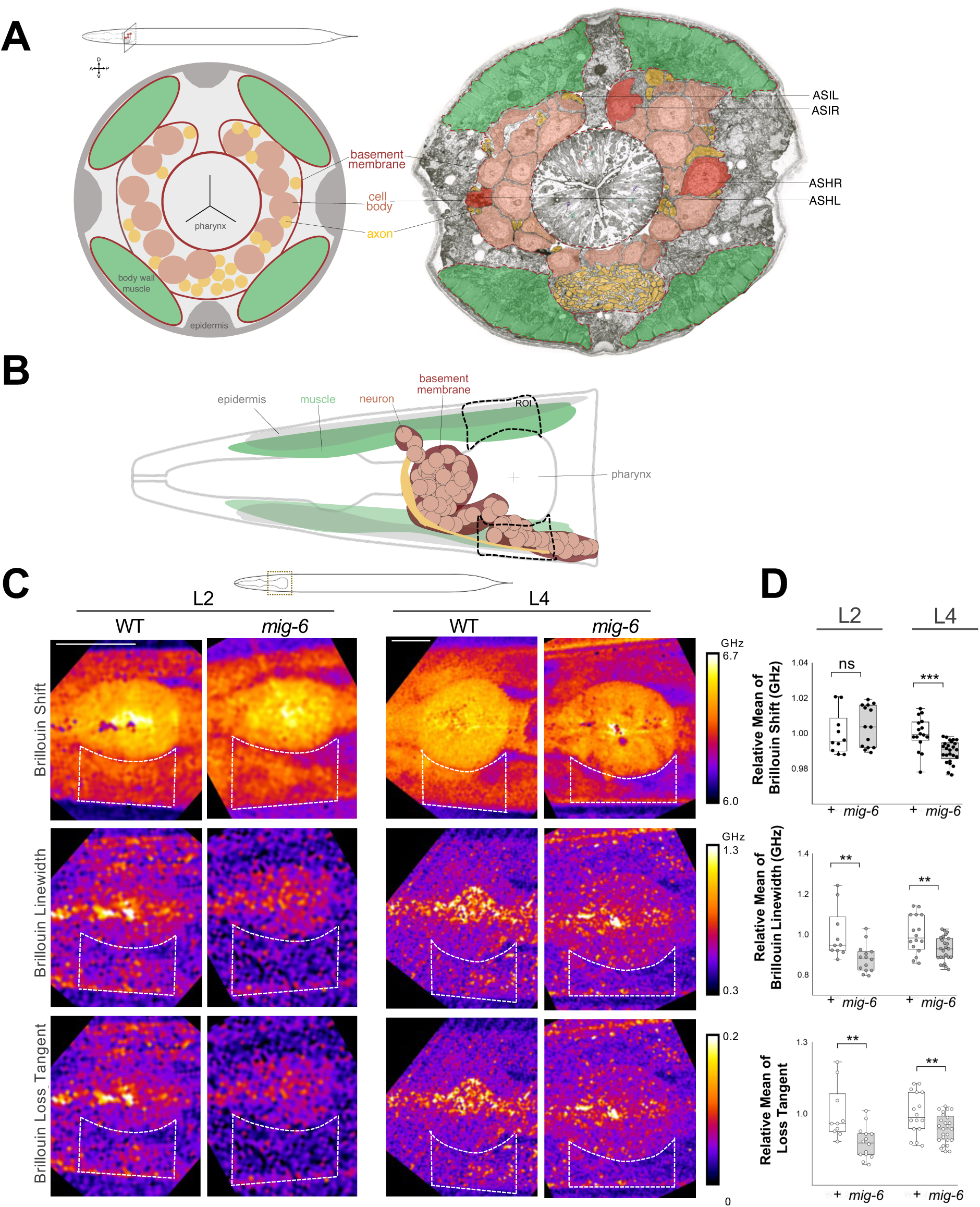
Loss of *mig-6/papilin* alters tissue biomechanical properties. (**A**) Colored electron micrograph of a cross section of an adult wild-type animal in the head region near the location of the ASHL/R and ASIL/R somas (White, Southgate et al. 1986); modified from WormAtlas.org). A schematic of the cross section is provided on the left. Neuronal cell bodies are pseudo-colored in peach, with ASHL/R and ASIR in darker peach; axons in yellow; muscles in green; the location of the basement membranes surrounding body wall muscles, pharynx, and head ganglia have been drawn in to provide context (dark red dashed lines). (**B**) Schematics of the *C. elegans* head region, highlighting prominent cell types in the region imaged by Brillouin microscopy, including neurons of head ganglia, which are surrounded by basement membrane, as well as the ROIs. (**C**) Brillouin microscopy shift, linewidth, and loss tangent images of the head region of L2 and L4 larval stages of wild-type and *mig-6(qv33)* animals. ROIs were drawn flanking the terminal pharyngeal bulb and include several tissues and cell types (muscle, epidermis, neurons, and glia), as well as basement membranes. Scale bar, 20 µm. The color bars indicate absolute values of the shift, linewidth, and loss tangent. (**D**) Quantification of ROIs shows a decrease in elasticity (Brillouin shift), viscosity (Brillouin linewidth), and loss tangent (viscoelasticity) in *mig-6(qv33)* mutants compared to wild type. Mean values of the ROIs were normalized to the wild type (for absolute values, see **Fig. S9** and **Supplementary Information**). Error bars are the standard error of the mean; t-tests were performed.

### *mig-6/papilin* and *mig-17/ADAMTS* function together to regulate extracellular collagen IV

Building on the previous finding that *mig-6* genetically interacts with *mig-17/ADAMTS* to suppress *sax-7* neuronal defects (**Fig. 3**), we investigated whether *mig-6* and *mig-17* also functionally interact in regulating collagen IV distribution. *mig-17(k174)* single mutants exhibit fibrotic-like structures in 75% of young adults and 95% of 2-day old adults (**Fig. 6A,B; Fig. S6**), similar to the phenotype observed in *mig-6*(*qv33*) and *mig-6*(*k177*) mutants. The simultaneous loss of *mig-6* and *mig-17* in double homozygous *mig-6 mig-17* mutants did not enhance the collagen IV fibrotic-like structures compared to single mutants, neither in penetrance (**Fig. 6B**), nor in expressivity (**Fig. 6D**), consistent with the notion that the extracellular matrix genes *mig-6* and *mig-17* function within the same pathway to regulate collagen IV distribution.

To reinforce this conclusion, we assessed the effect of simultaneously losing a single functional copy of each gene, which can provide insights into genetic interactions (avoiding the ceiling effect, as heterozygous animals are much less penetrant for this phenotype). In *mig-6(qv33)/+* single heterozygotes, we observed collagen IV fibrotic like-structures in approximately 10% of young adults, increasing to 75% of 2-day-old adults (**Fig. 6A,B; Fig. S6**), indicating a semi-dominant effect of *mig-6* also for this phenotype. In *mig-17(k174)/+* single heterozygotes, only around 10% of animals displayed fibrotic like-structures at both ages. Remarkably, in young adult double heterozygotes *mig-6 mig-17/++*, as many as 80% exhibited collagen IV fibrotic-like structures (increasing to 85% in 2-day-old adults; **Fig. 6A,B; Fig. S6**). Also, the number of fibrotic-like structures per animal strikingly increases in the double heterozygous *mig-6 mig-17/++* animals compared to single heterozygous (**Fig**. **6C**). Together, these results firmly establish that *mig-6* and *mig-17* functionally interact to regulate extracellular collagen IV.

**Figure 6.**
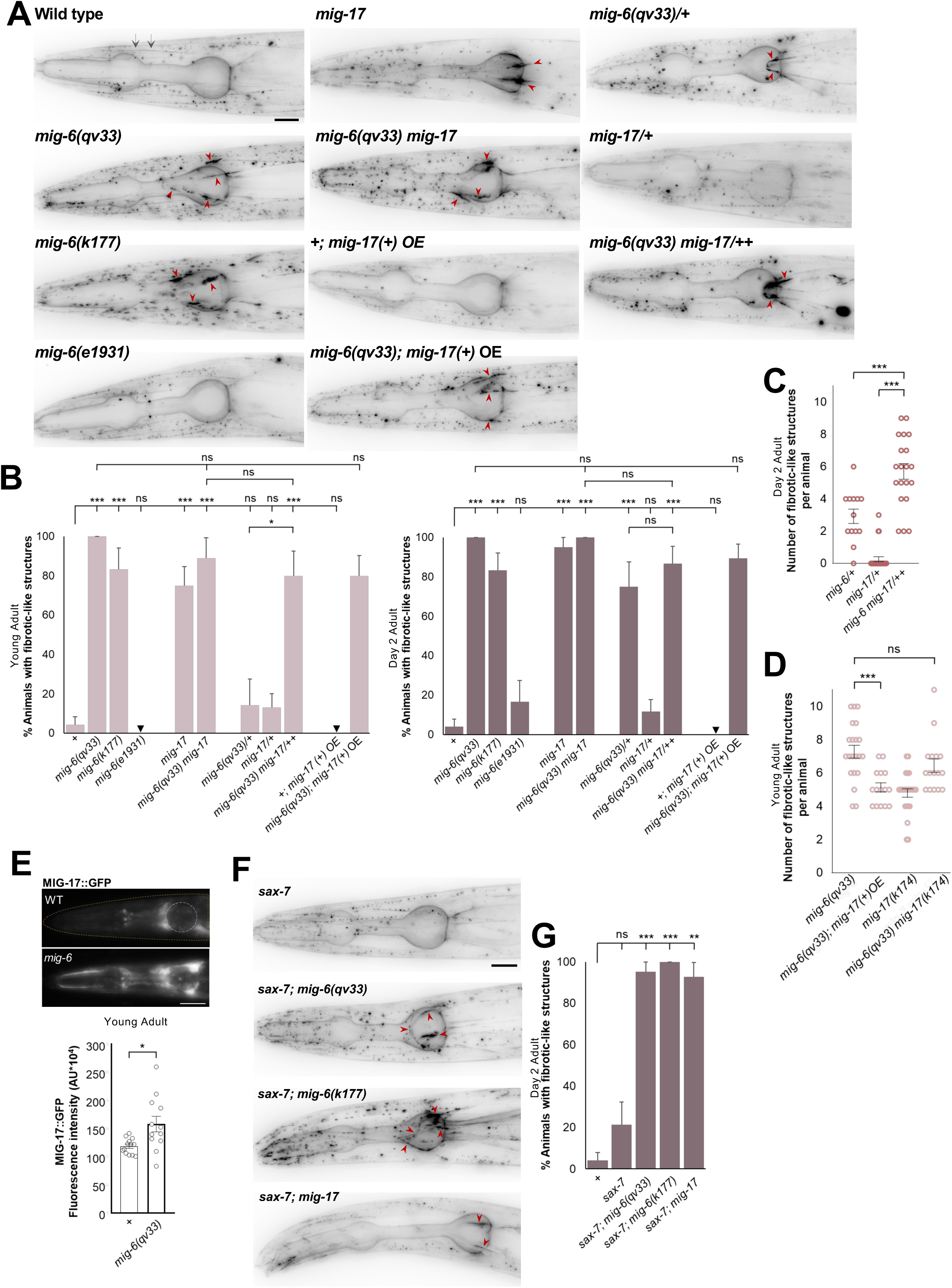
*mig-6/papilin* and *mig-17/ADAMTS* both regulate extracellular collagen IV remodeling. (**A**) Fluorescence images of the head region of 2-day-old adult animals using EMB-9::mCherry. Thin lines of collagen IV located in the basement membrane of muscle cells are observed in all genotypes (indicated by two black arrows); red arrowheads indicate fibrotic-like structures in mutants. Mutant animals for *mig-6*(*qv33* or *k177*) display collagen IV fibrotic-like structures and prominent intracellular accumulations (dots). *mig-17(k174)* mutants also exhibit fibrotic-like structures. In contrast, *mig-6L(e1931)* allele behaves like the wild type. Scale bar, 20 µm. (**B**) Quantification of the percentage of animals displaying collagen IV fibrotic-like structures in young adult and 2-day-old adult animals expressing EMB-9::mCherry. (**C**) Quantification of the number of collagen IV fibrotic-like structures observed per 2-day-old adult heterozygous animal expressing EMB-9::mCherry. (**D**) Quantification of the number of collagen IV fibrotic-like structures observed per young adult animal expressing EMB-9::mCherry. (**E**) Fluorescence images of MIG-17::GFP in the head region of young adults, quantified as fluorescence intensity in head region drawn in orange. Circle, terminal bulb of the pharynx for reference. Scale bar, 24 µm. (**F**) Fluorescence images of the head region of 2-day-old adults in *sax-7(qv30)* null mutant background, using reporter EMB-9::mCherry. Scale bar, 20 µm. (**G**) Quantification of the percentage of 2-day-old adult animals displaying collagen IV fibrotic-like structures. Error bars are the standard error of the proportion (in B, G) or of the mean (in C, D, E). z-tests in B and G, Wilcoxon Mann-Whitney test in C, ANOVA in D, and t-test in E. A.U., arbitrary units. OE, overexpression of wild-type copies. Mutant alleles are only indicated on images and graphs when several alleles of a given gene are used; thus, unless specified otherwise, "*mig-6*" is *mig-6(qv33)* and "*mig-17*" is *mig-17(k174)* throughout this work. Homozygous genotypes are written as *"mig-6*" and heterozygous as *"mig-6/+*".

MIG-17/ADAMTS has been hypothesized to function as a metalloprotease that degrades collagen IV ^74,75^. We thus asked whether overexpression of wild-type copies of *mig-17(+)* impacts the collagen IV phenotype of *mig-6* mutants. For this we used functional transgene P*mig-17::mig-17(+)::gfp* (Nishiwaki et al, 2000; Jafari et al, 2010) (**Fig. 6A** and **Fig. S6**), combining the integrated transgene (*evIs213*, gift of Joe Culotti) with *mig-6(qv33)*. We found that while the percentage of *mig-6* animals displaying fibrotic-like structures was unchanged (**Fig. 6B**), the number of fibrotic-like structures per young adult animal significantly decreases in *mig-6* mutants overexpressing the *mig-17(+)* transgene (**Fig. 6D**). This result indicates that the collagen IV defects of *mig-6* mutants can be partially offset by *mig-17/ADAMTS* overexpression.

As *mig-6* mutants display a fibrotic collagen IV phenotype, and the overexpression of *mig-17(+)* partially suppresses this defect (**Fig. 6D**), we wondered whether levels of MIG-17/ADAMTS may be lowered in *mig-6* mutants. To test this idea, we examined the distribution of MIG-17/ADAMTS in *mig-6(qv33)* mutants using P*mig-17::*MIG-17::GFP (*evIs213*). We found that instead of being decreased, MIG-17 is in fact upregulated in *mig-6* mutants (**Fig. 6E**), with new enrichments in different head regions compared to wild type. Given that increased *mig-17(+)* levels can at least partially suppress *mig-6* mutants (**Fig. 6D**), and that MIG-17 levels are elevated in *mig-6* mutants (**Fig. 6E**), together these results suggest that MIG-17/ADAMTS’s function requires functional MIG-6/papilin to robustly ensure normal collagen IV distribution.

### Suppression of *sax-7* neuronal maintenance defects upon loss of *mig-6/papilin* and *mig-17/ADAMTS* depends on collagen IV levels and cross-linking

Because *mig-6* and *mig-17* cooperate to regulate extracellular collagen IV (**Fig. 6A-D**) and maintain neuronal organization in *sax-7* mutants (**Fig. 3A,B,C**), we asked if the suppression of *sax-7* neuronal maintenance defects upon loss of *mig-6* or *mig-17* is linked to changes in collagen IV patterning in the ECM. We assessed the state of collagen IV distribution in the *sax-7* mutant background and found that *sax-7* single mutants behave like the wild type in this regard (**Fig. 6F,G; S6**). Also, double mutants *sax-7; mig-6(qv33)* and *sax-7; mig-6(k177)*, as well as *sax-7; mig-17(k174)* displayed a fibrotic-like structure phenotype like that of the respective *mig-6* or *mig-17* single mutants (**Fig. 6F,G; S6**). That *sax-7; mig-6* and *sax-7; mig-17* double mutants exhibit modified collagen IV pattern (**Fig. 6F,G**) and maintain organized head ganglia (**Fig. 3C**) is consistent with the notion that the state of the ECM plays a role in maintaining neuronal architecture. In line with this, *mig-6L*-specific allele *e1931* does not affect the pattern of collagen IV (normal levels and no fibrotic structures; **Fig. 6A,B; S6**) and does not suppress the neuronal maintenance defects of *sax-7* mutants (**Fig. 1F**).

We then investigated whether collagen IV levels and distribution contribute to the maintained neuronal organization of double mutants *sax-7; mig-6* and *sax-7; mig-17*. We depleted *emb-9/collagen IV* by RNAi treatment of animals from the 1st larval stage and examined head ganglia organization in adults. This *emb-9*(RNAi) knockdown effectively depleted EMB-9/collagen IV levels (**Fig. 7A, Fig. S10A**), and importantly, did not affect neuronal organization in wild-type animals, nor in single mutants. In contrast, depleting collagen IV reversed the suppression of *sax-7* neuronal defects by *mig-6* or *mig-17* mutation. Indeed, double mutant animals *sax-7; mig-6* and *sax-7; mig-17* showed increased neuronal disorganization upon *emb-9*(RNAi) (**Fig. 7B**). This result indicates that collagen IV levels are key for the suppression of *sax-7* neuronal defects by the loss of *mig-6* or *mig-17*. We then tested whether a higher level of collagen IV could mimic the effect of *mig-6* loss of function in suppressing *sax-7* neuronal maintenance defects. We found that *sax-7* animals overexpressing transgene *emb-9(+)* (in *sax-7; qyIs46* animals carrying the multicopy transgene P*emb-9::*EMB-9::mCherry) did not show suppression of neuronal maintenance defects (**Fig. 7C**). Thus, while sustained levels of collagen IV are required to suppress *sax-7* neuronal defects, elevated collagen IV level *per se* is insufficient to ensure neuronal maintenance.

**Figure 7.**
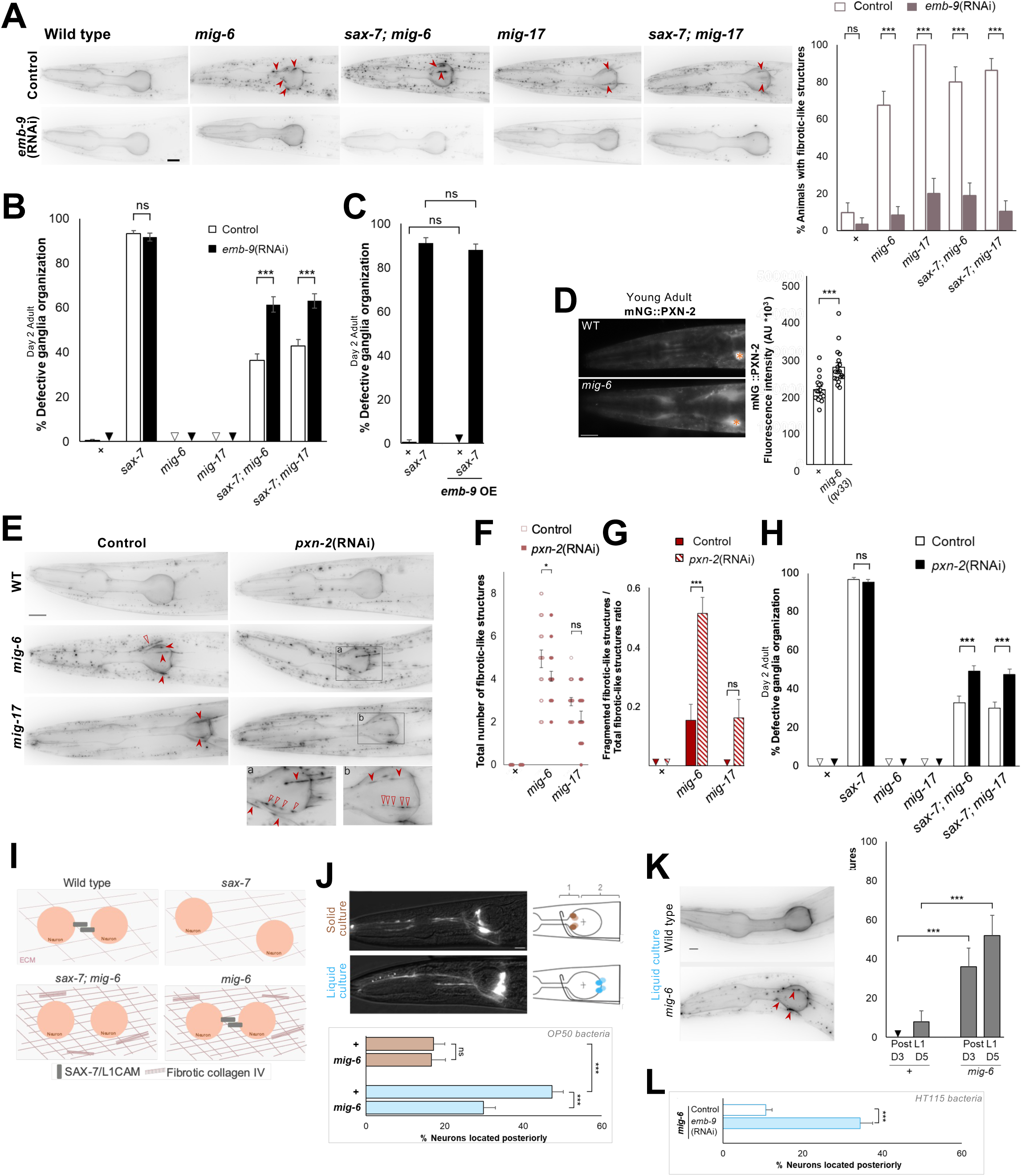
Extracellular collagen IV networks are key for *mig-6/papilin*’s role in neuronal maintenance, including under conditions of increased stress. (**A**) Fluorescence images of the head region of 2-day-old adults using alleles *mig-6(qv33), sax-7(qv30)* and *mig-17(k174)*, expressing EMB-9::mCherry, subjected to control (empty vector) or *emb-9*(RNAi) from the L1 stage; fibrotic-like structures indicated by red arrowheads. Quantification of the percentage of animals with fibrotic-like structures (see **Fig. S10A** for quantification of collagen IV levels). (**B**) Knockdown of collagen IV by *emb-9*(RNAi) reverses the suppression of *sax-7(qv30)* neuronal defects by loss of *mig-6(qv33)* or *mig-*17*(k174)*. Quantification of neurons ASH and ASI displacement (as in Fig. 1) in 2-day-old adults subjected to control (empty vector) or *emb-9*(RNAi) since the L1 stage. (**C**) Increase of *emb-9*/collagen IV levels alone (overexpression using *qyIs46* multicopy integrated transgene P*emb-9*::EMB-9::mCherry) is not sufficient to suppress *sax-7(qv30)* neuronal defects. (**D**) Fluorescence images of mNG::PXN-2 in wild-type and *mig-6(qv33)*, scale bar, 20 µm; and quantification (see **Fig. S12** for ROI and additional images). A.U., arbitrary units. (**E**) Knockdown of collagen IV crosslinking enzyme PXN-2/peroxidasin by *pxn-2*(RNAi) reduces the number of fibrotic-like structures and alters their state. Fluorescence images of collagen IV (*qyIs46* EMB-9::mCherry) in 2-day-old adults of wild type and mutants *mig-6(qv33)* and *mig-17(k174)*, subjected to control (empty vector) or *pxn-2*(RNAi) from the L1 stage. Insets (a and b) show detail of fibrotic-like structures that appear continuous (arrowheads) or fragmented (empty arrowheads). Scale bars, 20 µm. (**F**) Quantification of the number of fibrotic-like structures per animal expressing EMB-9::mCherry in wild type, *mig*-*6(qv33)*, and *mig-17(k174)* mutant animals in control (empty vector) and *pxn-2*(RNAi) conditions. Both fragmented and non-fragmented fibrotic-like structures are included in this quantification of fibrotic like-structures number. (**G**) Quantification of the ratio of fragmented to total fibrotic-like structures per animal in wild type and mutants *mig-6(qv33)* or *mig-17(k174)* expressing EMB-9::mCherry in control (empty vector) and *pxn-2*(RNAi)-treated conditions. (**H**) The suppression of *sax-7(qv30)* neuronal defects by mutations *mig-6(qv33)* or *mig-17(k174)* is reversed by deficient collagen IV crosslinking in *pxn-2*(RNAi)-treated animals. Quantification of the displacement of neurons ASH and ASI in 2-day old adult animals subjected to control (empty vector) or *pxn-2*(RNAi). (**I**) Summary of the relationship between ECM remodeling and neuronal maintenance in the context of *sax-7* and *mig-6* mutants. Loss of *mig-6* compensates for loss of neural adhesion molecule SAX-7 through extracellular remodeling, ensuring the maintenance of neuronal organization. (**J**) Fluorescence images of the head region of wild-type adults fed regular *E. coli* OP50 and imaged in 2-day adults when grown on solid media, or at the equivalent age of 5 days-post L1 hatching when grown in liquid. Neurons ASH and ASI were visualized using reporter P*sra-6::DsRed2*; drawings illustrate their soma position in wild-type animals grown on solid (brown), or in liquid (blue, notice the posterior placement of neurons when worms swam in liquid conditions). Scale bar, 10 µm. Quantification of neuronal placement in wild-type and *mig-6(qv33)* adults, when grown in solid (2-day adults) or liquid conditions (5 days-post L1 hatching). In each animal, neurons were considered posteriorly displaced when at least one of the four soma was in area 2 (posterior to the pharyngeal grinder, indicated by the cross). (**K**) Fluorescence images of collagen IV (EMB-9::mCherry) in wild type and *mig-6(qv33)* adults at 5 days-post L1 hatching, grown in liquid since L1 hatching. Scale bar, 10 µm. Quantification of the percentage of animals displaying collagen IV fibrotic-like structures in wild-type and *mig-6(qv33)* adults grown in liquid conditions since hatching (examined at 5 days-post L1 hatching). (**L**) Animals were grown in liquid conditions since hatching while being subjected to RNAi treatment; animals were fed *E. coli* HT115 bacteria harboring the empty vector (control RNAi) or the *emb-9*(RNAi) vector to deplete collagen IV. Quantification of neuronal placement (as in C) in wild-type and *mig-6(qv33)* adults, examined at 5 days-post L1 hatching. Collagen IV depletion prevents the stabilizing effect of *mig-6* mutation. Error bars are the standard error of the proportion (z-tests in A-C, H, J-L) or of the mean (t-test in D, Wilcoxon Mann-Whitney test in F and G).

We therefore investigated whether collagen IV organization plays a role in neuronal maintenance as well. Collagen IV molecules form complex crosslinked networks involving dimerization through their NC1 domain ^65,76,77^. The extracellular enzyme peroxidasin catalyzes sulfilimine S=N bonds between collagen IV NC1 domains ^76,78^, which are essential for collagen IV networks and basement membrane integrity ^79–82^. The *C. elegans* genome encodes two peroxidasins, of which PXN-2/peroxidasin is known for its effects on the ECM and genetic interactions with collagen IV genes ^83^. We examined the pattern of PXN-2/peroxidasin using a knock-in fluorescent reporter mNeonGreen::PXN-2 (driven under the *pxn-2* promoter ^21^, and found that the expression of PXN-2/peroxidasin is upregulated in *mig-6(qv33)* mutants compared to wild type (**Fig. 7D**), suggesting that *mig-6* is implicated in the regulation of peroxidasin 2 levels in the ECM. Knockdown of *pxn-2/peroxidasin* by RNAi (from the 1st larval stage) did not significantly lower the penetrance of fibrotic-like structures (**Fig. S10B**), but significantly reduced the number of fibrotic-like structures (**Fig. 7E,F**), and led to a striking increase of fragmented fibrotic collagen IV (**Fig. 7E,G**). Importantly, *pxn-2*(RNAi) knockdown reverses the suppression of *sax-7* mutants’ neuronal defects by loss of *mig-6* or *mig-17* (**Fig. 7H**). Together, these results highlight that the function of MIG-6/papilin and MIG-17/ADAMTS in neuronal maintenance is dependent on collagen IV and its crosslinking by the peroxidasin enzyme. Further, they support the notion that the elevated levels of crosslinked collagen IV in the ECM of *mig-6* mutants contribute to stabilizing neuronal position and maintaining neuronal architecture in *sax-7* mutants (**Fig. 7I**).

### Loss of *mig-6*/papilin counteracts neuronal disorganization induced by increased mechanical stress

Since loss of *mig-6* function positively impacts the maintenance of neuronal organization in animals lacking the cell adhesion molecule SAX-7/L1CAM, we hypothesized that it may also support neuronal architecture in otherwise wild-type animals that experience increased internal mechanical stress. We used the distinct locomotion patterns of *C. elegans* to probe this question. In liquid media, *C. elegans* swims, whereas on solid media, it crawls. Swimming and crawling differ in neuromuscular activity and speed, with swimming being faster ^84,85^. The associated exerted forces are also different: when the worm crawls on solid media, the forces exerted on the worm’s cuticle are higher than when swimming in liquid ^86–88^. By contrast, swimming worms perform many more body bends, with more numerous body wall muscles contractions, which is expected to result in higher mechanical stress on internally located neurons (e.g., in head ganglia) compared to the slower movements of worms crawling on solid media. We therefore subjected worms to continuous swimming in liquid culture from the time of hatching to adulthood, and assessed the position of sensory neurons ASH and ASI. Compared to the neurons in animals grown on solid medium, neuronal position in wild-type animals changed significantly when grown in liquid medium for 5 days, exhibiting a significant posterior displacement (**Fig. 7J**). However, *mig-6(qv33)* mutant animals raised in liquid medium showed a significant decrease in neuronal displacement (**Fig. 7J**), indicating that loss of *mig-6/papilin* function results in enhanced maintenance of neuronal organization upon high internal mechanical stress.

Because collagen IV is required for the stabilizing effect of *sax-7* mutants neuronal organization by loss of *mig-6* function, we examined the state of collagen IV in swimming animals. We observed that *mig-6* mutants display an altered collagen IV pattern (similar to that of *mig-6* mutants grown on solid medium), with the presence of fibrotic-like structures, increasing in penetrance from day 3 to day 5 post L1 hatching (**Fig. 7K**). We then asked if collagen IV was required for the neuronal protective effect conferred by the *mig-6* mutation in otherwise wild-type animals when grown in liquid. Depleting collagen IV by *emb-9*(RNAi) of swimming animals significantly weakened the *mig-6*-mediated stabilizing effect of neuronal organization of ASH and ASI neurons (**Fig. 7L**). This result indicates that collagen IV is critical in the neuronal protective mechanism involving *mig-6/papilin* in conditions of increased mechanical stress.

## DISCUSSION

Neuronal architecture established embryonically must persist throughout life to ensure nervous system function. However, the mechanisms sustaining neuronal organization over the long term remains poorly understood. This work uncovers a novel mechanism where ECM dynamics plays a critical role in maintaining neuronal architecture. Through a multidisciplinary approach, integrating forward genetic screening, incisive molecular genetic analysis, structural molecular predictions, quantitative live imaging, and measurement of biomechanical properties by Brillouin microscopy, we have identified the evolutionarily conserved extracellular matrix protein MIG-6/papilin as a key regulator of the long-term maintenance of the neuronal architecture. We show that MIG-6/papilin impacts neuronal maintenance by modulating the animal’s tissues biomechanical properties and remodeling the extracellular network of collagen IV, which is a major component of the basement membranes, including those surrounding neuronal assemblies. We also find that ECM metalloproteinase MIG-17/ADAMTS is important for sustaining neuronal architecture, and functionally cooperates with MIG-6/papilin in ECM remodeling in order to enable the long-term stability of neuronal architecture. Both the abundance and the cross-linking of collagen IV networks are essential for the MIG-6/papilin-remodeled ECM state that enables the maintenance of neuronal structures. Thus, this work reveals a previously unknown mechanism by which ECM remodeling enables the preservation of neuronal architecture (**Fig. 8**), in the face of age-progressive stresses, to preserve continuous neural function.

**Figure 8.**
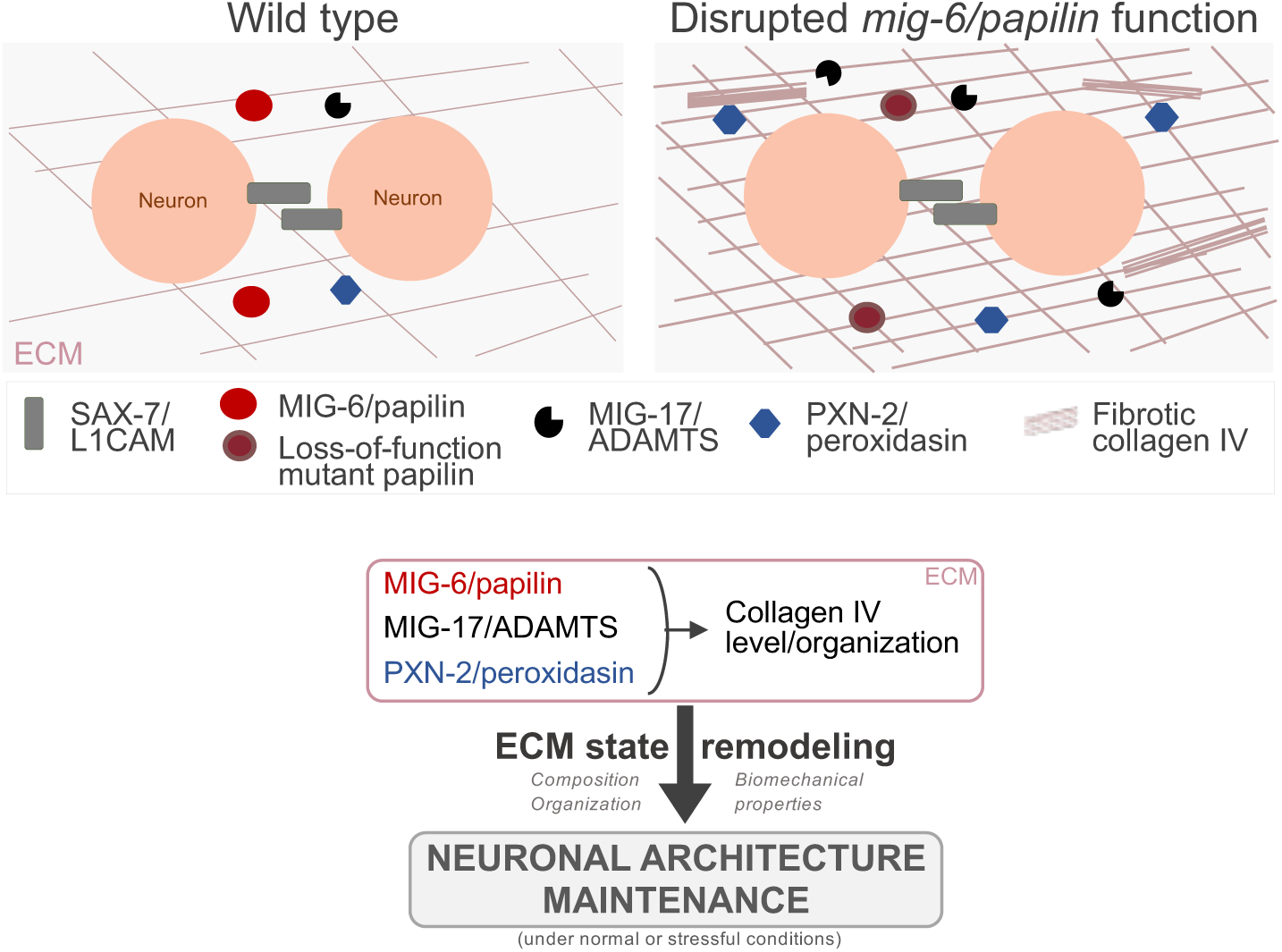
A model for the role of MIG-6S/papilin in collagen IV remodeling and neuronal maintenance. Summary of the cooperative functions of conserved ECM regulators MIG-6S/papilin, MIG-17/ADAMTS, and PXN-2/peroxidasin in the modulation of collagen IV levels and organization, impacting the long-term maintenance of neuronal architecture.

MIG-6/papilin is expressed throughout life, from embryogenesis to adulthood ^21,36,40,57,58^, in dynamic patterns that may reflect its requirements at the different life stages. Papilin indeed plays key developmental roles: the lack of papilin results in embryonic lethality in null mutant worms and in RNAi-depleted flies ^36,40^. Papilin is also required for organogenesis in flies ^36^, distal tip cell migration of the developing *C. elegans* gonad ^40^, as well as for enlargement of the gonad and the pharynx during *C. elegans*’ growth ^21,44^. A role for papilin in the nervous system has remained largely unexplored. There is one study in *C. elegans* showing that papilin participates axon guidance of the neuron ALA, which impacts primary dendrite development of the neuron PVD ^46^. In *Drosophila*, a recent report on a screen for regulators of central nervous system morphology mentions a papilin mutant found to have a misshapen central nervous system ^45^, which awaits further analysis. In our study, we isolated the mutation *mig-6(qv33)* in a screen for animals that suppress the age-progressive disorganization of *sax-7*/L1CAM mutants. Similar to other *mig-6* mutations, *mig-6(qv33)* mutant animals have gonad abnormalities, but otherwise display normal body morphology (i.e., normal musculature, pharynx, epidermis, and overall neuronal architecture). We uncovered a post-developmental role of *mig-6* in maintaining the positioning of neurons ASH and ASI, since depletion of *mig-6* function by RNAi treatment initiated from the first larval stage onwards suppressed the ASH and ASI neurons’ defects that progressively accumulate in *sax-7* null mutants, after having normally developed during embryogenesis.

*mig-6/papilin* encodes two isoforms, and our allelic series analysis and rescue assays demonstrate that the short isoform, MIG-6S, is active in neuronal maintenance, while MIG-6L is dispensable in this context. Isoform-specific roles of *mig-6* have been previously described in the gonad development ^40^ and ALA-PVD neuronal patterning ^46^. MIG-6S belongs to the ADAMTS-like ECM protein family, and consists of several domains, including the papilin cassette with its TSP1 repeats and the ADAMTS spacer, followed by numerous cysteine-rich lagrin repeats, and Kunitz domains. Our genetic and protein-domain analyses, combined with molecular predictions, point to the papilin cassette and the closest lagrin repeats as being critical for MIG-6S function in neuronal maintenance. Indeed, disruption of *mig-6S* function by *mig-6(qv33)* or other mutations affecting the papilin cassette/adjacent lagrin repeats suppressed the age-progressive neuronal disorganization of *sax-7/L1CAM* mutants, but *mig-6(sa580)* that affects a different lagrin repeat did not impact the neuronal maintenance. Allele-specific effects of mutations affecting this region of the protein have also been described in the context of pharynx growth, where mutants *mig-6(sa580)* do display a twisted pharynx, but the mutations *mig-6(k177, ev700,* or *ev701)*, which like *mig-6(qv33)* affect residues in more N-terminally located TSP1 domains or lagrin repeats, do not affect pharynx development ^44^. These allele-specific defects likely reflects the complexity of interactions of this multidomain MIG-6/papilin. Interestingly, *mig-6(qv33)* is a semi-dominant allele in neuronal maintenance, indicating that a minimum level of MIG-6/Papilin is required for proper function in this context. The *mig-6* locus was similarly described as haploinsufficient in gonad and ALA-PVD neuron development ^40,46^.

In both *C. elegans* and *Drosophila*, papilin is expressed by cells in charge of producing the ECM/basement membrane components, such as body wall muscles and epidermis in the worm, and hemocytes in flies. Indeed, we found that expression of MIG-6S from body wall muscles rescued its function in neuronal maintenance. Interestingly, MIG-6S/papilin itself localizes to the basement membrane of several organs, including the gonad, the pharynx, the intestine ^21,40^, as well as nerve structures (e.g., nerve tracts ^46^) in *C. elegans* larvae and adults. Similarly, in *Drosophila*, papilin localizes to the basement membranes, including those enveloping the central and peripheral nervous system of both larvae and adults ^36^. Interestingly, publicly available data shows that papilin is also expressed in central nervous system of adult mice (Allen Atlas, https://portal.brain-map.org/), suggesting papilin may function in the adult mammalian brain as well.

How might the basement membrane protein MIG-6/papilin regulate neuronal maintenance? The extent to which ECM remodeling determines the long-term preservation of the neuronal architecture laid out earlier in development is only beginning to be probed. We report that the role of the extracellular papilin MIG-6 and the ADAMTS protease MIG-17 in maintaining neuronal organization in *C. elegans* is through their cooperative function in regulating collagen IV. Papilin is a component of basement membranes but appears to have essential roles in their assembly or maintenance, since all of the *mig-6* mutants analyzed and *mig-6*(RNAi) treated animals display continuous basement membranes surrounding the pharynx, body wall muscles and the gonad, similar to wild type (this study, ^21^). Rather, papilin appears to affect specifically the remodeling of basement membranes, as disruption of MIG-6/papilin results in a dramatic build-up of extracellular collagen IV, a major component of ECM/basement membranes, indicating that MIG-6/papilin regulates collagen IV removal and distribution in the ECM. In *mig-6S* mutants and *mig-6*(RNAi) depleted animals, collagen IV accumulation is visible during larval stages and increases with age (both in terms of penetrance and of the extent of collagen build-up per animal), including within given individual animals as shown by our longitudinal analysis. Moreover, we observed an increase in intracellular collagen IV levels in body wall muscles, which produce both ECM components and MIG-6/papilin, suggesting that MIG-6/papilin may also impact collagen IV synthesis or degradation. RNAi-depletion of *mig-6* also results in collagen IV accumulation in the gonadal basement membrane ^21^.

MIG-6/papilin is an ADAMTS-like protein, sharing structural similarities with ADAMTS secreted ECM metalloproteinases, but lacking a catalytic domain. Its biochemical function is unclear, but *Drosophila* papilin can bind a procollagen N-proteinase ADAMTS *in vitro*, inhibiting its activity non-competitively, without directly interfering with the enzyme’s catalytic site ^36^. The papilin cassette alone could also inhibit the procollagen N-proteinase. The ‘papilin cassette’ in the papilin ADAMTS-like proteins is important for binding to ECM ^89^, and papilin domains often interact with ADAMTS proteases also containing a papilin cassette ^36^, further supporting a regulatory role of this key region in ECM remodeling. MIG-17 is an atypical ADAMTS enzyme as it lacks TSP1 domains and thus a papilin cassette; yet MIG-17 is classified as belonging to the ADAMTS family based on the other significant structural similarities to ADAMTS proteins ^54,90^. Our genetic and molecular analyses revealed that ADAMTS-like protein MIG-6/papilin and ADAMTS metalloproteinase MIG-17 function within the same pathway to regulate ECM remodelling. Indeed, (i) the loss of *mig-17* mirrors the *mig-6S* loss-of-function phenotype of extracellular collagen IV build up; (ii) the simultaneous loss of both genes in double homozygous mutant animals does not enhance the collagen IV fibrotic phenotype, and (iii) loosing half of the function of both genes in double heterozygous animals strongly enhances the defects. Although they function together to promote removal of extracellular collagen IV, *mig-17* and *mig-6S* mutants do have phenotypic differences, notably relating to pharynx development; also, *mig-6S* mutants display a higher level of intracellular collagen IV in muscle cells, while *mig-17* mutants do not. As a note, another genetic lesion of *mig-17*, *ola226*, was reported to have extracellular collagen IV accumulation in the head region ^51^. In sum, our data support the notion that MIG-6/papilin and MIG-17/ADAMTS functionally cooperate to regulate collagen IV remodeling. MIG-17/ADAMTS has been hypothesized to possess proteolytic activity toward collagen IV, based on studies both in *C. elegans* and *Drosophila* ^74,75,91^. It is thus conceivable that the build-up of collagen IV in the ECM of *mig-6* mutants could result from MIG-17 being inhibited or less efficient, especially since we found that MIG-17 levels are increased in the head region of *mig-6* mutant animals. We directly tested this by overexpression of functional MIG-17/ADAMTS, which did not reverse the collagen IV fibrotic phenotype in *mig-6* mutants, suggesting that MIG-17/ADAMTS is in its active form in *mig-6S* mutants yet unable to degrade collagen IV, perhaps due to its high degree of crosslinking ^92–94^. In this scenario, MIG-6 regulates the level and activity of an ADAMTS through its impact on the ECM state.

Interestingly, we show that MIG-6/papilin influences the levels and distribution of MIG-17/ADAMTS and of the extracellular collagen IV crosslinking enzyme PXN-2/peroxidasin in the vicinity of the affected neuronal structures. Other studies have also documented that MIG-6/papilin affects the distribution of MIG-17 in the basement membrane of the developing gonad ^40^, and *mig-6* depletion by RNAi increased the levels of the ADAMTS proteinases MIG-17 and GON-1, and of PXN-2/peroxidasin-2 in the gonadal basement membrane ^21^. Whether this involves a physical interaction (direct or indirect) between MIG-6/papilin and these proteins is to be determined. Regardless, these observations together suggest that papilin might play a broad role in collagen IV/ECM remodeling. Importantly, we show that loss of *mig-6S* or loss of *mig-17* profoundly suppresses the neuronal maintenance defects that occur in *sax-7/L1CAM* mutants. Furthermore, losing the function of both MIG-6 and MIG-17 in homozygous double mutant animals did not enhance the suppression of the neuronal maintenance defects of *sax-7* mutants, and loss of one copy of each gene in double heterozygous animals significantly enhanced the suppression compared to each heterozygous single mutant. These observations are consistent with the notion that MIG-6 and MIG-17 function in the same pathway to impact the maintenance of neuronal architecture. Evidence for a functional relationship between MIG-6 and MIG-17 exists also in the context of the *C. elegans* developing gonad, where *mig-6* and *mig-17* genetically interact ^40^. Importantly, MIG-17 is not involved in ALA-PVD neurons patterning, indicating that MIG-6/Papilin operates through distinct mechanisms depending on the biological context, which is consistent with the specificity of defects displayed by different alleles affecting *mig-6S*, possibly interacting with distinct functional partners through distinct regions of this multidomain protein.

At the level of multicellular neuronal structures, such as ganglia or nerve cords, a delicate balance must exist between ECM stability, which preserves the architecture of the existing neuronal structures, and ECM remodeling, which accommodates growth of the neuronal structures during post-natal life, as well as adapting to shape changes that accompany the animal’s movements. The shared fibrotic collagen IV phenotype between *mig-6S* and *mig-17* mutants suggests that the altered state of collagen IV in these two mutants contributes to their ability to sustain neuronal architecture in *sax-7* mutants. An excess of crosslinked collagen IV may reinforce the integrity of the basement membrane, thereby supporting the maintenance of neuronal organization. We favor a model in which enhanced basement membrane integrity leads to maintained neuronal architecture for several reasons. First, the *mig-6* and *mig-17* mutations that do suppress *sax-7*-neuronal disorganization display a dramatic accumulation of extracellular collagen IV. Second, both collagen IV abundance and its crosslinking are required for neuronal maintenance, as reducing collagen IV by *emb-9*(RNAi) significantly reversed the stabilizing effect brought about loss of MIG-6S or of MIG-17, as does reducing the crosslinking of collagen IV by RNAi knockdown of PXN-2/peroxidasin ^95^. Collagen IV was also key in the role of MIG-6/papilin in modulating the response of neuronal architecture to heightened mechanical stress, as loss of *mig-6* was protective of head ganglia organization in animals subjected to swimming which leads to increased mechanical stress on the nervous system, due to the constant and rapid swimming muscle contractions was also dependent on collagen IV levels. Collectively, these findings underscore that extracellular collagen IV networks are key in neuronal maintenance. The fibrotic-like structures displayed by *mig-6* and *mig-17* mutants are unlikely directly involved in stabilizing neuronal architecture; rather, these fibrotic accumulations are the most obvious manifestations of dysregulated ECM remodelling, which also affects the basement membranes surrounding neuronal structures in *mig-6* and *mig-17* mutants.

Collagen IV, thanks to its unique ability to form intermolecular covalent bonds, provides the basement membrane with the capacity to withstand mechanical stress ^76,96^. Thus, we characterized the mechanical properties that result from the loss of functional MIG-6/papilin, more specifically, by analyzing the high-frequency longitudinal modulus of tissues, using Brillouin microscopy ^71^. We imaged an area neighboring the neurons under study, and compared mechanical properties of *mig-6* mutants and wild-type animals at two ages, earlier in larval life, and just before becoming adults. The area imaged, located in the posterior region of the animal’s head, comprises several cell types, including neurons, muscles, glia, pharynx, and their ECM. While the contribution of each individual adjacent cell and of the local ECMs to the measured mechanical proprieties cannot be reliably discriminated in intact animals with the current resolution of the Brillouin microscope, ECM including basement membranes, is known to exert a key influence on tissue biomechanics ^97–99^. Thus, tissue viscoelastic properties are significantly determined by the ECM ^97,99^. Our Brillouin spectral analysis revealed that loss of MIG-6/papilin results in altered biomechanical properties in the head region which houses the neuronal ganglia we primarily analyze. Collectively, the imaged tissues and associated ECMs in *mig-6* mutants have reduced viscosity and elasticity, indicating impaired viscoelastic properties. Importantly, cellular viscoelasticity is a regulator of cell behavior, associated with both physiological and pathological states across species ^97,99^. Thus, having been able to capture changes that inform on the viscoelastic properties of animals lacking MIG-6/papilin is a key finding, especially since few such *in vivo* measurements have been achieved to date ^88,100^.

The viscosity of a substrate is known to influence cell migration, with cells from normal tissue and tumor cells both exhibiting increased migration speed on highly viscous substrates or extracellular fluids ^101^ ^{Bera, 2022^ ^#723 102^. Also, both higher and lower cellular elasticity are linked to the motility of cancer cells ^103–105^. Conversely, cell adhesion can occur on the surface of low viscosity liquids ^106,107^. Thus, the decreased viscosity of *mig-6* mutants may somehow, possibly via distinct cell-ECM interactions, result in enhanced cell adhesion enabling neurons to maintain their normal architecture. In addition, *mig-6* mutants present a decrease in loss tangent that translates into decreased viscoelasticity, suggesting that their tissues exhibit more solid-like properties with reduced energy dissipation ^73^. Tissues and matrix mechanics are sensed by cells and converted into chemical signals through mechanotransduction ^97,108^. Thus, the decreased viscoelasticty in *mig-6* mutants, and the proposed associated reduction in energy dissipation, could modulate mechanosensing and regulate cellular responses ^109,110^, to better preserve tissue shape and maintain neuronal architecture. Indeed, this may be related to the altered collagen IV levels, organization, and remodeling that we uncovered in *mig-6* mutants, which could profoundly impact the overall ECM composition and organization. A parallel could be drawn with the excessive production of ECM components in tissue fibrosis, which results in decreased viscoelasticity ^108^. Such fibrotic states also lead to progressive matrix stiffening ^108^. The build-up of collagen IV occurring in *mig-6* mutants, and that depleting the crosslinking enzyme peroxidasin/PXN-2 attenuated their fibrotic state, suggests that collagen IV molecules in *mig-6* mutants have increased covalent sulfilimine cross-links, which could lead to increased ECM stiffness ^81,111^, and consequently, a likely decreased flexibility. Given the expected increased stiffness in *mig-6* mutants and the similar altered remodeling of ECM collagen IV in double *sax-7; mig-6* mutant animals, the mechanical proprieties arising from loss of *mig-6* could help maintain neuronal architecture through increased stiffness.

Overall, we propose that the animal’s biomechanical changes resulting from the loss of MIG-6/papilin are linked to their altered ECM state. The differences in biomechanical properties are likely to bring about changes in ECM-neuron interactions, and/or in the state of neurons, such that neuronal architecture is preserved, even in the absence of SAX-7/L1CAM, or in conditions of heightened physical stress from the incessant muscle contractions of continuous swimming. Future studies could further elucidate the underpinnings of this remarkable state resulting from changes in ECM remodeling by the conserved ECM regulator MIG-6/papilin, which safeguards neuronal architecture during post-natal life and into adulthood.

Interestingly, while MIG-6/papilin plays a crucial role in defining the state of the ECM, its effects are specific to the precise molecular landscapes in distinct neuronal structures of the animal. For instance, we found that whereas loss of *mig-6* suppresses the maintenance defects of axon position in the ventral nerve cord that are caused by the loss of adhesion molecule SAX-7/L1CAM, or by loss of basement membrane protein DIG-1, it fails to suppress similar maintenance defects of the same axons when caused by the loss of two-Ig domain proteins ZIG-3 and ZIG-4. Similarly, whereas loss of *mig-6* suppresses the head ganglia defects in both *sax-7/L1CAM* and *dig-1* mutants, it had no effect on tail ganglia maintenance defects displayed by *dig-1* mutants. These observations underscore the complexity of the molecular interactions involving distinct ECM networks that surround different neuronal assemblies. The specificity of MIG-6/papilin’s action is also evident in the different developmental consequences of *mig-6* mutations across different contexts, including the gonad, the pharynx, and the ALA-PVD neurons in the lateral nerve tract. This specificity is also reflected in its functional interaction with MIG-17/ADAMTS, which affects head ganglia maintenance and distal tip cell migration (this work,^40^), but not for ALA-PVD neuronal patterning ^46^. Future studies will help elucidate the interactions among other ECM components that may participate in the remodeling process orchestrated by papilin.

The extracellular matrix (ECM) has emerged as a key regulator of nervous system development and maintenance across diverse species ^112–114^. In *C. elegans*, the ECM modulates synaptic development ^115^, as well as synaptic maintenance, with collagen IV and metalloproteinase GON-1 being implicated in sustaining synapse morphology of neuromuscular junctions ^116^ ^117^. In the mammalian central nervous system, the ECM is a large part of the neural tissue and serves various functions ranging from supporting cell migration, to regulating synaptic transmission and plasticity, to actively modulating the neural tissue after injury. In particular, the perineuronal nets (PNNs), a specialized form of ECM surrounding dendritic spines, have been shown to be dynamically regulated, impacting both structural and functional plasticity ^118^. The ECM composition of PNNs is regulated by the expression of proteases that target distinct PNN, enabling the transition from states of plasticity to stability ^119^. Disruptions in PNN composition is linked to neurodegenerative diseases ^13,14^. Collagen IV is a well-conserved component of basement membranes, including in the vertebrate central nervous system. It is conceivable that functional interactions between papilin, ADAMTS metalloproteinases and ECM molecules, similar to those described in this study, may also occur in mammals to maintain neuronal architecture throughout life.

Neuronal structures need to withstand deformations caused by the animal’s growth and body movements to prevent structural damage to neural circuits. How multicellular neuronal assemblies endure mechanical stress to sustain their architecture on the long term remains poorly understood. This work provides a mechanism by which the regulation of ECM remodeling enables and supports the maintenance of neuronal architecture postnatally and into adulthood. Other mechanisms previously described to critically impact the maintenance of neuronal architecture also rely on non-cell-autonomous biological functions. For instance, the secreted immunoglobulin proteins ZIG-3 and ZIG-4 are thought to stabilize axons positioning by modulating inter-axon adhesive properties ^30,31^. The cell adhesion molecule SAX-7/L1CAM mediates cell surface homophilic and heterophilic interactions between neurons and its neighboring cells (e.g, other neurons or epidermal) ^25,27,46^. The secreted basement membrane protein DIG-1 is proposed to bridge interactions between the basement membranes ensheathing neuronal structures and adjoining muscle cells ^28^. Collagen IV and ADAMTS/GON-1 ensures the maintenance of synaptic morphology at the neuromuscular junction ^116,117^. MIG-17/ADAMTS maintains synapse location and morphology during post-embryonic growth by modulating muscle basement membrane, which impacts interactions between epidermis, glia, and the associated synapses ^51,120^. The two-immunoglobulin domain protein ZIG-10 expressed on the epidermis underlying the nerve cord maintains synaptic density as the animal grows ^121^. More recently, the interplay between epithelial cells, their ECM, cell junctions and glial cells was shown to ensure the preservation of glia morphology in the face of environmental challenges, which in turn protects the associated neuron’s shape and function ^88,122^. Finally, cytoskeletal components too can act non-cell-autonomously from the underlying epidermis embedding the axon of a neuron to preserve its integrity ^123.^. Thus, the combined actions of both intrinsic and extrinsic mechanisms safeguard the intricate multicellular structures of the nervous system. Understanding general principles governing the long-term maintenance of the neuronal architectures underlying neural circuits is crucial for elucidating the bases of neurodegenerative conditions.

## Acknowledgements

We thank members of the Bénard laboratory for advice throughout this study and comments on the manuscript; Mark Alkema, Michael Francis, Kota Mizumoto, Nicolas Pilon, Saïd Kourrich, and Jean-Claude Labbé for stimulating discussions; Grégoire Bonnamour and Denis Flipo (UQAM) for confocal microscopy expertise; several laboratories for sharing strains and/or plasmids, including J. Culotti, D. Sherwood, A. Fire, and S. Hekimi; Wormbase (www.wormbase.org) provided information about genome sequence and annotations; the *Caenorhabditis* Genetics Center, which is funded by NIH Office of Research Infrastructure Programs (P40 OD010440) for strains; WormAtlas and WormImage, which are funded by NIH OD010943 to David H. Hall; John White and Jonathan Hodgkin for donation of the MRC/LMB archives to the D. Hall laboratory for curation.

## Supplementary Materials

### MATERIALS AND METHODS

#### *C. elegans* strains and genetics

Strains were cultured at 20°C on nematode growth medium (NGM) agar plates seeded with OP50 bacteria as described ^1^, unless otherwise specified. N2 is the reference wild-type strain. Mutant alleles and reporter strains were outcrossed at least three times prior to strain generation or analysis (listed in **Table S1**). Strains were constructed using standard genetic procedures and are listed in **Table S1** and **Table S2**. Genotypes were confirmed by visible phenotypes when possible, and by PCR or sequencing (primers listed in **Table S3**).

#### EMS forward genetic screen

We carried out a forward F2 clonal genetic screen for modifiers of neuronal maintenance defects using VQ90 *sax-7(qv24); zig-5(ok1065) zig-8(ok561); glp-4(bn2ts); oyIs14* as our screening strain, which was maintained at 15°C. *glp-4(bn2ts)* was used for efficient examination of adult worms, as it is a temperature-sensitive sterile mutation when shifted to 25°C ^2^. Allele *qv24* of *sax-*7 will be described elsewhere (C.R.B and C.Y.B., unpub. results). For each mutagenesis round, VQ90 L4 worms were mutagenized with 25 µM ethyl methanesulfonate and allowed to recover at 15°C. Following recovery, five P_0_ mutagenized worms were plated onto five plates (25 P_0_s in total) and put at 15°C to lay their broods. 50 F1s from each P_0_ plate were picked singly as L4 or young adults. From each of the 250 F1s, seven F2s were picked clonally (1750 F2s per round of mutagenesis) and grown at 15°C for two days (plates containing singled F1s were kept at 15°C for future use). After two days, the plates where single F2 animals had been picked were shifted to 25°C. When the F3 animals were 5-day-old adults or older, they were examined under a Zeiss V8 Discovery fluorescence stereoscope to identify rare broods exhibiting a modified phenotype compared to VQ90 control. Upon identification of a candidate modifier mutation, single animals were picked clonally from the corresponding F1 plate (kept at 15°C) to re-isolate the modifier, and their progeny were mounted on slides to examine and quantify the phenotype using an upright Zeiss Axioskop 2 Plus fluorescence microscope. If the modifier mutation was re-isolated and exhibited a robust phenotype, we outcrossed it. For outcrosses, we used males from the non-mutagenized strain VQ47 *sax-7(qv24); zig-5(ok1065) zig-8(ok561); hdIs29*, which is similar to the one used in the screen, but carries a red fluorescent marker to readily identify cross progeny. We thus could determine if the new modifier mutation was heritable, and at the same time remove *glp-4(bn2ts)*, and outcross other irrelevant EMS-induced mutations from the genetic background. Subsequent outcrosses with wild-type N2 males ensured that all strains used are wild type for *zig-5, zig-8,* and *glp-4*.

#### Isolation and molecular identification of *qv18*

*qv18* was isolated as a suppressor of the *sax7(qv24)* neuronal maintenance defects in the genetic screen described above. Outcrosses of the new strain bearing the suppressor mutation *qv18* (as described above) showed that the suppressor was heritable, penetrant, and likely monogenic. We also noticed abnormal segregation ratios, consistent with a possible chromosomal rearrangement resulting from mutagenesis; we therefore later engineered the *qv18* causal suppressor mutation in a clean genetic background. To identify the causal mutation in *qv18*, we performed whole genome sequencing on the pooled DNA of 18 independent suppressor lines (each line derived from one of 18 independent F2s from the outcross, with careful re-homozygosing of the suppression of the neuronal maintenance defect confirmed over 3 generations ^3^. The sequencing data were uploaded to the public server at usegalaxy.org and we used the Cloudmap pipeline to analyze the data ^4^, which pointed to a *qv18* linkage region on chromosome V. A candidate variant therein was a mutation in the gene *mig-6*, which we pursued further by (i) performing rescue experiments, (ii) introducing the candidate mutation by genome editing in *sax-7* mutants with an otherwise wild-type *mig-6* gene (which yielded *qv33*), (iii) analyzing the effect of loss of *mig-6* function by *mig-6*(RNAi) experiments, and (iv) examining other *mig-6* mutant alleles.

#### Generation of *qv33* by CRISPR-Cas9

To confirm the molecular identity of the causal mutation suppressing *sax-7* defects in the *qv18* suppressor strain, the candidate *mig-6* mutation was reintroduced in a clean *sax-7(qv30); oyIs14* genetic background using CRISPR-Cas9 genome editing. A target sequence against *mig-6* was selected (ggtaccgctgaatgtggtgg) and cloned to generate a gRNA plasmid (pPP30). A 100 nt oligo carrying the candidate *qv18* missense mutation, at nucleotide 3409 of cosmid C37C3, served as the repair template (oCB1599 acatggtacacatcttcatggtccgagtgtaccgctgaatgtggtggtgAatcccaagatcgtgtcgctgtttgcttgaactacgataagaagccagttc, IDT synthesized). A DNA mix containing the Cas9 plasmid (pDD162), the gRNA plasmid pJA58 (*dpy-10* target) and *dpy-10(cn64)* ssDNA repair template for CRISPR co-conversion, as well as the *qv18* gRNA plasmid pPP30 and repair template oCB1599 was injected as described ^5,6^. Rol or Dpy F1 animals were picked clonally and their F2 progeny were screened by PCR (using primers listed in Table S1), followed by restriction enzyme digestion, as a BamHI site present in the wild-type sequence is disrupted by the *qv18/qv33* single nucleotide substitution.

The same CRISPR-Cas9 strategy was used to generate the double mutant *mig-6(qv33) mig-17(k174)*, by introducing *qv33* into the *mig-17(k174)* background, as the loci are 2 cM apart on chromosome V.

#### Prediction and analysis of the three-dimensional structure of MIG-6S

The three-dimensional structure of MIG-6S (Uniprot accession O76840-2) was predicted using ColabFold 1.5.2 locally installed on the servers of the Digital Research Alliance of Canada. This version of ColabFold uses AlphaFold 2.3.1 and MMSeqs2 14-7e284. The depicted structure corresponds to the model presenting the highest pLDDT values that was obtained using a recycle count of 12. The images of the structure were made using PyMOL 2.5.4. Sequence logos were generated using Weblogo 3.7.12 and an alignment of 250 protein sequences from Ecdysozoa species, including nematodes and arthropods obtained using MMSeqs2.

#### Visualization and quantification of neuroanatomy and EMB-9/collagen IV structures

1st larval stage (L1), 2^nd^ larval stage (L2), 4th larval stage animals (L4), or 2-day old adult (selected as late L4 and observed 48 hours later) nematodes were mounted on agarose pads, immobilized with 75 mM NaN_3_. These animals were observed under Nomarski or fluorescence microscopy Axio Scope.A1 or Axio Imager.M2 (Zeiss), with a 40x objective or a 100x oil immersion objective (for ventral nerve cord examination). Images were acquired using an AxioCam camera (Zeiss) and processed using ZEN (Zeiss).

##### Analysis of ASH and ASI

The cell bodies of ASH/ASI and their axons located in the nerve ring were visualized in L2, L4, and 2-day old adults, using reporter P*sra-6::DsRed2* (*hdIs26*). In the wild type, the ASHL/R and the ASIL/R soma are located posterior to the nerve ring. An animal was counted as mutant when along the antero-posterior axis of the animal, at least one of the ASH/ASI soma was not posterior to the nerve ring (but either anterior to or flanking the nerve ring).

##### Analysis of PVQ

The axons of neurons PVQL/R in the ventral nerve cord were visualized using P*sra-6::DsRed2* (*hdIs26*), in freshly hatched L1 and in L4 larvae, as described ^7^. Briefly, in the wildtype, the axon of PVQL is positioned along the left fascicle of the ventral nerve cord, whereas the axon of PVQR is in the right fascicle. Animals were counted as having an axon flip-over defect when an axon was flipped to the opposite fascicle at any point along the ventral nerve cord.

##### Analysis of collagen IV fibrotic-like structures

We consider as fibrotic collagen IV the elongated structures or enrichments of EMB-9::mCherry signal (also reporters EMB-9::Dendra or EMB-9::mNG) that are observed in the posterior head region of *mig-6* or *mig-17* mutants (see **Figs. 4 and 6**). These structures are very rarely observed in the wild type, and they are different from the wild-type pattern of collagen IV signal present along the body wall muscles basal lamina (collagen IV concentrates in the basal lamina underneath each dense body and M line of sarcomeric muscles, including in the head region). To unequivocally assess the presence of collagen IV fibrotic-like structures in each animal, z-stacks were captured, and all the z-planes images were examined (the experimenter was blinded for genotype). Collagen IV fibrotic structures vary in number, length, and position (see **Fig. 4B, D, F**); animals displaying at least one such structure were counted as having fibrotic collagen IV.

As a result of the *pxn-2*(RNAi) (**Fig. 7F-G**), a different collagen IV pattern occurred, where the fibrotic-like structures appeared ’fragmented’. We consider a fibrotic like-structure to be fragmented if accumulation of collagen IV appeared as a series of elongated puncta, which were connected to one another by a thin line of collagen IV.

#### Microscopy and fluorescence intensity quantification

Animals were observed at precise ages, namely "young adults" (just molted from the L4 stage, <3 hours post L4 molt) and "day 2 adults" (48 hours after the young adult stage), mounted on 5% agarose pads with 75 mM NaN_3_. Images were acquired as a z-stack using a Plan Apo 40x/0.95 NA objective on a Zeiss Axio Imager.M2 (equipped with an AxioCam camera and ZEN software). AutoQuant X deconvolution software was used to remove blur and enhance contrast and resolution. A Nikon A1 laser scanning confocal microscope (equipped with an EMCCD camera and NIS elements software) was also used for image acquisition and analysis, using either a Plan Apo λ 40x/0.95 NA or Plan Apo λ 60x/1.4 NA oil immersion objective.

EMB-9::mCherry, MIG-17::GFP, and mNG::PXN-2 total fluorescence intensities in z-projections were quantified using ImageJ. All image acquisition parameters (including exposure time, excitation intensity, and gain), for each imaging channel, were fixed across genotypes for a given fluorescent reporter. The region of interest (ROI) containing mCherry, GFP or mNG signals on the images was either outlined manually (the whole head) or defined as a rectangle. To correct for background fluorescence, four ROIs located outside but adjacent to the animal’s head were used; the mean fluorescence of background ROIs was subtracted from the signal intensity for the same image, using the following formula: Corrected IntDen = IntDen of experimental ROI - (mean of background ROI X area of experimental ROI). 3D isosurface renderings of EMB-9::mCherry in *mig-6* mutants were constructed using Imaris 7.4 software (Bitplane).

#### Brillouin microscopy and imaging of mechanical properties

Brillouin microscopy measures tissue mechanical properties (elasticity, viscosity) in the GHz frequency range through the interaction of light with the sample’s acoustic phonons ^8,9^. The shift and linewidth of the Brillouin scattered light spectrum gives information about the longitudinal modulus, which is directly related to the elastic and viscous modulus of the material, respectively. Brillouin imaging was performed using a Brillouin microscope previously described ^10^, and animal preparation and imaging protocols previously described ^8^. Briefly, a commercial Zeiss body (Axiovert 200 M) coupled with a home-built spectrometer based on a 2-VIPA configuration, provides a precision of 22 and 56 MHz for Brillouin shift and linewidth measurements, respectively, for our measurement parameters (100 ms exposure time, 4 mW optical power on the sample) and a measured spectral resolution of 520 MHz. To account for the finite spectral resolution, the latter (520 MHz) was subtracted from the measured linewidth values (subtraction corresponds to deconvolution, as detailed in Chan et al 2021). The effective NA of the objective (0.85) leads to ∼100–110 MHz downshift in the Brillouin shift. A 532 nm laser (Torus, Laser Quantum) was used for Brillouin imaging. Wild-type and *mig-6* mutant animals of the L2 and L4 stages were imaged. Animals were grown in the absence of food for 2 h before experiments, to minimize variability of Brillouin signal in the pharynx due to the presence of bacteria food. Animals were anesthetized using M9 buffer containing 10 mM NaN_3_ and mounted on 2% agarose pads containing 10 mM NaN_3_. Brillouin images were acquired with a 40x/1.0 NA Zeiss objective and an integration time for a single point of 100 ms. The optical power on the sample was kept below 4 mW and no apparent photodamage was observed after imaging.

For Brillouin image analysis, only images of healthy animals (with no vacuoles around the bulb) were used. Also, we only analyzed images of animals that had moved very little, specifically <1.5 µm for L2 and <2.2 µm for L4 animals (the brightfield imaging acquisition preceding and following Brillouin image acquisition enabled us to measure any animal movement during imaging). Regions of interest for analysis of Brillouin microscopy images were drawn flanking the 2nd pharyngeal bulb to the animal’s body edge (cuticle). Spatial maps of elasticity and viscosity are plotted from the acquired data, using function “Measure” and Lookup table mpI-Inferno in Fiji and adjusting the “Brightfield and Contrast” at 7.5–8.2 or 1.02–2.05 respectively. From the raw Brillouin shift and linewidth values we compute the Brillouin elastic and viscous contrast, analogous to Antonacci et al. 2020. Mean Brillouin linewidth, shift, and loss tangent of the ROIs were measured for each animal, averaged for each genotype, and compared between genotypes. Graphs and statistical analyses were made on raw and relative values. Relative values consist of raw values divided by the mean value obtained for the wild-type genotype.

Refractive index of animals of each genotype was measured by performing label-free 3D Holo-Tomographic Live Cell Imaging, using the Nanolive 3D Cell Explorer Fluo, as previously described ^8^. Briefly, L2 or L4 animals were mounted on slides using Pluronic F127 36% w/v + 1 mM tetramisole solution. Brightfield images were acquired using a 60x/0.8 NA objective using a FITC filter of the Nanolive Module LED DAPI-FITC-TRITC/Cy5 4X B 000.

#### Transgenes and rescue experiments

##### *mig-6S* minigene (pCB411)

The *mig-6S* minigene includes exons 1-11, introns 1-4 of isoform a, and the endogenous *mig-6S* 3’UTR, which were amplified out of pZH125 ^11^ using primers oCBQc17 (CAAGCTCCCGGGATGAGGTTGCTGCTCTTCTCGG) and oCBQc18 (CATGATACTAGTGCGCAACAATGGGTGAAGAAAGC), adding XmaI and SpeI sites. This insert was cloned into vector PCR2.1 TOPO.

##### Prgef-1::mig-6S (pCB408)

Vector backbone and promoter were obtained from plasmid pCB199 (P*rgef-1::rib-1*cDNA) by digesting with XmaI [3480] and SpeI [5422] to remove *rib-1*; the *mig-6S* insert [5625 nt] released from p 411 by XmaI and SpeI digestion was ligated.

##### P*dpy-7::mig-6S* (pCB409)

Vector backbone and promoter were obtained from plasmid pCB249 (P*dpy-7::lon-1a* cDNA) by digesting with XmaI [271] and SpeI [2003] to remove *lon-1*; the *mig-6S* insert [5625 nt] released from pCB411 by XmaI and SpeI digestion was ligated.

##### P*myo-3::mig-6S* (pCB416)

Plasmid pPD95.86 (P*myo-3::unc-54* 3’UTR) was digested with XbaI [2414] and ApaI [3319] to remove the *unc-54* 3’UTR. The released vector was ligated with the *mig-6S* insert [5643 nt], which was amplified from pZH125 using primers oCBQc44 (TCGGAGTCTAGAATGAGGTTGCTGCTCTTCTCGG) and oCBQc20 (CAAGATGGGCCCGCGCAACAATGGGTGAAGAAAGC) to add on XbaI and ApaI sites, and digested.

##### P*myo-3::mig-6SΔpapilin cassette/lagrin repeats* (pCB483)

Plasmid pCB416 (P*myo-3::mig-6S*) was digested using NcoI [2715] and ClaI [7527] to remove the sequences corresponding to the end of exon 2 (from the NcoI site) until exon 11 (up to ClaI site) of *mig-6S*. This fragment was ligated with a second one to generate a plasmid that lacks the sequences coding for the papilin cassette and the lagrin repeats, but included the rest of coding sequence of *mig-6* until the Stop codon and the 3’UTR as in pCB416. The second fragment consists of most of exon 8 and exons 9, 10 and 11, which was PCR amplified from plasmid pCB416 using primers oCB2266 (CATGATCCATGGccaacttgcgttgactctg, to add an NcoI site) and oCBQc25 (GAACTTACTCGGGCATCTCG, taking advantage of an internal ClaI site) and digested, to reconstitute an in-frame *mig-6SΔpapilin cassette* version by ligation.

##### P*myo-3::mig-6SΔKunitz domains* (pCB492)

A fragment of the *mig-6S* gene containing sequence from the initiator ATG in exon 1 all the way to exon 8, thus lacking sequence encoding the Kunitz inhibitor domains, was PCR amplified using primers oCBQc44 (tcggagtctagaatgaggttgctgctcttctcgg, containing an XbaI site) and oCB1810 (catgatAGCGCTTTAattacaggcggcgattttatg, to add an AfeI site) from plasmid pCB416 (P*myo-3::mig-6S*) and digested. This fragment was ligated with another one containing the vector backbone and promoter obtained from plasmid pCB423 (P*myo-3::sdn-1cDNA*) by digesting with XbaI and AfeI.

All inserts of finalized clones were verified by sequencing.

Transgenic animals were generated by standard microinjection techniques ^12^. Plasmid pZH125, which contains a *mig-6*S minigene expressed under the *mig-6* endogenous promoter ^11^, was injected at a concentration of 5 ng/μL. For tissue-specific rescue assays, pCB416 [*Pmyo-3::mig-6S*] was injected at 1 ng/μL with *ceh-22::gfp* or *lgc-11::gfp* as a marker of transgenesis; pCB408 [*Prgef-1::mig-6S*] was injected at 7 ng/μL with *lgc-11::gfp* as co-injection marker; and pCB409 [*Pdpy-7::mig-6S*] was injected at 0.5 ng/μL and 0.1 ng/μL with *lgc-11::gfp* as co-injection marker. Co-injection markers were injected at 50 ng/μL and pBSK(+) was added to each mix to reach a final DNA concentration of 200 ng/µL. DNA mixes were injected into a strain where the *mig-6* gene is balanced: *sax-7(qv30); mig-6(qv33)/dpy-11(e224) oyIs14*. Transgenic strains are listed in Table S4.

#### Liquid cultures

After hatching into L1 larvae, animals were grown in liquid culture for up to 5 days, kept under constant agitation on a nutator for them to swim continuously. Synchronized L1 larvae were obtained by treating gravid adults with a bleach solution ^13^, resuspending the collected embryos in 2 mL S medium in 15 mL conical tubes, and incubating them overnight at 20°C on a nutator to allow their hatching into L1 larvae that are arrested in the absence of food. The L1 larvae culture was then supplemented with 200 μL of concentrated *E. coli* OP50 and kept under constant agitation on a nutator at 20°C for 5 days, with additional concentrated *E. coli* OP50 added at days 1 and 3 post hatching. Animals were examined by microscopy at 3 and/or 5 days after hatching (post-L1 days 3 or 5 is equivalent to the age of young or 2-day-old adults of animals grown on solid plates, respectively).

#### RNA interference assays

RNAi experiments were performed by the feeding method using RNAi bacterial clones from the Ahringer RNAi library ^14,15^. Single colonies for the L4440 empty vector negative control, and for clones containing *emb-9* and *pxn-*2 sequences, were obtained on LB plates with ampicillin (75 μg/mL) and tetracycline (12.5 μg/mL). They were then grown in LB medium with ampicillin (75 μg/mL) for 16h at 37°C, to which 1 mM IPTG was then added and incubated for an additional hour to induce dsRNA expression. Concentrated cultures of these RNAi bacteria were seeded onto NGM plates (75 μg/mL Amp, 1 mM IPTG) and left to dry at room temperature for overnight induction. For *mig-6*(RNAi) experiments, *sax-7(qv30)* animals were placed on RNAi plates at the L2 stage and examined as day 2 adults (48 hours post L4). For *emb-9*(RNAi) and *pxn-2*(RNAi), synchronized L1 larvae ^13^ were distributed onto RNAi plates, incubated at 20°C, and examined as day 2 adults (48 hours post L4).

For RNAi assays done with animals swimming in liquid (**Fig. 7L**), worms were synchronized as L1 larvae and then grown in liquid media as described above, except that they were fed with HT115 RNAi bacteria (carrying the control vector or the *emb-9*(RNAi) plasmid) in S medium supplemented with ampicillin (75 μg/mL) and IPTG (1 mM). 250 μL of concentrated bacteria was added to the liquid cultures initially (after hatching), and then at day 3 post L1. Optimal induction of dsRNA expression by the RNAi bacteria was achieved by growing bacteria for 14 hours at 37°C, and then adding 5 mM IPTG for growth for another 4 hours. Animals were recovered from the liquid culture and examined at day 5 post L1.

#### Statistical analyses

All statistical analyses were performed using R (version 4.1.2) using the R Stats package (’stats’ version 4.4.2). Data are presented as mean ± standard error of proportion, or as mean ± standard error of the mean. As indicated in each Figure Legend, statistical tests were performed using z-test, unpaired two-tailed Student’s t-test, or one-way ANOVA when applicable (for parametric datasets). For non-parametric datasets, the Wilcoxon-Mann-Whitney test was used. Appropriate *post-hoc* tests were performed for multiple comparisons: Bonferroni correction was applied after z-tests and Wilcoxon-Mann-Whitney tests, while Tukey HSD correction was applied following ANOVA. All sample sizes and raw data are available in **Supplementary Information**.

---

**Figure S1.**
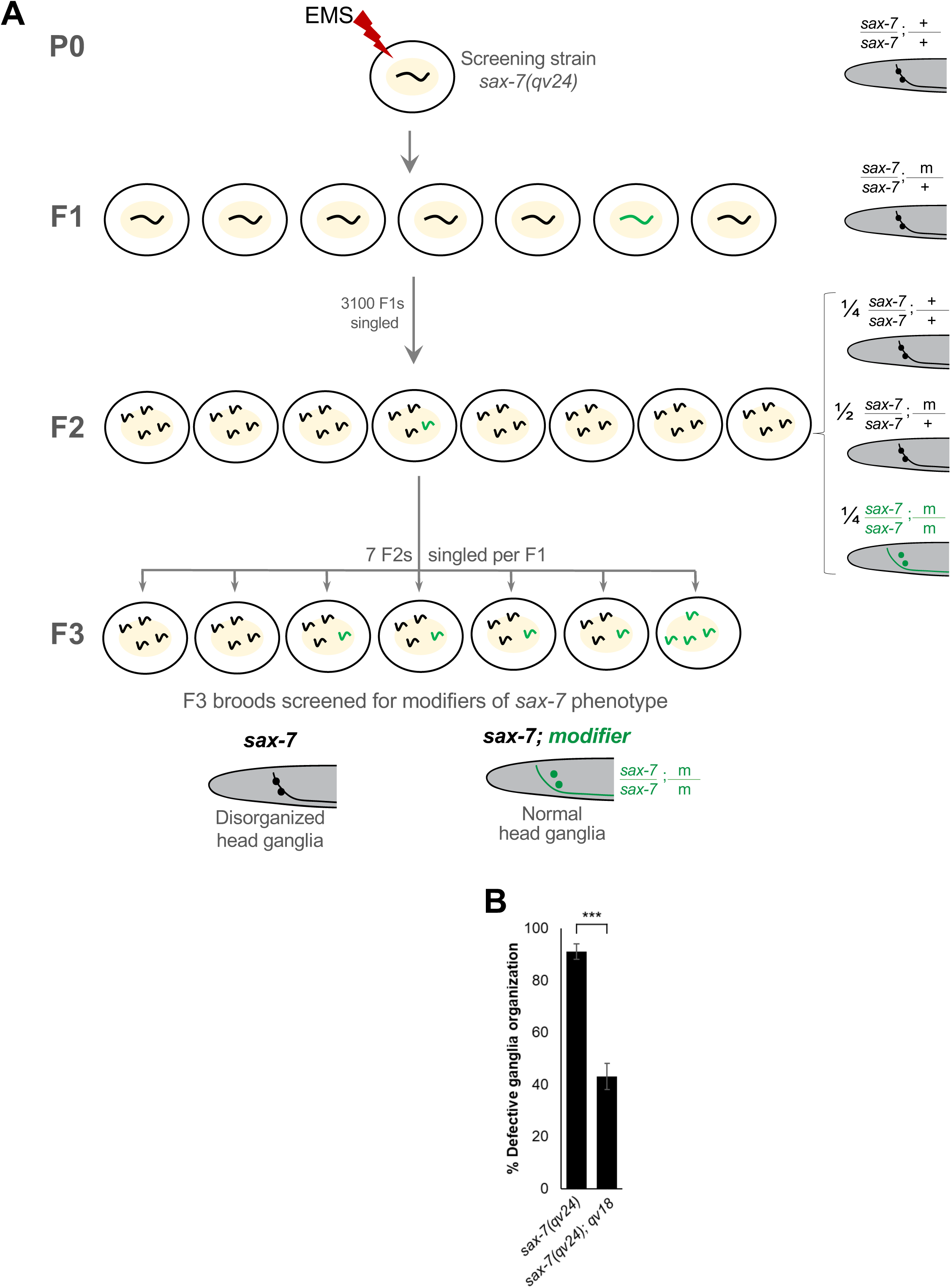
Forward F2 clonal genetic screen to identify novel neuronal maintenance factors by searching for modifiers of the defective head neuronal organization of *sax-7* mutants. (**A**) Schematic of the forward genetic screen carried out to identify novel neuronal maintenance factors. P0 *sax-7(qv24)* L4 worms were mutagenized with ethyl methanesulfonate (EMS). 3 100 F1 progeny of the mutagenized P0s were picked singly onto new plates; rare animals containing a potential modifier mutation of the *sax-7* head neuronal disorganization phenotype would be heterozygous at this generation and visible only if the mutation were dominant and highly penetrant. Thus, for each F1, 7 F2s were picked singly onto new plates. If an F2 animal was heterozygous for a modifier mutation, then one quarter of its brood (F3) would be homozygous for the modifier mutation (represented in green). If an F2 animal was homozygous for a modifier mutation, its F3 brood would be homozygous for the modifier mutation. Broods of F3 adult animals were screened by fluorescence microscopy on a stereoscope to find those where a large proportion of animals displayed a modified head ganglia organization phenotype compared to non-mutagenized *sax-7(qv24)* adults. (**B**) Quantification of chemosensory neurons ASH and ASI disorganization in 2-day-old adults of *sax-7(qv24)* and double mutant animals *sax-7(qv24); qv18*, which were less defective compared to *sax-7* mutants, indicating that the newly isolated mutation *qv18* supresses the neuronal defects of *sax-7* mutants. As a note, the molecular lesion present in allele *mig-6*(*qv33)* used in this report is identical to that of *qv18* as *qv33* was CRISPR-Cas9 generated to introduce the mutation *qv18* in a clean background. Error bars are the standard error of the proportion. Asterisks denote significant difference: \*\*\**P* ≤ 0.001 (z-test). Sample sizes and data in **Supplementary Information**.2

**Figure S2.**
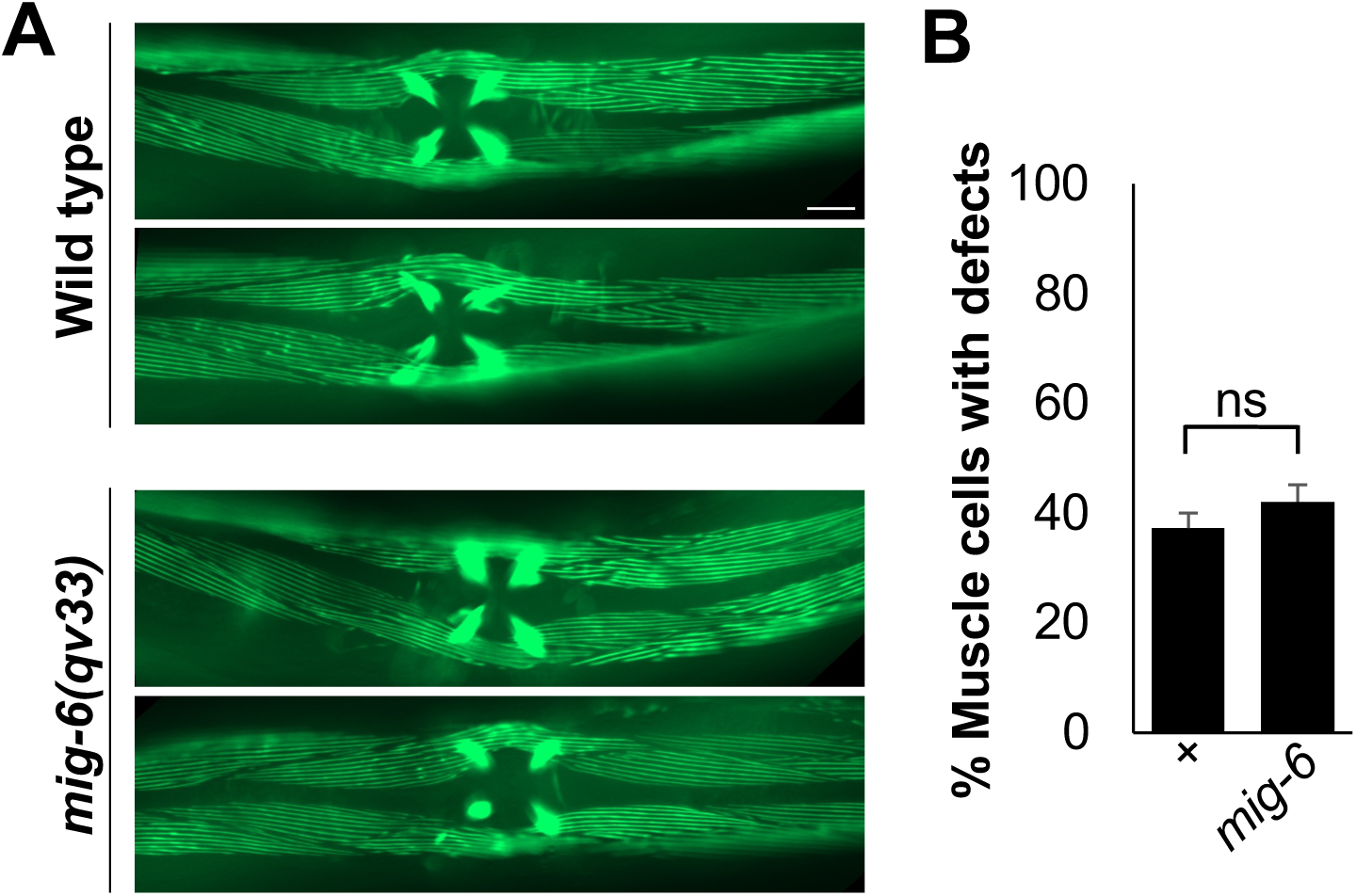
*mig-6* mutants have normal body wall muscle structure. (**A**) Images of body wall muscles in the central region of the animal’s body in wild-type and *mig-6(qv33)* mutant 1-day-old adults visualized with the *stEx30* P*myo-3*::GFP::MYO-3 reporter. Muscles of the vulva are also visible (center of images). (**B**) Quantification of body wall muscle phenotype (sarcomere organization) in wild type and *mig-6* mutants. Error bars are the standard error of the proportion. Sample sizes and data in Supplementary Information. “+” indicates wild-type strain; n.s., not significant (z-test). Scale bar, 20 µm.

**Figure S3.**
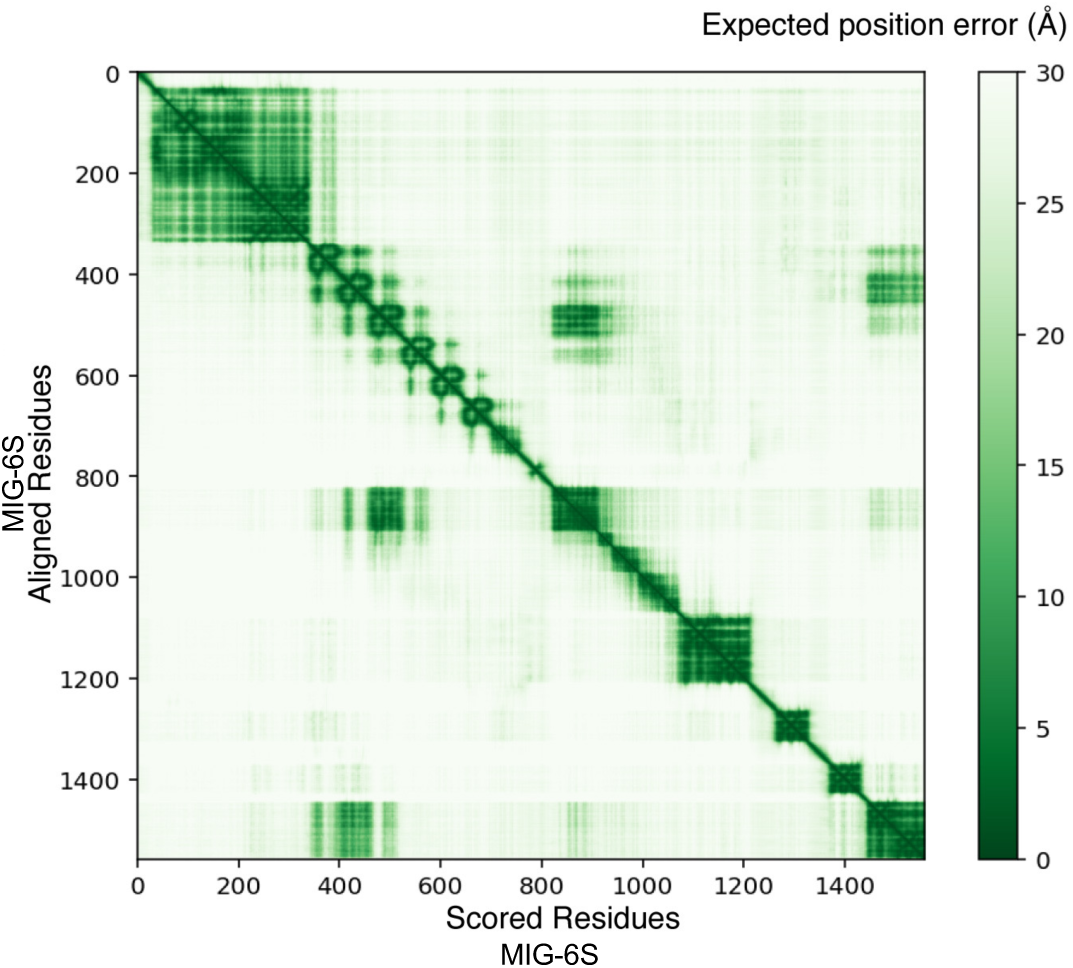
ColabFold 1.5.2-related information on MIG-6S. Predicted aligned errors (PAE) corresponding to the predicted structure of MIG-6S shown on Fig. 1D". This PAE analysis shows regions of high confidence (dark green) and low confidence (pale green) in relative domain positioning.

**Figure S4.**
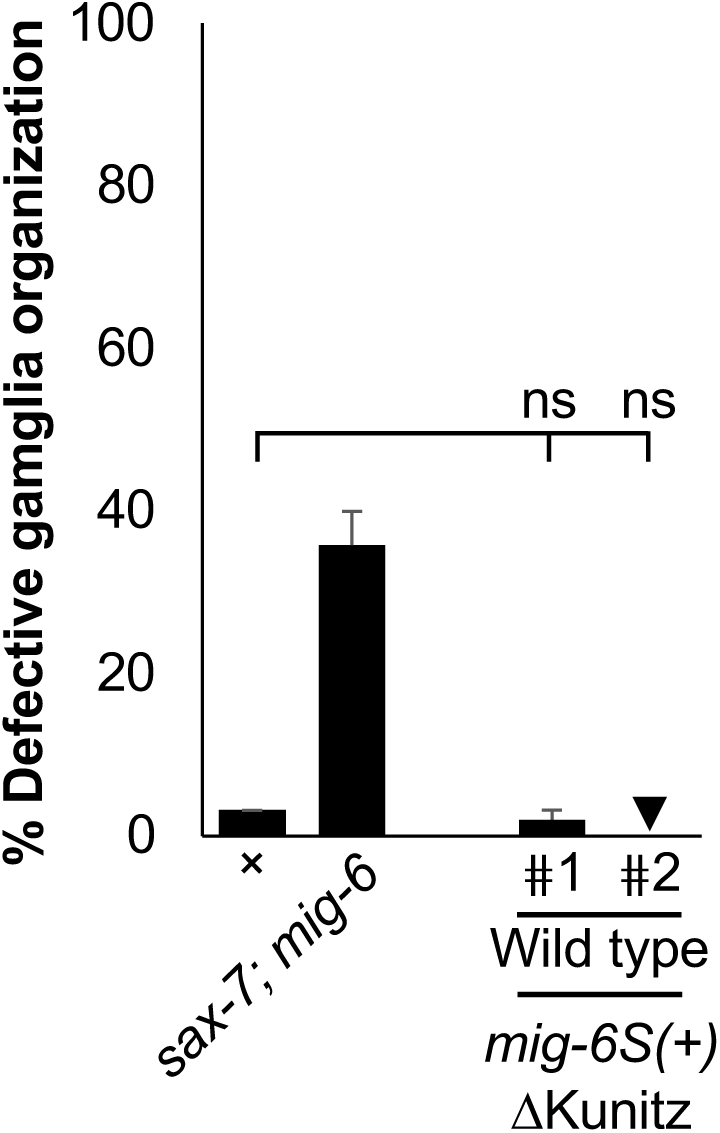
Control assays using a recombinant transgene of *mig-6S* lacking the Kunitz domains in wild-type animals. Two independent transgenic lines overexpressing *mig-6S* lacking the Kunitz domains display normal neuronal organization, indicating that overexpression of this transgene does not lead to neuronal defects. Thus, the restoration of neuronal defects in *sax-7; mig-6* double mutants expressing this transgene (Fig. 1E) is the result of this transgene’s rescuing activity. Error bars are the standard error of the proportion. Sample sizes and data in Supplementary Information. “+” indicates wild-type strain; n.s., not significant (z-test).

**Figure S5.**
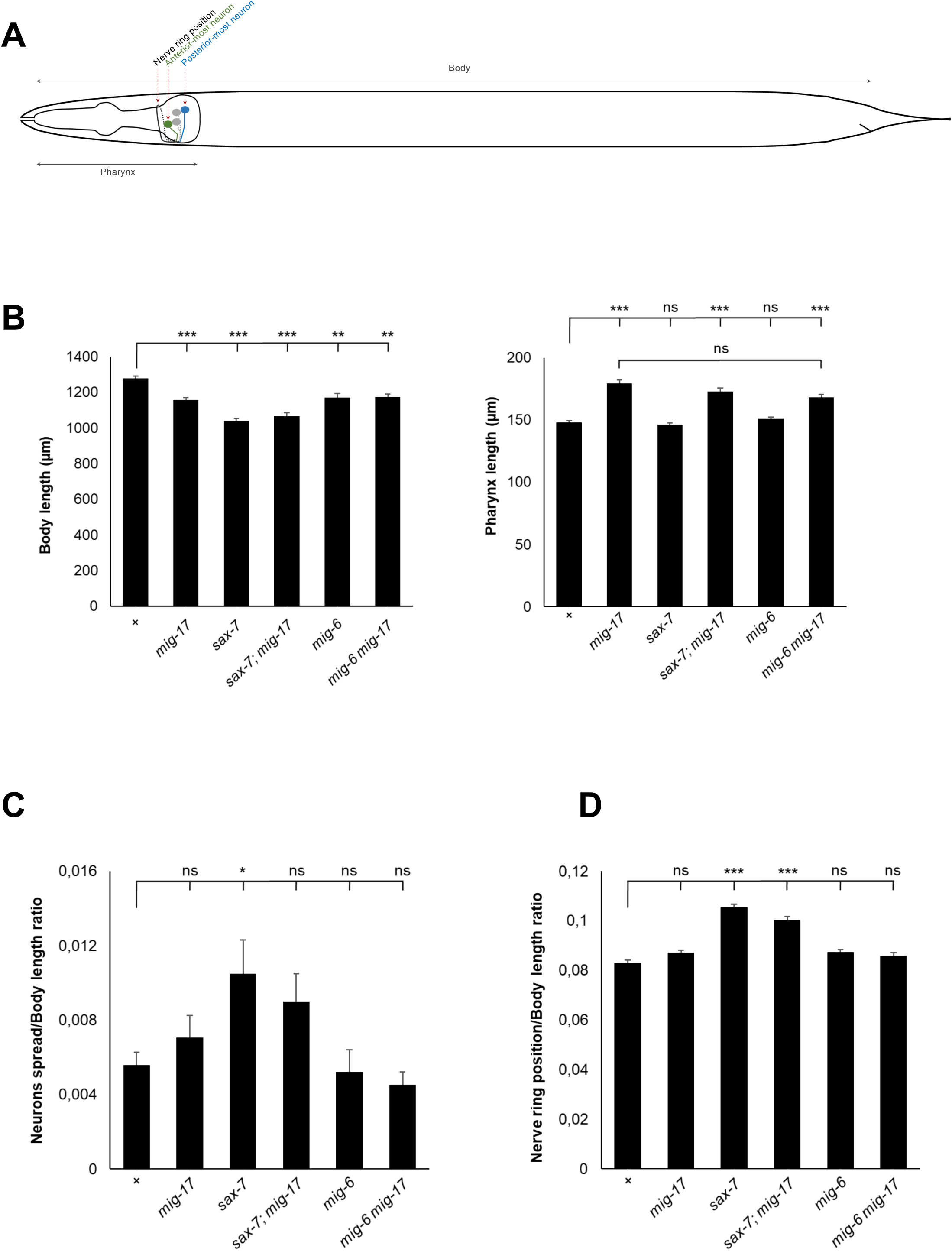
The longer pharynx of *mig-17* mutants does not affect the position of ASH and ASI soma and of the nerve ring relative to body length. We measured the distribution of neuronal soma positions and the position of the nerve ring, and calculated ratios of neuronal position to body length. (**A**) Schematic diagram of *C. elegans* hermaphrodite with four chemosensory neurons ASI and ASH drawn, left and right for each pair (as would be visualized using reporter *hdIs26* P*sra-6::DsRed2*). The most anterior soma of the four neurons, the most posterior soma of the four neurons, and the nerve ring are indicated. Body length was measured as the distance from the mouth opening to the anus, and pharynx length was measured as the distance between the mouth opening and the posterior edge of the terminal bulb. (**B**) Analysis of body and pharynx lengths of 2-day-old wild-type animals and mutants *mig-17*, *mig-6* and *sax-7*, as well as single or combined mutant animals of *mig-17* with *sax-7* or *mig-6* mutations. (**C**) Quantification of the ratio between the spread of neuronal positions (distance between the position of the anterior most neuron to the posterior most neuron in each animal) and the body length of 2-day-old animals. The higher the ratio, the more posterior the neuronal structures are. For example, because of head ganglia defects in *sax-7* mutants, the ratios appear higher in these mutants. (**D**) Quantification of the ratio between the nerve ring position and the body length in 2-day-old animals. The position of head neuronal structures, including soma and nerve ring, does not depend on pharynx length, being in a conserved position relative to body length. Thus, in *mig-17* mutants, despite their longer pharynx, there is no difference in neuronal position of their somas and nerve ring compared to the wild type. Error bars are the standard error of the mean. Asterisks denote significant difference: \**P* ≤ 0.05, \*\*\**P* ≤ 0.001 (Wilcoxon Mann-Whitney test for body length and ANOVA for pharynx length in B; ANOVA in C and D); *P*-values were corrected by multiplying by the number of comparisons, Bonferroni correction; sample sizes and data in **Supplementary Information**). “+” indicates wild-type strain; n.s., not significant.

**Figure S6.**
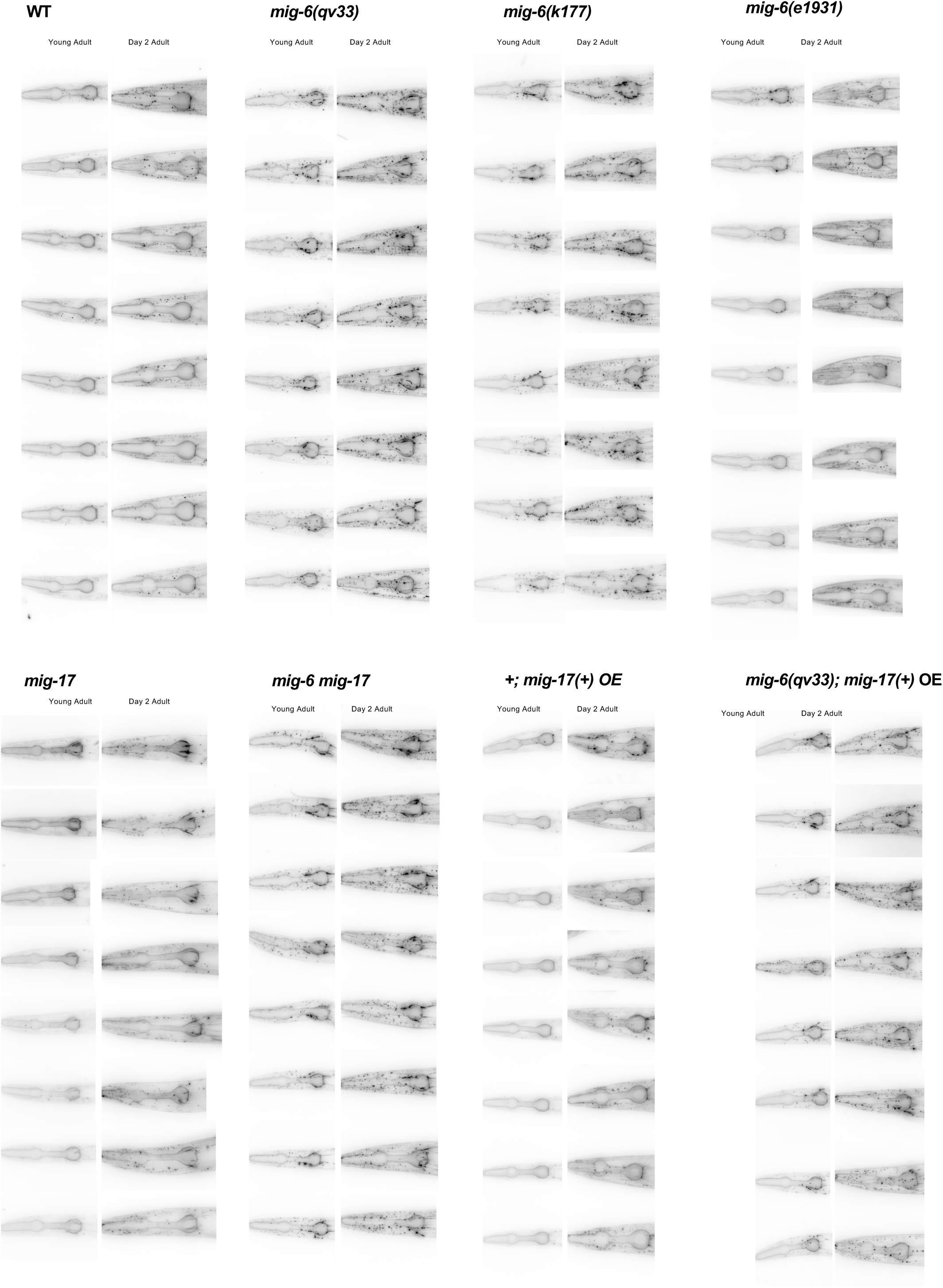
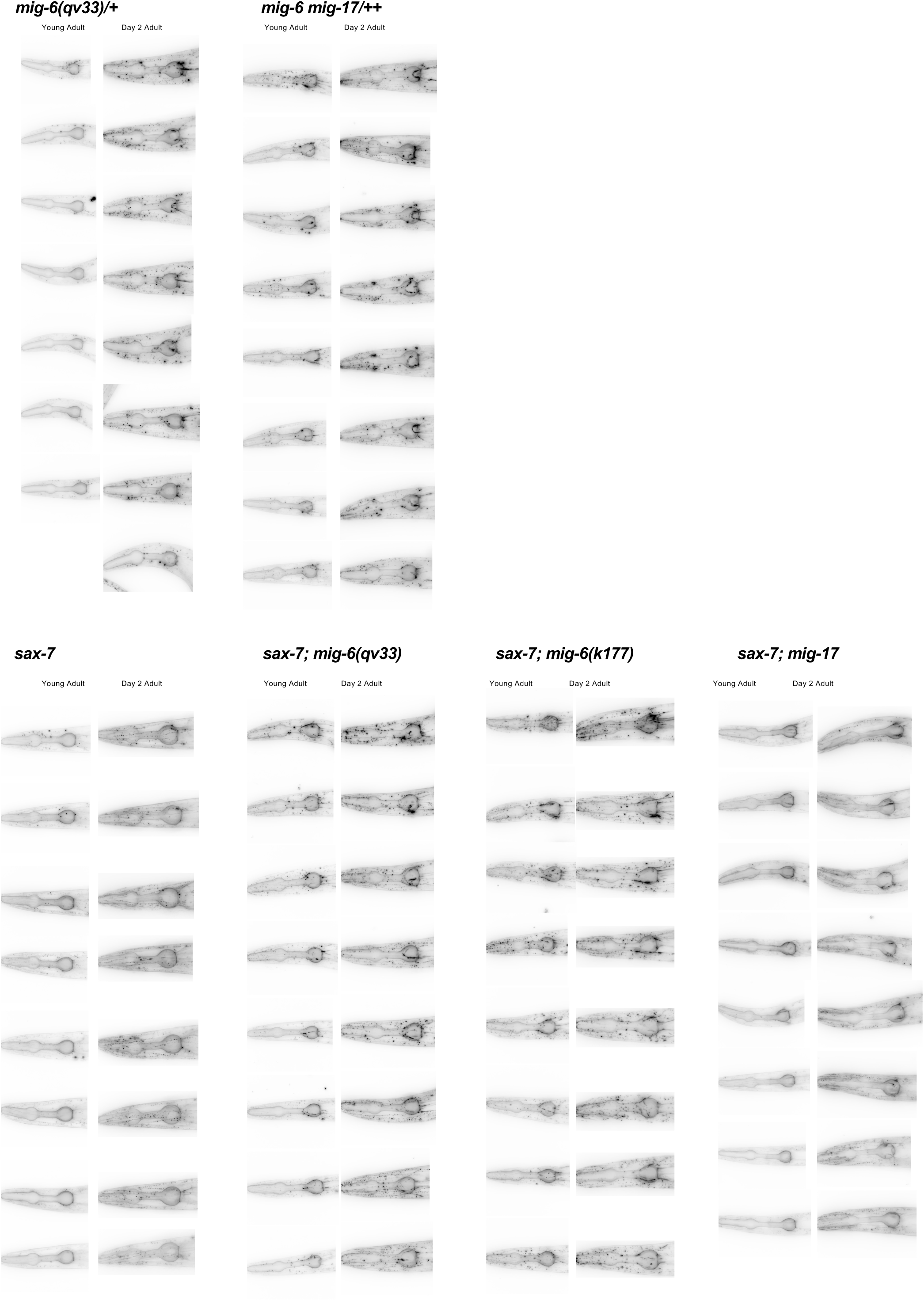
Distribution of collagen IV in different mutants. Panels of fluorescence images (sum projection) of EMB-9::mCherry (*qyIs46* P*emb-9::emb-9::mCherry*) for 2-day-old adults of genotypes analyzed in this study. The robustness of the fibrotic-like structures accumulation phenotype of *mig-6* and *mig-17* single and combined mutants can be appreciated, as well as the range of their positions in the head (coinciding mostly with the region around terminal bulb of the pharynx).

**Figure S7.**
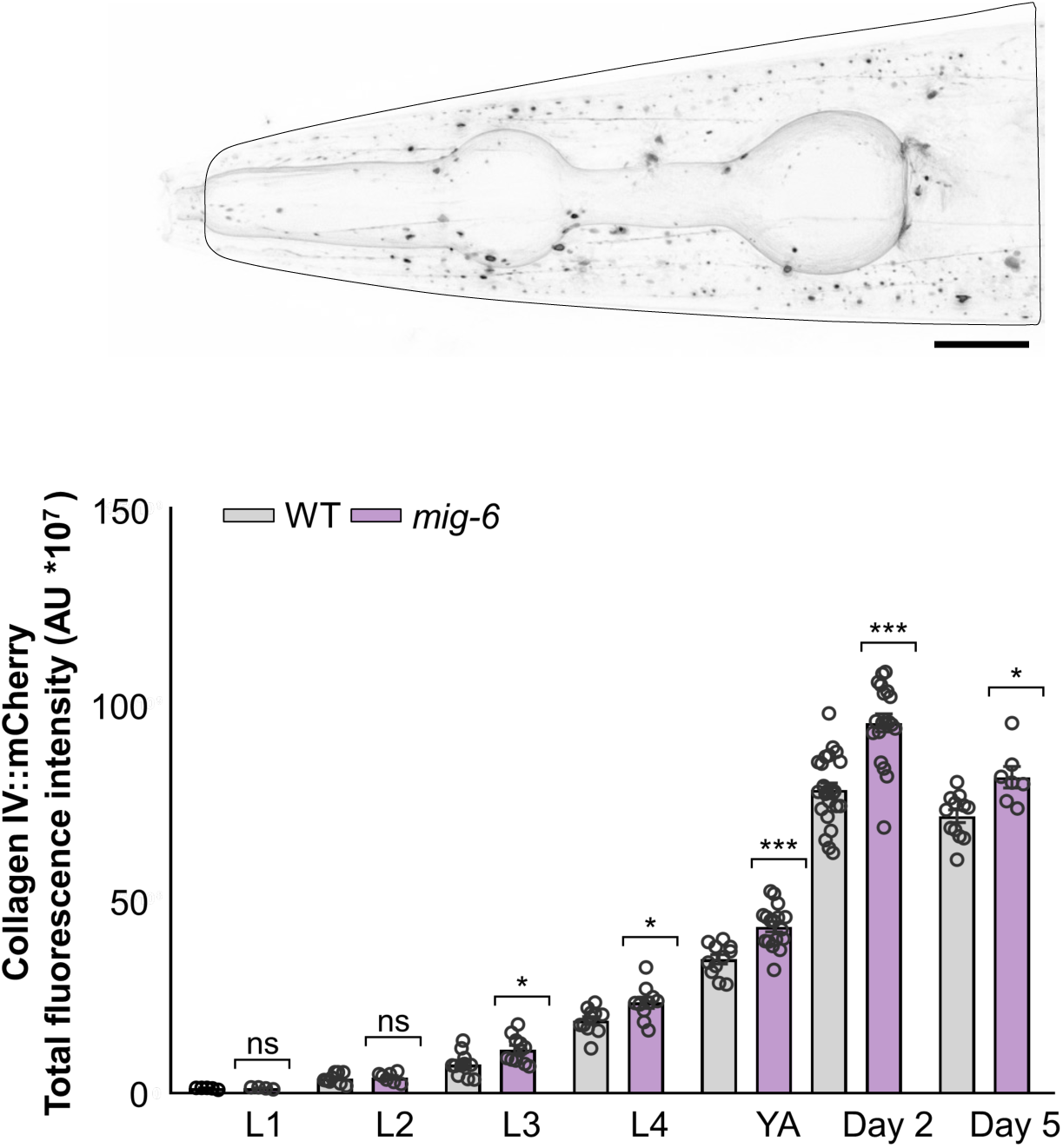
Collagen IV levels increase with age in *mig-6* mutant animals. Quantification of fluorescence intensity of EMB-9::mCherry (*qyIs46* P*emb-9::emb-9::mCherry*) in the head region at larval stages (L1, L2, L3, and L4), and adult ages (young adult, 2- and 5-day-old adult animals) in control animals and *mig-6(qv33)* mutants. ROI was drawn from the mouth opening to a position posterior to terminal bulb by 20% of the pharynx length. A.U., arbitrary units. Scale bar, 20 µm. Error bars are the standard error of the mean. Asterisks denote significant difference: \**P* ≤ 0.05, \*\*\**P* ≤ 0.001 (Wilcoxon test to compare data of the first larval stage L1 between WT and *mig-6* mutants and t-test for the other stages); *P*-values were corrected by multiplying by the number of comparisons, Bonferroni correction; sample sizes and data in **Supplementary Information**). n.s., not significant.

**Figure S8.**
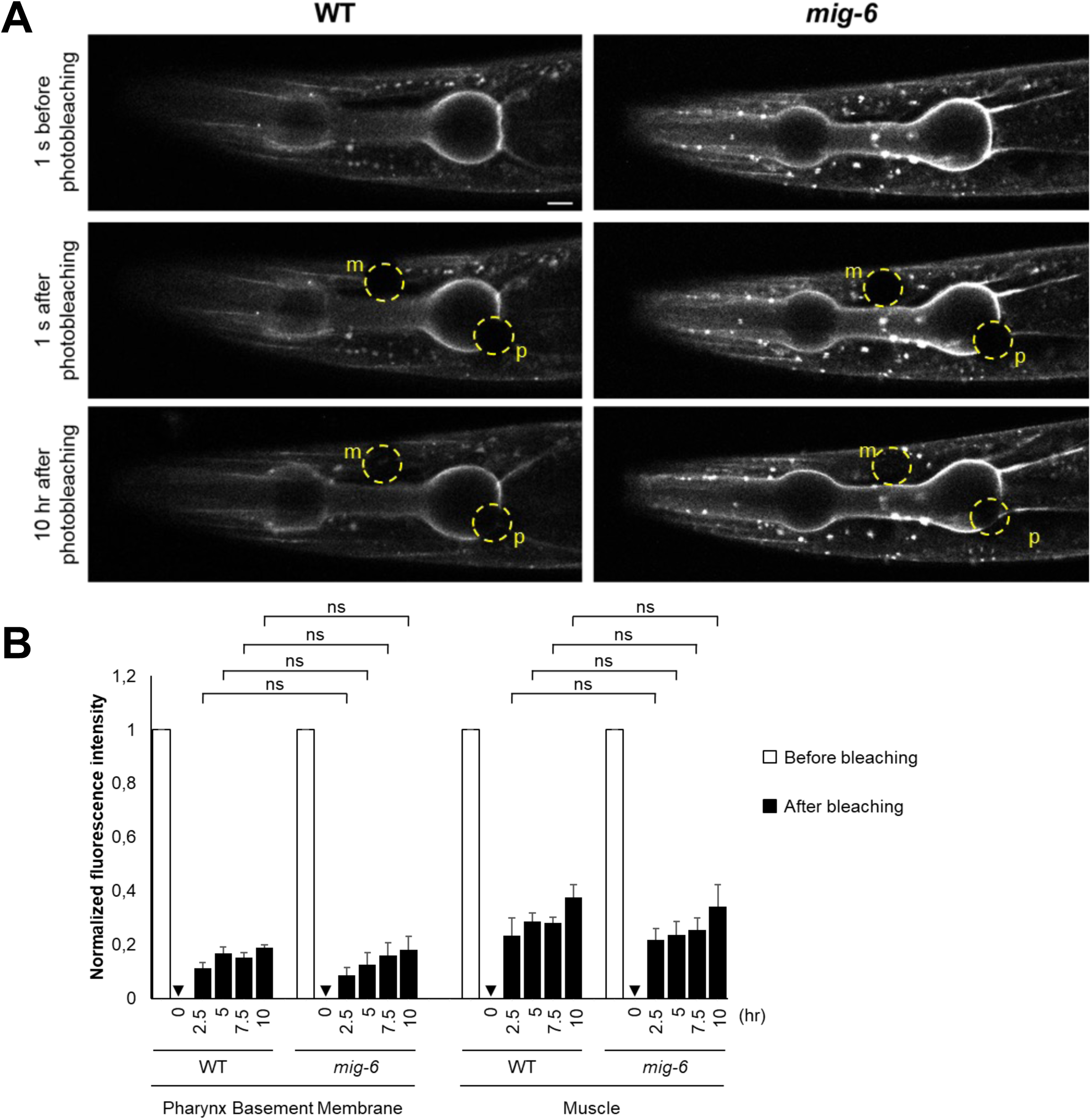
Collagen IV stability in the basement membrane is not affected by *mig-6* loss of function. We used fluorescence recovery after photobleaching (FRAP) analysis to assess EMB-9::mCherry fluorescence recovery in wild-type and *mig-6* mutants. (**A**) Images show a single confocal z slice of the head region. ROI of photobleached regions is indicated by circles (m for muscle and p for pharynx). A control region located outside of the animal is also used (not shown). (**B**) Quantification of normalized fluorescence recovery in wild-type and mutant *mig-6* young adults, in pharynx basement membrane and in muscle after 0, 2.5, 5, 7.5, and 10 hours. After photobleaching a specific region of EMB-9::mCherry in muscle and pharyngeal basement membrane, we measured FRAP and normalized the data against an unbleached control region. n.s., not significant (t-test). sample sizes and data in **Supplementary Information**).

**Figure S9.**
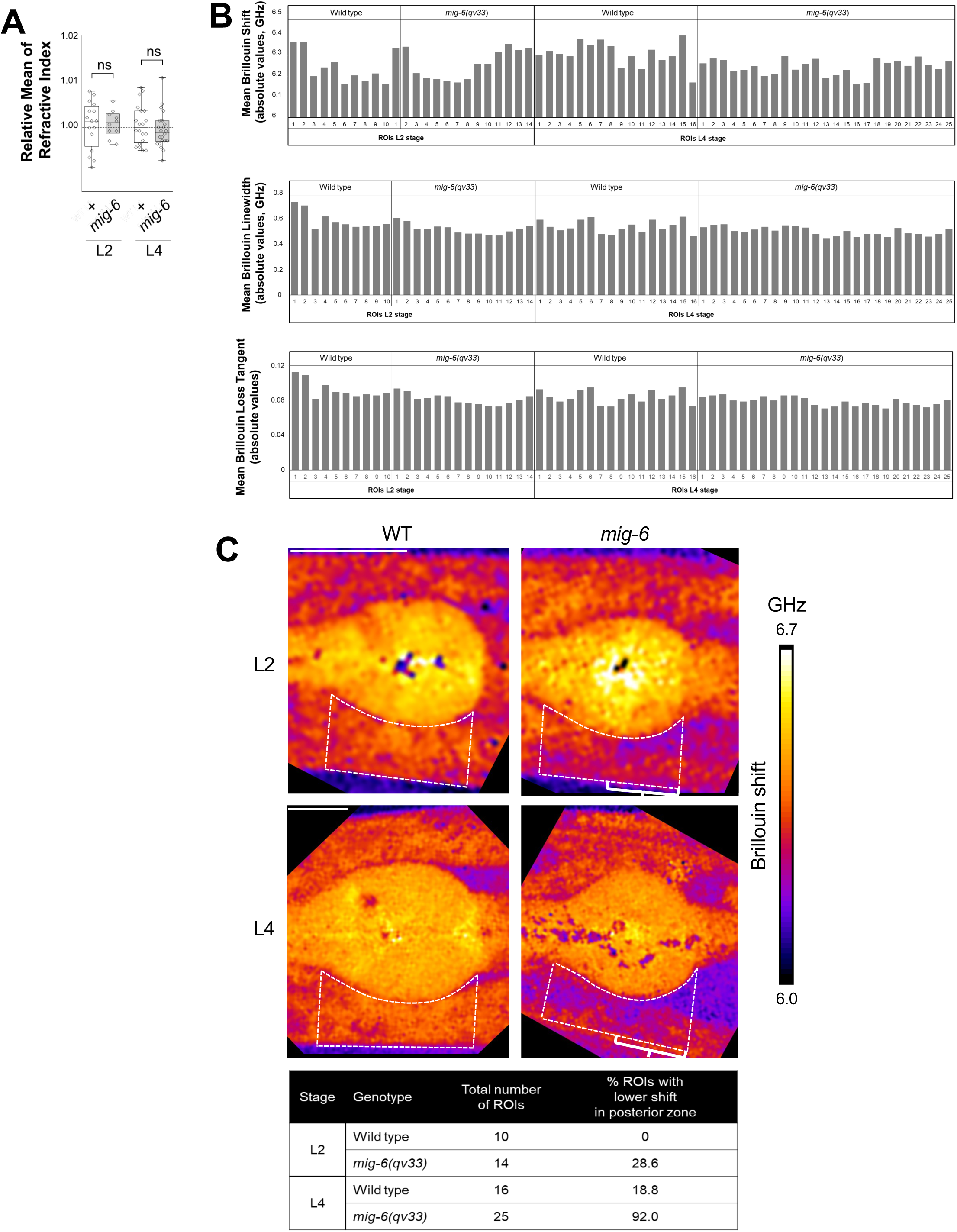
Refractive index measurements, as well as Brillouin shift, linewidth, and loss tangent quantifications, in wild-type and *mig-6* mutant animals. (A) Absolute values of the refractive index of ROIs measured in wild-type and *mig-6* mutant animals are not significantly different. Error bars are the standard error of the mean; n.s., not significant (t-test); sample sizes and data in Supplementary Information). (**B**) Quantification of Brillouin shift, linewidth, and loss tangent in each ROI in wild-type and *mig-6* mutant animals at L2 and L4 stages. (**C**) The posterior zone of the head region around the pharynx (indicated by brackets) shows less elasticity in *mig-6* mutants. Brillouin shift images of wild-type and *mig-6* mutant animals at L2 and L4 stages; scale bar, 20 µm. Table provides the quantification of the percentage of ROIs with a lower Brillouin shift in this posterior zone. Source data in **Supplementary Information**).

**Figure S10.**
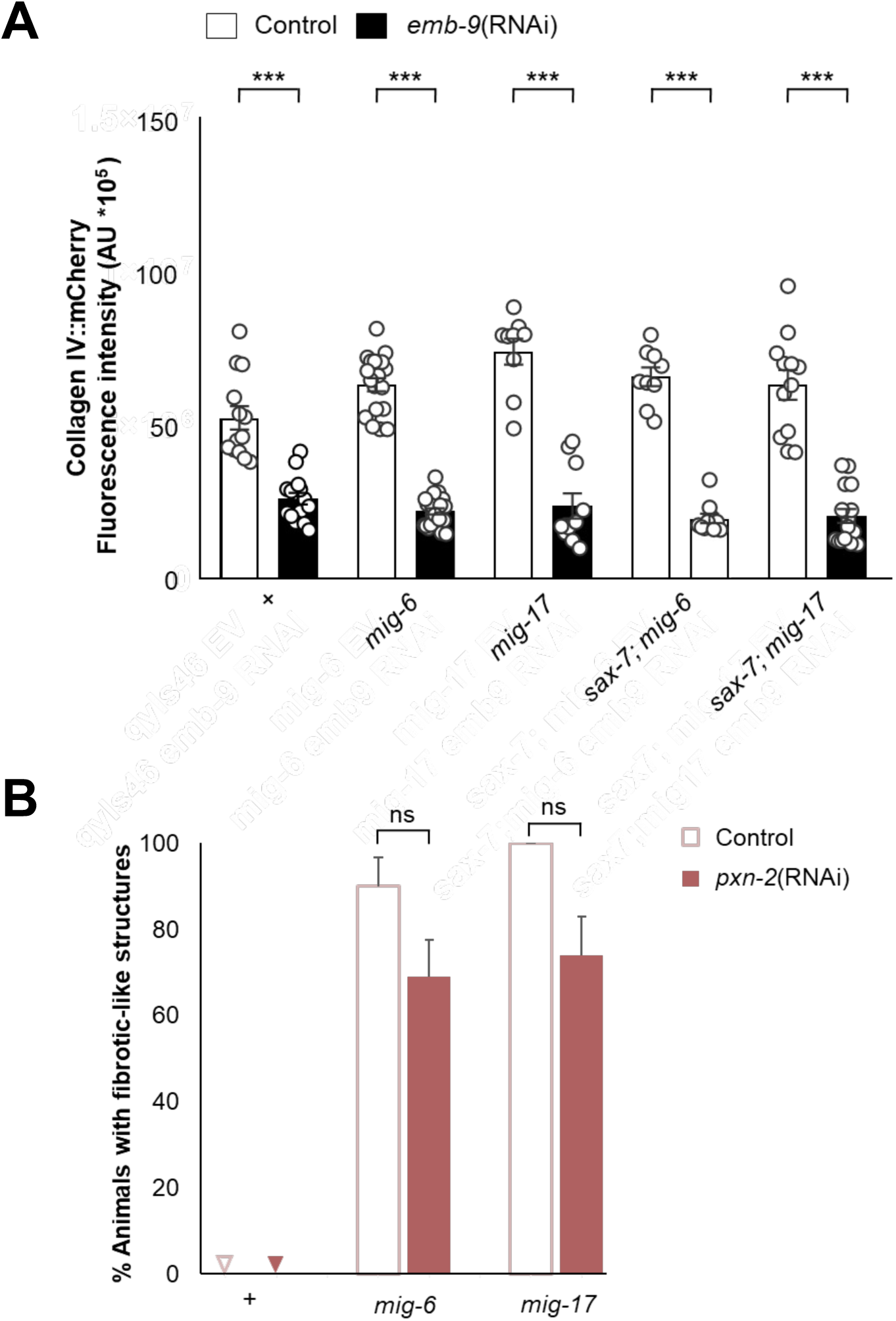
*emb-9*(RNAi) treatment decreases collagen IV levels as expected, and *pxn-2*(RNAi) treatment does not significantly affect the penetrance of the fibrotic phenotype. (**A**) Quantification of fluorescence intensity of EMB-9::mCherry in the head region in 2-day-old adult animals, which were subjected to control (empty vector) or *emb-9*(RNAi), indicating efficient collagen IV depletion. Same ROI as for **Fig. S6**. A.U., arbitrary units. Error bars are the standard error of the mean. Asterisks denote significant difference: \**P* ≤ 0.05, \*\*\**P* ≤ 0.001 (Wilcoxon Mann-Whitney test); *P*-values were corrected by multiplying by the number of comparisons, Bonferroni correction; sample sizes and data in Supplementary Information). n.s., not significant. (**B**) Quantification of percentage of 2-day-old animals displaying fibrotic-like structures in adult animals, which were subjected to control (empty vector) or *pxn-2*(RNAi). *pxn-2* depletion does not significantly decrease the percentage of animals displaying fibrotic-like structures (however, the number and continuity of these fibrotic-like structures are profoundly affected by *pxn-2*(RNAi), see Fig. 6 **E, F, G**). n.s., not significant (z-test). Sample sizes and data in **Supplementary Information**).

**Figure S11.**
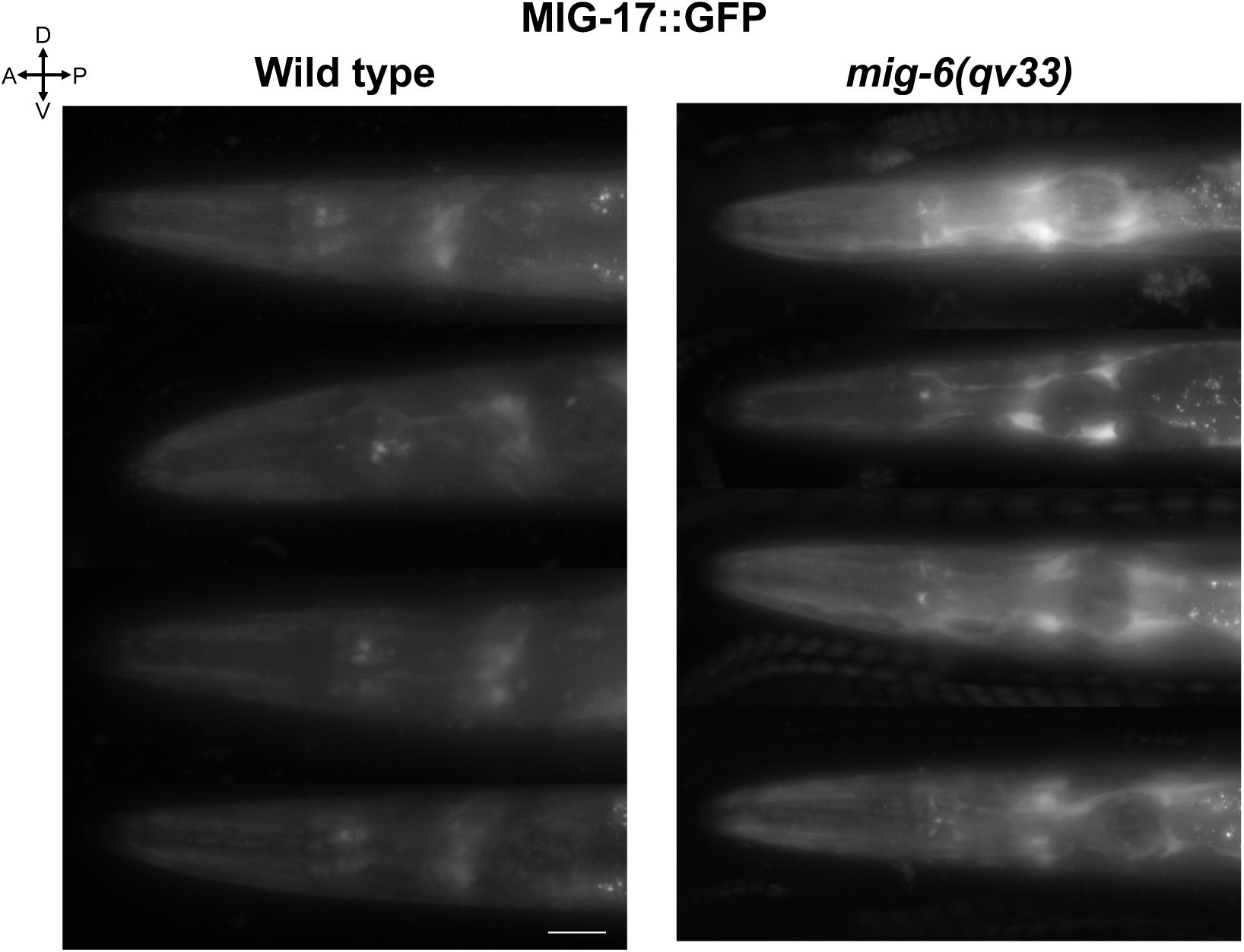
MIG-17/ADAMTS is upregulated in *mig-6* mutants. Representative fluorescence images of wild-type and mutant *mig-6(qv33)* young adults showing the level and the distribution of MIG-17::GFP using reporter *evIs213* P*mig-17::gfp*. ROI was as in Fig. 5D. Scale bar, 20 µm.

**Figure S12.**
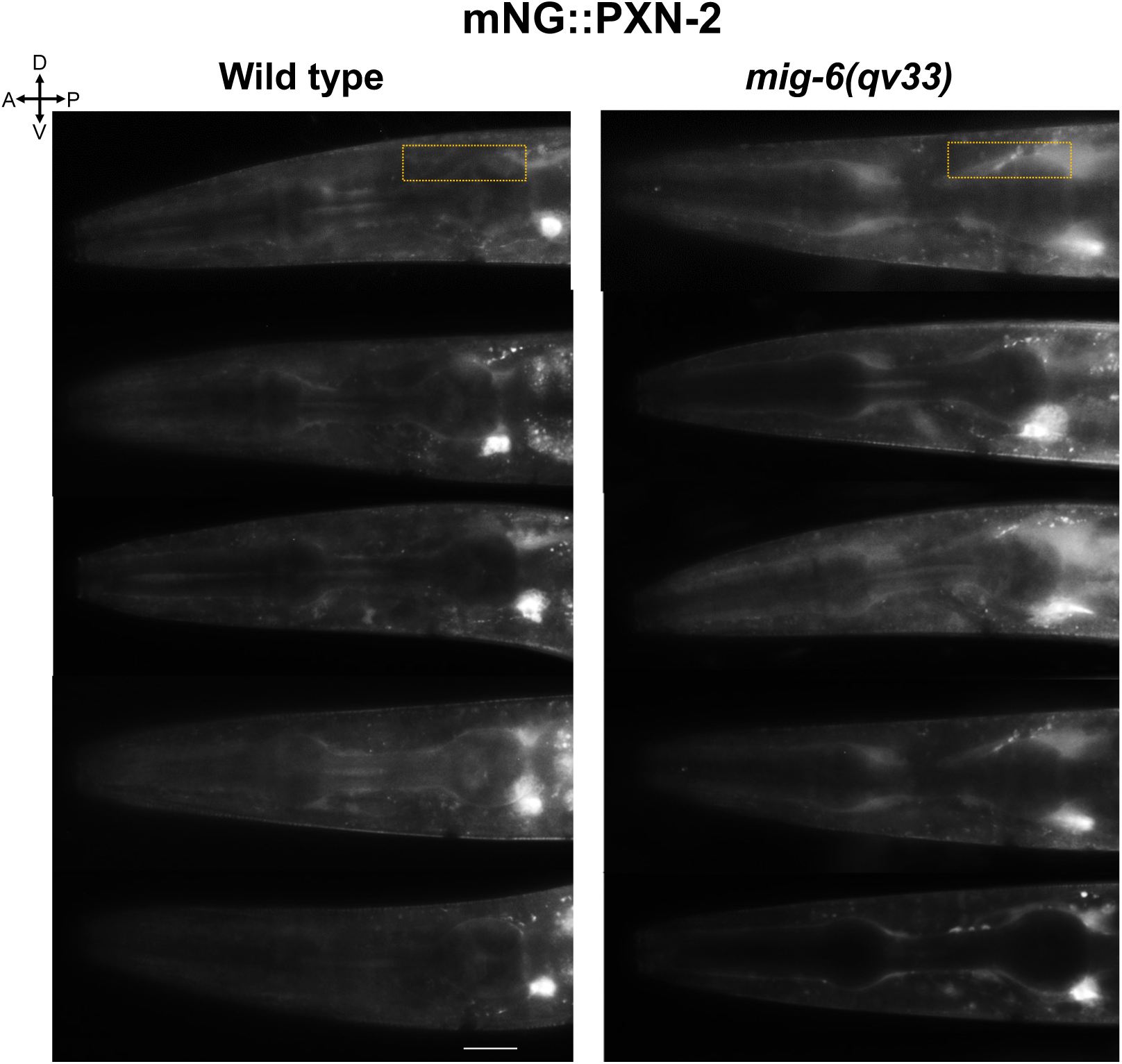
PXN-2 is upregulated in *mig-6* mutants. Representative fluorescence images of wild-type and *mig-6(qv33)* mutant young adults showing the level and distribution of mNG::PXN-2 using reporter *qy76 mNeonGreen::pxn-2 N-term*. The orange rectangle indicates the ROI used to quantify fluorescence intensity in Fig. 6D. Scale bar, 20 µm.

**S1 Table.**
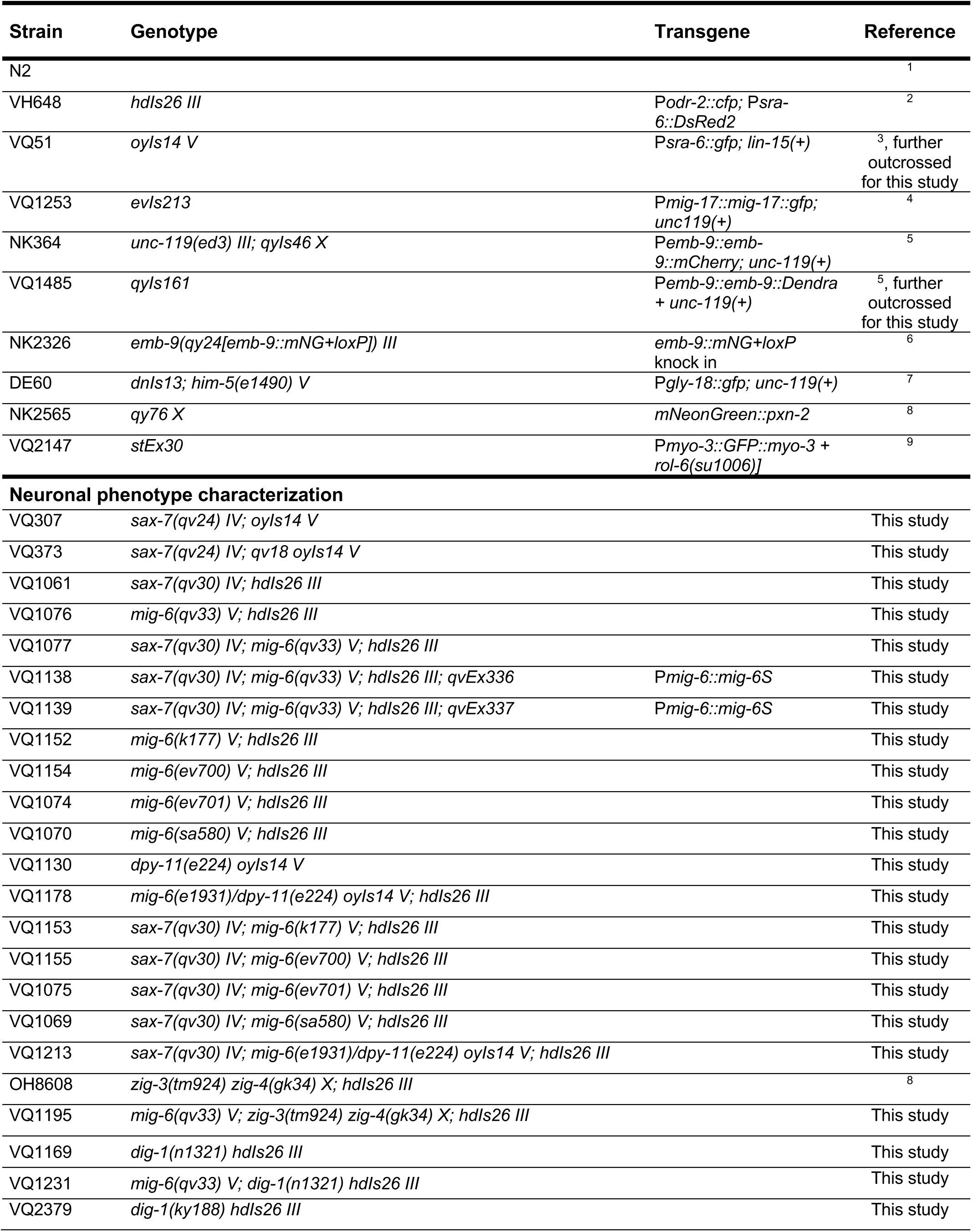
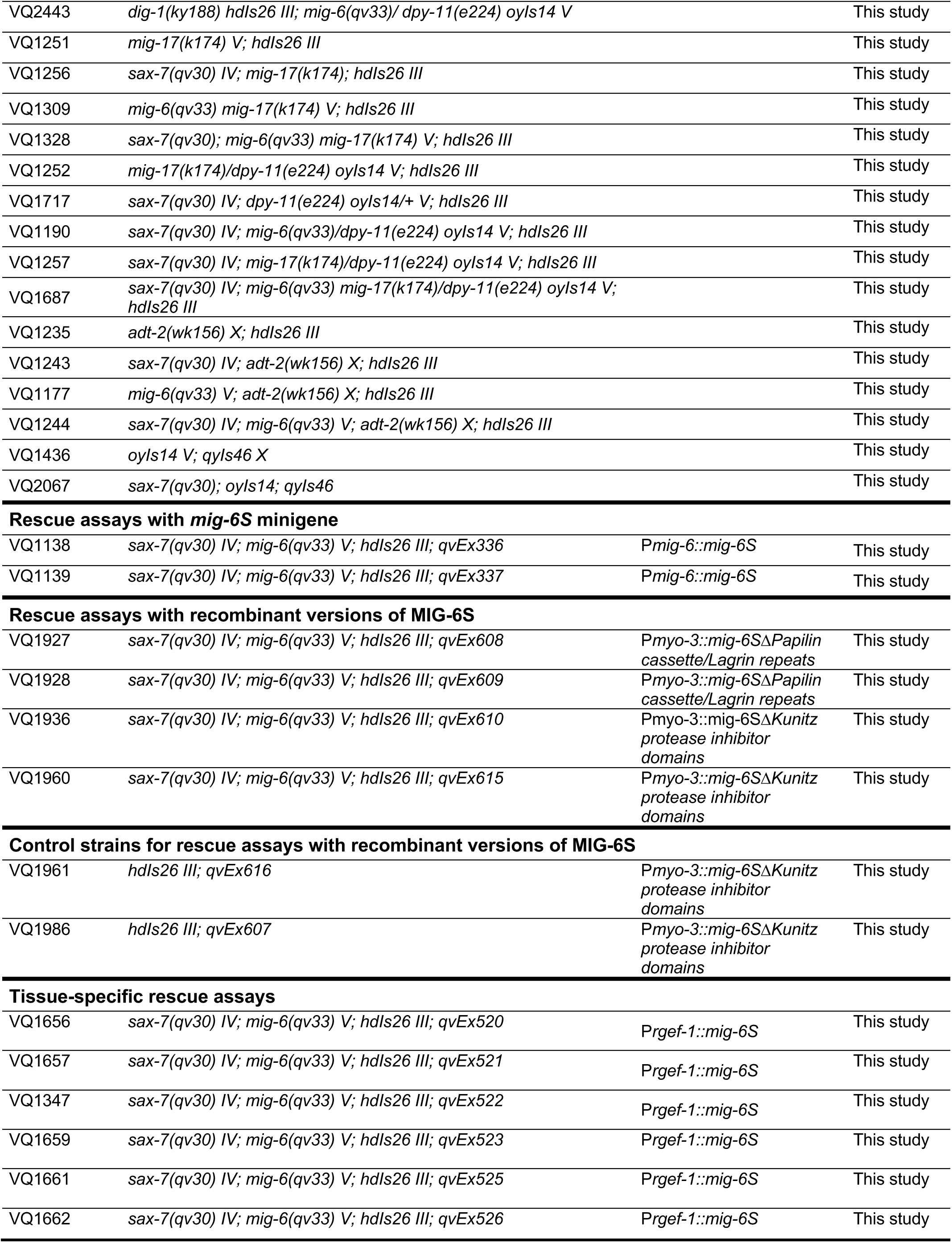
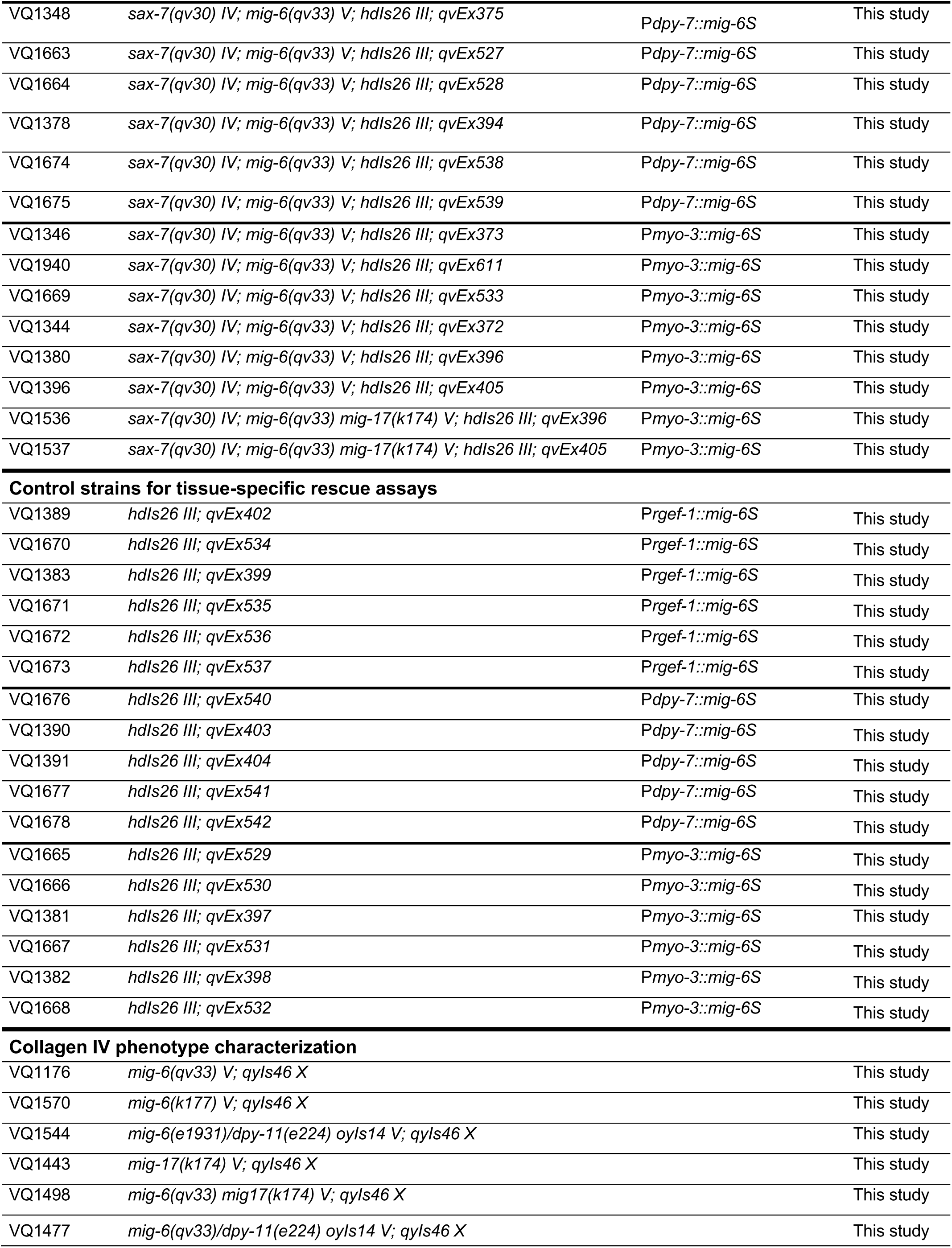
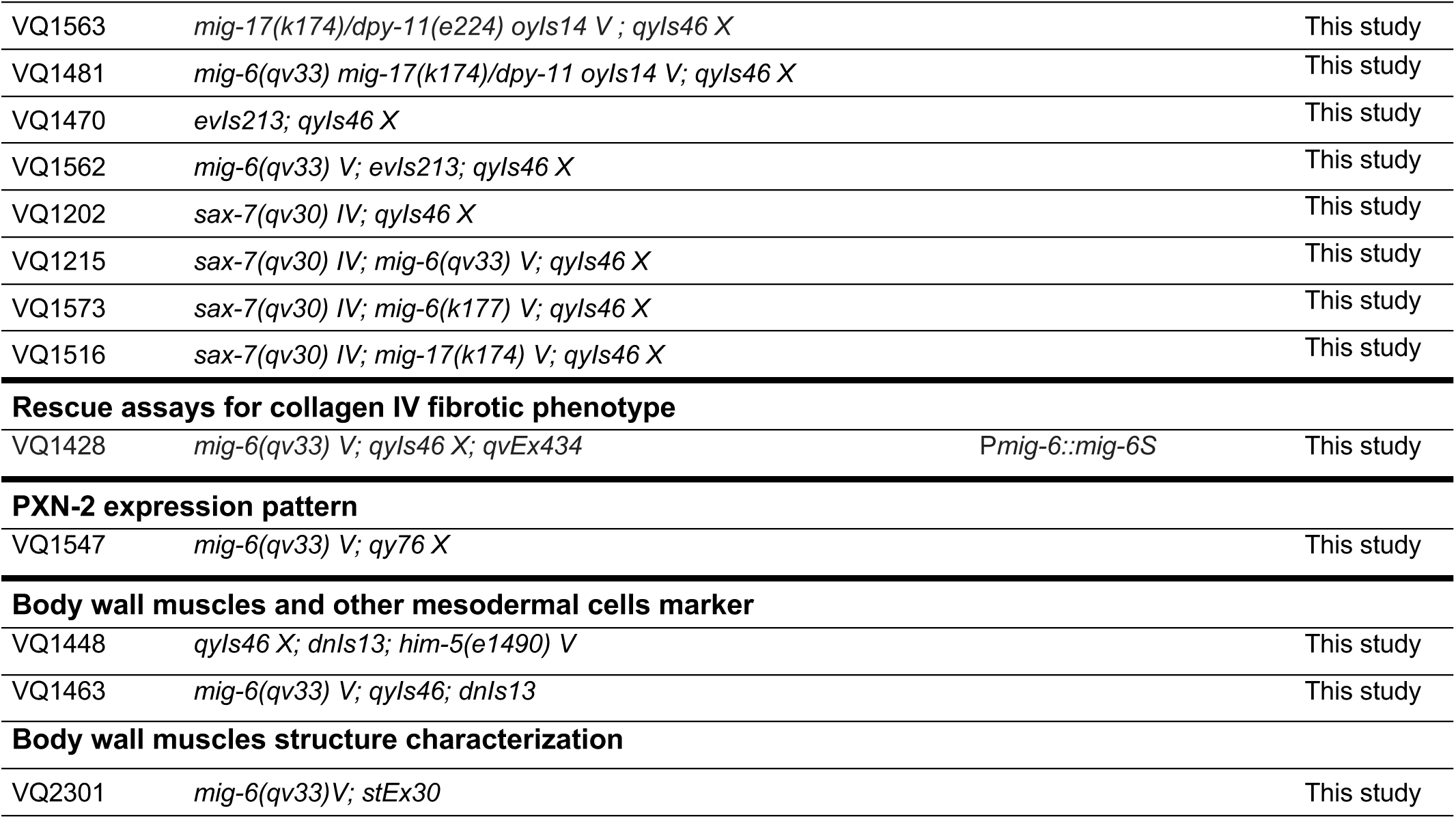
List of strains used.

**S2 Table.**
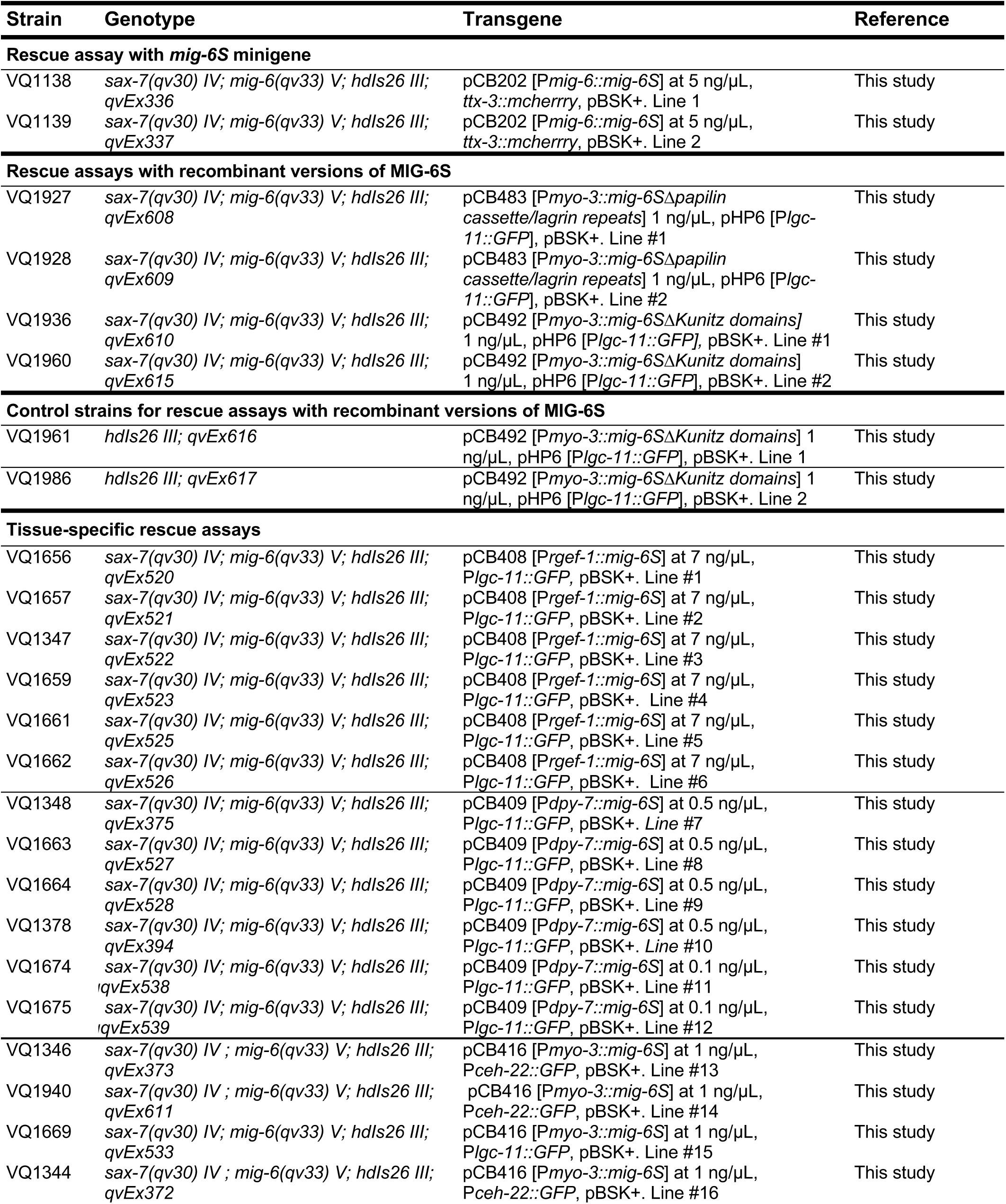
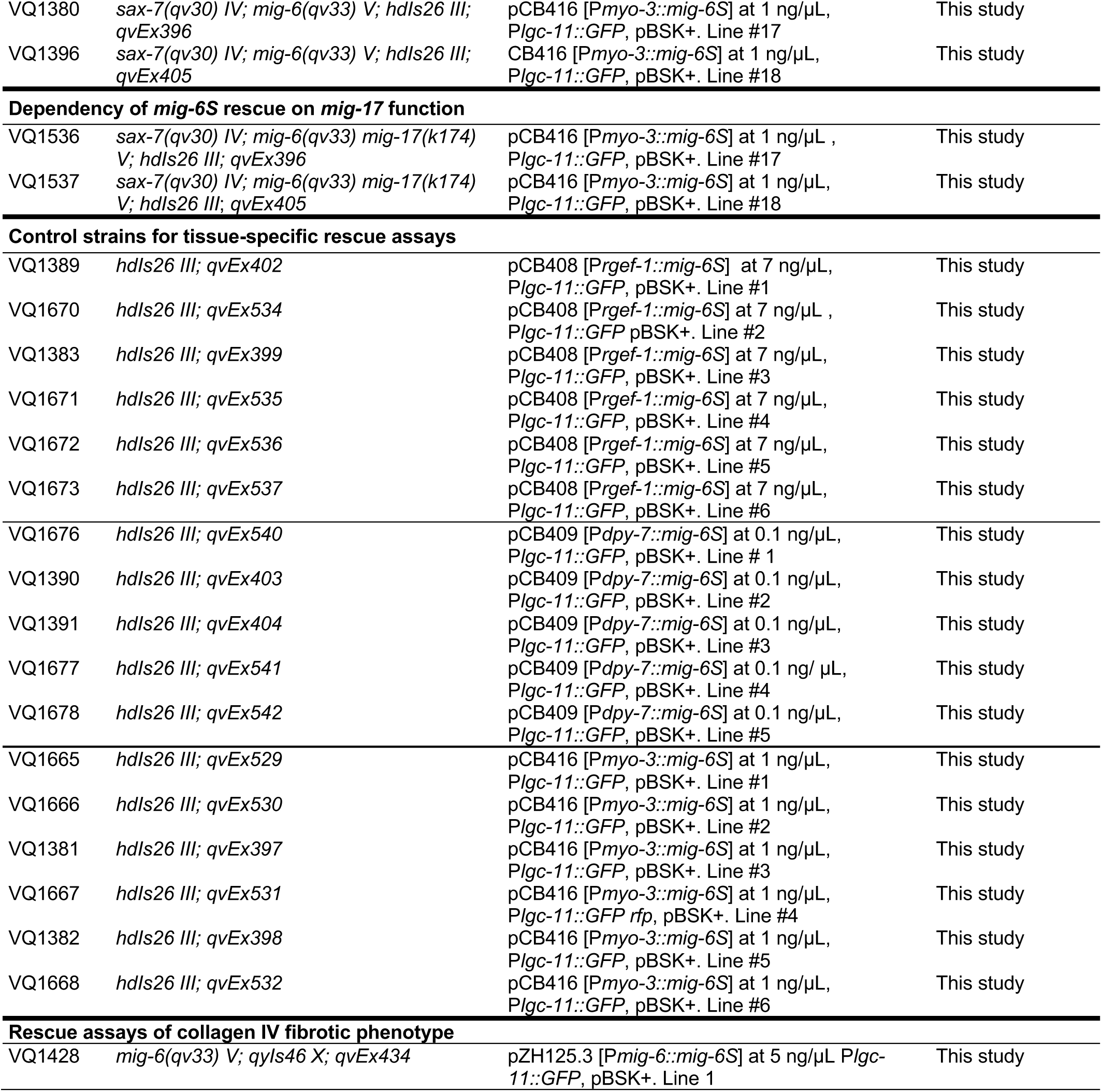
Detailed information on transgenic strains used.

| Strain | Genotype | Transgene | Reference |
| --- | --- | --- | --- |
| **Rescue assay with *mig-6S* minigene** |  |  |  |
| VQ1138 | *sax-7(qv30) IV; mig-6(qv33) V; hdlIs26 III; qvEx336* | pCB202 [*Pmig-6::mig-6S*] at 5 ng/μL, *ttx-3::mcherry*, pBSK+. Line 1 | This study |
| VQ1139 | *sax-7(qv30) IV; mig-6(qv33) V; hdlIs26 III; qvEx337* | pCB202 [*Pmig-6::mig-6S*] at 5 ng/μL, *ttx-3::mcherry*, pBSK+. Line 2 | This study |
| **Rescue assays with recombinant versions of MIG-6S** |  |  |  |
| VQ1927 | *sax-7(qv30) IV; mig-6(qv33) V; hdlIs26 III; qvEx608* | pCB483 [*Pmyo-3::mig-6SΔpapilin cassette/lagrin repeats*] 1 ng/μL, pHP6 [*Plgc-11::GFP*], pBSK+. Line #1 | This study |
| VQ1928 | *sax-7(qv30) IV; mig-6(qv33) V; hdlIs26 III; qvEx609* | pCB483 [*Pmyo-3::mig-6SΔpapilin cassette/lagrin repeats*] 1 ng/μL, pHP6 [*Plgc-11::GFP*], pBSK+. Line #2 | This study |
| VQ1936 | *sax-7(qv30) IV; mig-6(qv33) V; hdlIs26 III; qvEx610* | pCB492 [*Pmyo-3::mig-6SΔKunitz domains*] 1 ng/μL, pHP6 [*Plgc-11::GFP*], pBSK+. Line #1 | This study |
| VQ1960 | *sax-7(qv30) IV; mig-6(qv33) V; hdlIs26 III; qvEx615* | pCB492 [*Pmyo-3::mig-6SΔKunitz domains*] 1 ng/μL, pHP6 [*Plgc-11::GFP*], pBSK+. Line #2 | This study |
| **Control strains for rescue assays with recombinant versions of MIG-6S** |  |  |  |
| VQ1961 | *hdlIs26 III; qvEx616* | pCB492 [*Pmyo-3::mig-6SΔKunitz domains*] 1 ng/μL, pHP6 [*Plgc-11::GFP*], pBSK+. Line 1 | This study |
| VQ1986 | *hdlIs26 III; qvEx617* | pCB492 [*Pmyo-3::mig-6SΔKunitz domains*] 1 ng/μL, pHP6 [*Plgc-11::GFP*], pBSK+. Line 2 | This study |
| **Tissue-specific rescue assays** |  |  |  |
| VQ1656 | *sax-7(qv30) IV; mig-6(qv33) V; hdlIs26 III; qvEx520* | pCB408 [*Prgef-1::mig-6S*] at 7 ng/μL, *Plgc-11::GFP*, pBSK+. Line #1 | This study |
| VQ1657 | *sax-7(qv30) IV; mig-6(qv33) V; hdlIs26 III; qvEx521* | pCB408 [*Prgef-1::mig-6S*] at 7 ng/μL, *Plgc-11::GFP*, pBSK+. Line #2 | This study |
| VQ1347 | *sax-7(qv30) IV; mig-6(qv33) V; hdlIs26 III; qvEx522* | pCB408 [*Prgef-1::mig-6S*] at 7 ng/μL, *Plgc-11::GFP*, pBSK+. Line #3 | This study |
| VQ1659 | *sax-7(qv30) IV; mig-6(qv33) V; hdlIs26 III; qvEx523* | pCB408 [*Prgef-1::mig-6S*] at 7 ng/μL, *Plgc-11::GFP*, pBSK+. Line #4 | This study |
| VQ1661 | *sax-7(qv30) IV; mig-6(qv33) V; hdlIs26 III; qvEx525* | pCB408 [*Prgef-1::mig-6S*] at 7 ng/μL, *Plgc-11::GFP*, pBSK+. Line #5 | This study |
| VQ1662 | *sax-7(qv30) IV; mig-6(qv33) V; hdlIs26 III; qvEx526* | pCB408 [*Prgef-1::mig-6S*] at 7 ng/μL, *Plgc-11::GFP*, pBSK+. Line #6 | This study |
| VQ1348 | *sax-7(qv30) IV; mig-6(qv33) V; hdlIs26 III; qvEx375* | pCB409 [*Pdpy-7::mig-6S*] at 0.5 ng/μL, *Plgc-11::GFP*, pBSK+. Line #7 | This study |
| VQ1663 | *sax-7(qv30) IV; mig-6(qv33) V; hdlIs26 III; qvEx527* | pCB409 [*Pdpy-7::mig-6S*] at 0.5 ng/μL, *Plgc-11::GFP*, pBSK+. Line #8 | This study |
| VQ1664 | *sax-7(qv30) IV; mig-6(qv33) V; hdlIs26 III; qvEx528* | pCB409 [*Pdpy-7::mig-6S*] at 0.5 ng/μL, *Plgc-11::GFP*, pBSK+. Line #9 | This study |
| VQ1378 | *sax-7(qv30) IV; mig-6(qv33) V; hdlIs26 III; qvEx394* | pCB409 [*Pdpy-7::mig-6S*] at 0.5 ng/μL, *Plgc-11::GFP*, pBSK+. Line #10 | This study |
| VQ1674 | *sax-7(qv30) IV; mig-6(qv33) V; hdlIs26 III; qvEx538* | pCB409 [*Pdpy-7::mig-6S*] at 0.1 ng/μL, *Plgc-11::GFP*, pBSK+. Line #11 | This study |
| VQ1675 | *sax-7(qv30) IV; mig-6(qv33) V; hdlIs26 III; qvEx539* | pCB409 [*Pdpy-7::mig-6S*] at 0.1 ng/μL, *Plgc-11::GFP*, pBSK+. Line #12 | This study |
| VQ1346 | *sax-7(qv30) IV; mig-6(qv33) V; hdlIs26 III; qvEx373* | pCB416 [*Pmyo-3::mig-6S*] at 1 ng/μL, *Pceh-22::GFP*, pBSK+. Line #13 | This study |
| VQ1940 | *sax-7(qv30) IV; mig-6(qv33) V; hdlIs26 III; qvEx611* | pCB416 [*Pmyo-3::mig-6S*] at 1 ng/μL, *Pceh-22::GFP*, pBSK+. Line #14 | This study |
| VQ1669 | *sax-7(qv30) IV; mig-6(qv33) V; hdlIs26 III; qvEx533* | pCB416 [*Pmyo-3::mig-6S*] at 1 ng/μL, *Plgc-11::GFP*, pBSK+. Line #15 | This study |
| VQ1344 | *sax-7(qv30) IV; mig-6(qv33) V; hdlIs26 III; qvEx372* | pCB416 [*Pmyo-3::mig-6S*] at 1 ng/μL, *Pceh-22::GFP*, pBSK+. Line #16 | This study |
| VQ1380 | *sax-7(qv30) IV; mig-6(qv33) V; hdlIs26 III; qvEx396* | pCB416 [Pmyo-3::mig-6S] at 1 ng/μL, Plgc-11::GFP, pBSK+. Line #17 | This study |
| VQ1396 | *sax-7(qv30) IV; mig-6(qv33) V; hdlIs26 III; qvEx405* | CB416 [Pmyo-3::mig-6S] at 1 ng/μL, Plgc-11::GFP, pBSK+. Line #18 | This study |
| Dependency of mig-6S rescue on mig-17 function |  |  |  |
| VQ1536 | *sax-7(qv30) IV; mig-6(qv33) mig-17(k174) V; hdlIs26 III; qvEx396* | pCB416 [Pmyo-3::mig-6S] at 1 ng/μL, Plgc-11::GFP, pBSK+. Line #17 | This study |
| VQ1537 | *sax-7(qv30) IV; mig-6(qv33) mig-17(k174) V; hdlIs26 III; qvEx405* | pCB416 [Pmyo-3::mig-6S] at 1 ng/μL, Plgc-11::GFP, pBSK+. Line #18 | This study |
| Control strains for tissue-specific rescue assays |  |  |  |
| VQ1389 | *hdlIs26 III; qvEx402* | pCB408 [Prgef-1::mig-6S] at 7 ng/μL, Plgc-11::GFP, pBSK+. Line #1 | This study |
| VQ1670 | *hdlIs26 III; qvEx534* | pCB408 [Prgef-1::mig-6S] at 7 ng/μL, Plgc-11::GFP pBSK+. Line #2 | This study |
| VQ1383 | *hdlIs26 III; qvEx399* | pCB408 [Prgef-1::mig-6S] at 7 ng/μL, Plgc-11::GFP, pBSK+. Line #3 | This study |
| VQ1671 | *hdlIs26 III; qvEx535* | pCB408 [Prgef-1::mig-6S] at 7 ng/μL, Plgc-11::GFP, pBSK+. Line #4 | This study |
| VQ1672 | *hdlIs26 III; qvEx536* | pCB408 [Prgef-1::mig-6S] at 7 ng/μL, Plgc-11::GFP, pBSK+. Line #5 | This study |
| VQ1673 | *hdlIs26 III; qvEx537* | pCB408 [Prgef-1::mig-6S] at 7 ng/μL, Plgc-11::GFP, pBSK+. Line #6 | This study |
| VQ1676 | *hdlIs26 III; qvEx540* | pCB409 [Pdpy-7::mig-6S] at 0.1 ng/μL, Plgc-11::GFP, pBSK+. Line #1 | This study |
| VQ1390 | *hdlIs26 III; qvEx403* | pCB409 [Pdpy-7::mig-6S] at 0.1 ng/μL, Plgc-11::GFP, pBSK+. Line #2 | This study |
| VQ1391 | *hdlIs26 III; qvEx404* | pCB409 [Pdpy-7::mig-6S] at 0.1 ng/μL, Plgc-11::GFP, pBSK+. Line #3 | This study |
| VQ1677 | *hdlIs26 III; qvEx541* | pCB409 [Pdpy-7::mig-6S] at 0.1 ng/μL, Plgc-11::GFP, pBSK+. Line #4 | This study |
| VQ1678 | *hdlIs26 III; qvEx542* | pCB409 [Pdpy-7::mig-6S] at 0.1 ng/μL, Plgc-11::GFP, pBSK+. Line #5 | This study |
| VQ1665 | *hdlIs26 III; qvEx529* | pCB416 [Pmyo-3::mig-6S] at 1 ng/μL, Plgc-11::GFP, pBSK+. Line #1 | This study |
| VQ1666 | *hdlIs26 III; qvEx530* | pCB416 [Pmyo-3::mig-6S] at 1 ng/μL, Plgc-11::GFP, pBSK+. Line #2 | This study |
| VQ1381 | *hdlIs26 III; qvEx397* | pCB416 [Pmyo-3::mig-6S] at 1 ng/μL, Plgc-11::GFP, pBSK+. Line #3 | This study |
| VQ1667 | *hdlIs26 III; qvEx531* | pCB416 [Pmyo-3::mig-6S] at 1 ng/μL, Plgc-11::GFP rfp, pBSK+. Line #4 | This study |
| VQ1382 | *hdlIs26 III; qvEx398* | pCB416 [Pmyo-3::mig-6S] at 1 ng/μL, Plgc-11::GFP, pBSK+. Line #5 | This study |
| VQ1668 | *hdlIs26 III; qvEx532* | pCB416 [Pmyo-3::mig-6S] at 1 ng/μL, Plgc-11::GFP, pBSK+. Line #6 | This study |
| Rescue assays of collagen IV fibrotic phenotype |  |  |  |
| VQ1428 | *mig-6(qv33) V; qyls46 X; qvEx434* | pZH125.3 [Pmig-6::mig-6S] at 5 ng/μL Plgc-11::GFP, pBSK+. Line 1 | This study |

**S3 Table.**
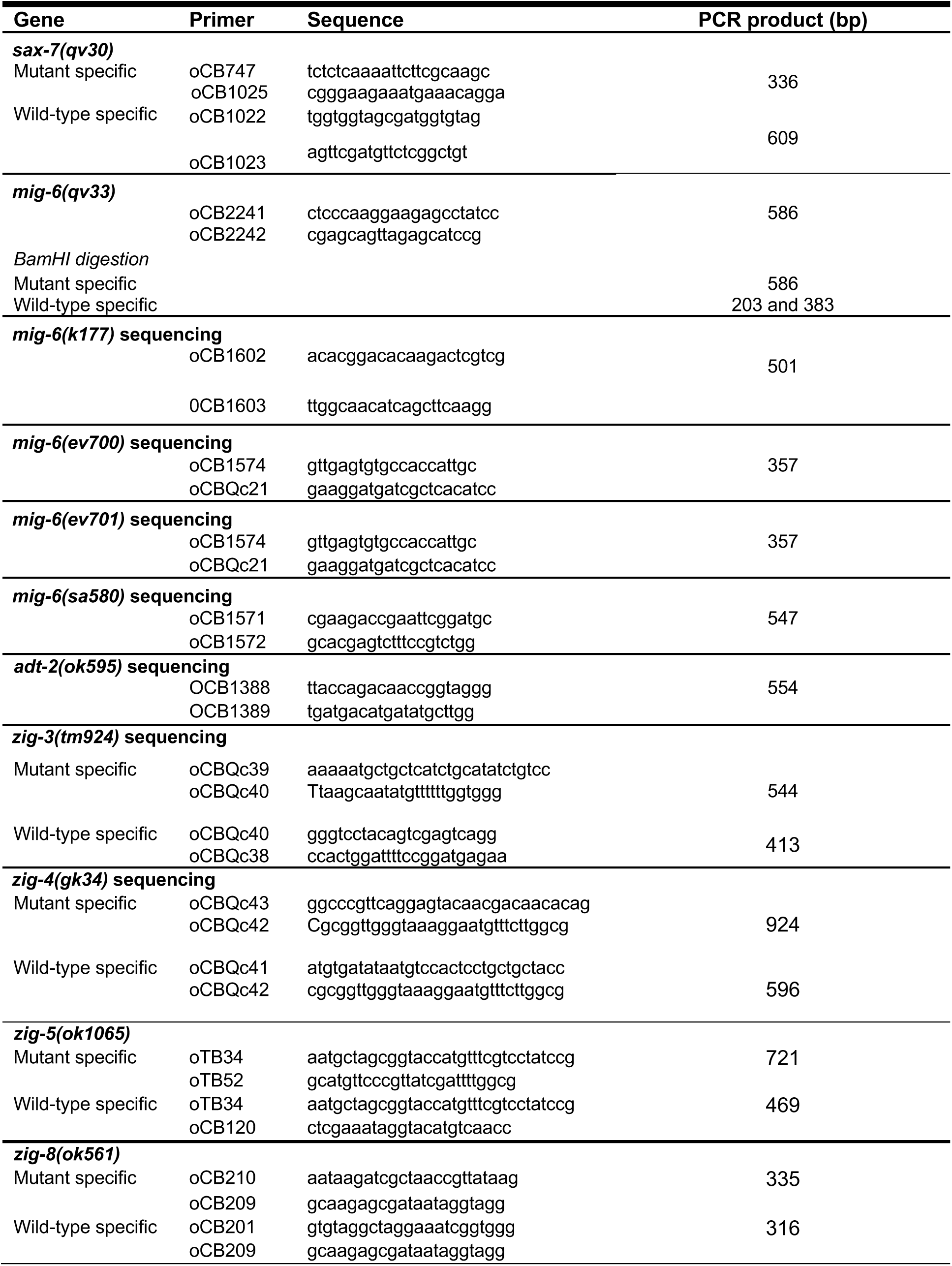
List of primers used.

## Description of Additional Supplementary Files

**File Name: Supplementary Movie 1**

**Description: *mig-6(qv33)* mutant animals display increased collagen IV and fibrotic-like structures phenotype.** Projection of fluorescence images of the head region in a 2-day-old adult *mig-6(qv33)* mutant animal expressing collagen IV reporter EMB-9::mCherry (*qyIs46)*. Collagen IV can be seen as (i) a layer surrounding the spherical terminal bulb of the pharynx, (ii) a ring at the level of the pharyngo-intestinal valve, (iii) numerous and intense small circular accumulations are inside muscle cells, (iv) prominent elongated fibrotic-like structures around the general area of the terminal bulb of the pharynx, and (v) faint lines along muscle cells.

**File Name: Supplementary Movie 2**

**Description: *mig-6(qv33)* mutant animals display increased collagen IV and fibrotic-like structures phenotype.** Projection of fluorescence images of the region of the terminal bulb of the pharynx in the head of a 2-day-old adult *mig-6(qv33)* mutant animal expressing collagen IV reporter EMB-9::mCherry (*qyIs46)*. Collagen IV can be seen as (i) a layer surrounding the spherical terminal bulb of the pharynx, (ii) a ring at the level of the pharyngo-intestinal valve, (iii) numerous and intense small circular accumulations are inside muscle cells, and (iv) prominent elongated fibrotic-like structures around the general area of the terminal bulb of the pharynx.

**File Name: Supplementary Movie 3**

**Description: Normal pattern of collagen IV in the wild type.** Projection of fluorescence images of the region of the terminal bulb of the pharynx in the head of a 2-day-old adult wild-type animal expressing collagen IV reporter EMB-9::mCherry (*qyIs46)*. Collagen IV can be seen as (i) a layer surrounding the spherical terminal bulb of the pharynx, (ii) a ring at the level of the pharyngo-intestinal valve, and (iii) small circular accumulations are inside muscle cells.

